# Identification of antimicrobial resistance-associated genes through whole genome sequencing of high and low MIC isolates of *Mycoplasma bovis*

**DOI:** 10.1101/2020.06.30.180182

**Authors:** Lisa Ledger, Jason Eidt, Hugh Y. Cai

**Affiliations:** Animal Health Lab, University of Guelph

**Keywords:** *Mycoplasma bovis*, antimicrobial resistance, whole genome sequencing, MIC, cgMLST

## Abstract

Antimicrobial resistance (AMR) in *Mycoplasma bovis* has previously been associated with mutations within topoisomerases and ribosomal genes rather than specific resistance-conferring genes. This study used whole genome sequencing (WGS) to identify potential new AMR mechanisms. It was found that a 2019 clinical isolate with high MIC (2019-043682) had a new core genome multilocus sequencing (cgMLST) type (ST10-like) and 91% homology with the published genome of *M. bovis* PG45. Closely related to PG45, a 1982 isolate (1982-M6152) shared the same cgMLST type (ST17), 97.2% homology and similar low MIC results. Known and potential AMR-associated genetic events were identified through comparison with the published *M. bovis* PG45 genome. Isolate 2019-043682 had 507 genes with non-synonymous mutations (NSMs) and 67 genes disrupted. Isolate 1982-M6152 had 81 NSMs and 20 disruptions. Based on functional roles and known mechanisms of antimicrobials, a 55 gene subset was assessed for potential AMR mechanisms. Of these, 14 were previously identified from other bacteria as sites of AMR mutation, 41 shared similar functions to them, and 11 contained gene-disrupting mutations. This study indicated that *M. bovis* may obtain high AMR characteristics by mutating or disrupting other functional genes, in addition to topoisomerases and ribosomal genes.

## 1. Introduction

*Mycoplasma bovis* is a member of the Mollicutes; bacteria which are bound in a trilayered membrane rather than a cell wall, which precludes the use of many common antimicrobial agents such as the β-lactams [1]. In cattle, *M. bovis* is a causative agent of pneumonia, arthritis, otitis media, and reproductive disease and is a contributor to the bovine respiratory disease (BRD) complex, also known as ‘shipping fever’, which is a major source of morbidity, mortality and financial loss in calf and feedlot operations. Additionally, *M. bovis* is capable of persisting for the life of a colonized animal, which may remain asymptomatic while acting as a source of infection for herdmates or offspring [1,2].

Of note, many of the antibiotics to which *M. bovis* shows resistance do not have a label claim for usage in treating *M. bovis* infections [3] but may be used in the treatment of other bovine bacterial pathogens. Given the asymptomatic nature of many *M. bovis* infections, and the high rates of colonization when animals are co-mingled (potentially over 90%) [4,5], conditions are favourable for the development of multi-drug resistant strains. With global rates of antimicrobial resistance increasing, understanding the molecular mechanisms underlying antimicrobial resistance, particularly for multi-drug resistant (MDR) strains, is critical for determining effective treatment [6], or potentially to design treatment protocols that use evolutionary approaches to counter or reverse antimicrobial resistance [7].

Unlike other members of the BRD complex such as *Mannheimia haemolytica*, *Histophilus somni* or *Pasturella multocida*, *M. bovis* is not known to possess defined antimicrobial resistance genes [8] but appears to have the molecular mechanisms of its resistance rooted in point mutations within several ribosomal and topoisomerase genes. Previous studies have used whole-genome sequencing (WGS) paired with minimum inhibitory concentration (MIC) testing to establish that mutations within *gyrA* and *parC* are linked to increased resistance to fluoroquinolones, that increased resistance to spectinomycin and the tetracyclines is linked to *rrs1-rrs2* mutations, and that *rrl1-rrl2* mutations are linked to increased resistance to florfenicol, lincosamides, macrolides and pleuromutilins [9], with *rrl3* also implicated in macrolide resistance [10].

In a large-scale MIC study of *M. bovis* strains isolated between 1978 to 2009, fluctuations in antimicrobial susceptibility over time were observed [3] with the MIC50 values, and thus resistance, increasing for several tetracycline and macrolide-class antimicrobial drugs over the span of the study. Additionally, associations between MIC50 values were observed for different antimicrobials; although sequencing these isolates fell beyond the scope of the study, the association between lincosamides, pleuromutilins and florfenicol in terms of *rrl1-rrl2* mutations seen in Sulyok *et al.* was mirrored by a similar observed association in MIC50 values with these historical samples.

A high MIC M. bovis was isolated from lung tissues of a two week old male Holstein calf submitted to the Animal Health Lab in July of 2019 for post-mortem examination. In the interest of determining possible genetic factors for this high level of resistance, whole-genome sequencing using the Illumina MiSeq platform was conducted, in tandem with WGS of a historical isolate of *M. bovis* (1982-M6152) previously categorized as low MIC for most antimicrobials [3].

## 2. Results

### 2.1. MIC testing

Three isolates of *M. bovis* were tested in triplicate, and the results of MIC testing (Table 1) were identical within all triplicates. Relative to *M. bovis* PG45, isolate 1982-M6152 shows a two-fold increase in MIC for oxytetracycline, but is otherwise identical in response to other antimicrobial compounds. Isolate 2019-043682 shows increased MICs for multiple fluoroquinolones, macrolides and tetracyclines, as well as a lincosamide, a pleuromutilin and two inhibitors of protein synthesis (gentamicin and florfenicol). For aminoglycosides the results are mixed, with increased MIC observed for spectinomycin, but no change in MIC for neomycin. All three isolates have high MICs for sulfonamides, although they retain a low MIC for combination Trimethoprim/Sulfa.

**Table 1:** Average results of MIC testing for three isolates of Mycoplasma bovis, by μg of antimicrobial compound required to inhibit growth. Isolates were tested in triplicate with identical results within each triplicate group for all antimicrobials tested.

| Antimicrobial | PG45 | 2019-043682 | 1982-M6152 |
| --- | --- | --- | --- |
| Neomycin | >32 | >32 | >32 |
| Spectinomycin | <8 | 16 | <8 |
| Trimethoprim/Sulfa | >2/38 | >2/38 | >2/38 |
| Danofloxacin | 0.5 | >1 | 0.5 |
| Enrofloxacin | 0.5 | 2 | 0.5 |
| Clindamycin | <0.25 | >16 | <0.25 |
| Tilmicosin | <4 | >64 | <4 |
| Tulathromycin | 8 | >64 | 8 |
| Tylosin Tartrate | 1 | >32 | 1 |
| Tiamulin | 1 | 8 | 1 |
| Gentamicin | 8 | 16 | 8 |
| Florfenicol | 4 | >8 | 4 |
| Sulphadimethoxine | >256 | >256 | >256 |
| Chlortetracycline | <0.5 | >8 | <0.5 |
| Oxytetracycline | <0.5 | >8 | 1 |
| Ceftiofur | >8 | >8 | >8 |

### 2.2. Whole-Genome Sequencing

Raw sequencing yield for the isolates was 167.8 Mb for 1982-M6152, and 54.02 Mb for 2019-043682. Assembled using Geneious 11 (Biomatters, Auckland, New Zealand), genome sizes were 980,739 bp for 1982-M6152 and 972,371 bp for 2019-043682. Given a sequence length of 2×251 and the documented genome size of 1.003 Mb for *M. bovis* PG45 (GenBank accession NC_014760.1), raw sequencing coverage, as defined by the equation C=NL/G provided by Illumina, (Where C = coverage, N = sequencing yield, L = fragment size and G = genome size), was 84,000x coverage for 1982-M6152 and 27,026x coverage for 2019-043682 (Illumina, Inc.). Multiple sequence alignment (MSA) of both isolates with M. bovis PG45 in Geneious 11 (Figure 1) revealed that 2019-043682 had 91% homology with PG45. 1982-M6152 had 97.2% homology to PG45. Annotation of the MSA in MegAlign using feature data for PG45 identified 878 unique features (genes and pseudogenes), with divergences by isolate summarized in Table 2. Features were reported as Not_Mapped by MegAlign if their %identity score fell below 95%. Unmapped features have been further categorized as excised, truncated or highly variable based on their %coverage score.

**Figure 1:**
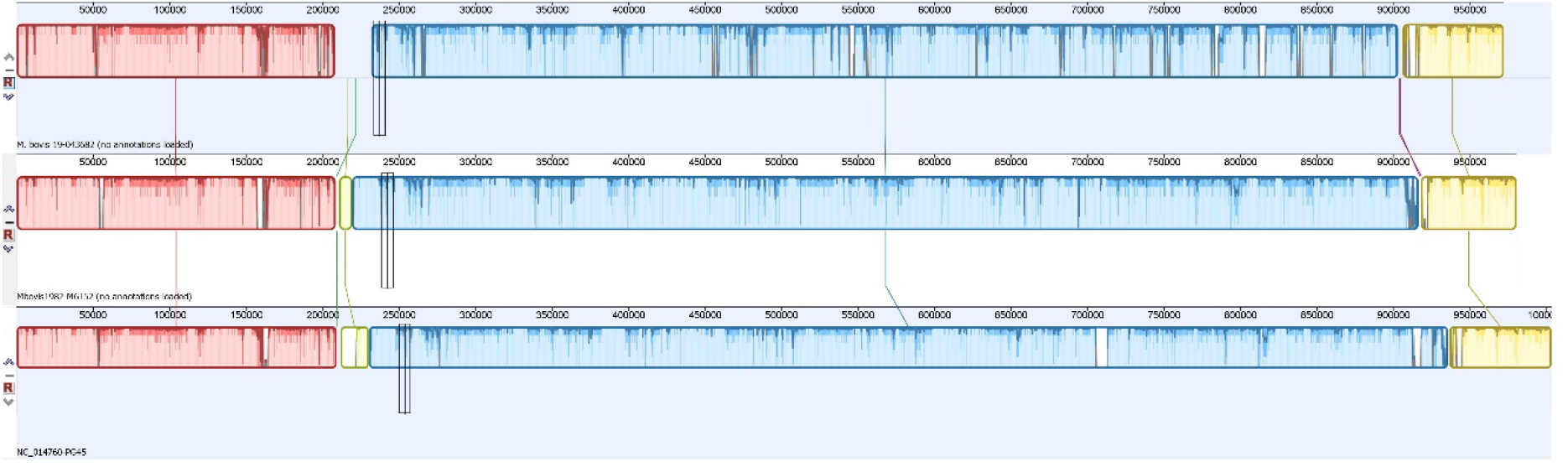
Graphical output of multiple sequence alignment (Geneious 11) for three M. bovis isolates 2019-43682 (top), 1982-M6152 (middle) and PG45 (bottom) displaying general features and depth of sequencing.

**Table 2:** Summary of variation of isolates from M. bovis PG45, by feature count, generated using MegAlign v17 for multiple sequence alignment.

| Genetic events |  | 1982_M6152 | 2019-043682 |
| --- | --- | --- | --- |
| Variation types | Identical | 696 | 183 |
|  | Deletion | 8 | 18 |
|  | Deleted_end_3prime | 3 | 3 |
|  | Deleted_end_5prime | 1 | 3 |
|  | Indel | 3 | 4 |
|  | Insertion | 22 | 22 |
|  | Not_Mapped | 39 | 173 |
|  | Substitution | 105 | 471 |
|  | **TOTAL** | **877** | **877** |
| Unmapped features | Excised (<5% coverage) | 24 | 81 |
|  | Truncated (5-95% coverage) | 11 | 48 |
|  | Highly variable (>95 % coverage) | 4 | 44 |
|  | **TOTAL** | **39** | **173** |

To determine which mutations could potentially alter gene function, further analysis using DNAStar’s ArrayStar software revealed 3285 individual nonsynonymous mutations relative to *M. bovis* PG45, across 513 genes and pseudogenes. Isolate 1982-M6152 contained 81 genes with non-synonymous mutations, 20 of which were disrupted. 507 genes in isolate 2019-043682 contained non-synonymous mutations, with 67 genes disrupted. 17 genes with non-synonymous mutations were common to both isolate 1982-M6152 and isolate 2019-043682, with 14 of the mutations identical between the 2 isolates. Four genes with disrupting mutations were common to both isolates, with an insertion mutation in gene MBOVPG45_RS03940 (insertion TTGT, genome reference position 918372) identical between isolates. A subset of 55 genes (Table 3) containing NSMs was selected for further consideration based on their functional role and known mechanisms of antimicrobials. A full list of nonsynonymous mutations, their sequence and their positions is available as supplementary data (Table S1, Table S2).

**Table 3:** Count of non-synonymous mutations (NSMs) relative to M. bovis PG45, by gene and by isolate, for a subset of NSM-containing genes identified as potentially associated with antimicrobial resistance.

| Gene: | 1982-M6152 | 2019-043682 | Description |
| --- | --- | --- | --- |
| gyrA | 0 | 8 | DNA gyrase subunit A |
| gyrB | 0 | 6 | type IIA DNA topoisomerase subunit B |
| parC | 1* | 19* | DNA topoisomerase IV subunit A |
| parE | 0 | 4 | DNA topoisomerase IV subunit B |
| topA | 0 | 1^ | type I DNA topoisomerase |
| MBOVPG45_RS00465 | 0 | 2 | tRNA (cytidine(34)-2'-O)-methyltransferase |
| MBOVPG45_RS00470 | 0 | 7 | RNA methyltransferase |
| MBOVPG45_RS02280 | 0 | 4 | 16S rRNA (uracil(1498)-N(3))-methyltransferase |
| rlmB | 0 | 5 | 23S rRNA (guanosine(2251)-2'-O)-methyltransferase RlmB |
| rlmD | 0 | 11 | 23S rRNA (uracil(1939)-C(5))-methyltransferase RlmD |
| rlmH | 0 | 1 | 23S rRNA (pseudouridine(1915)-N(3))-methyltransferase RlmH |
| rsmA | 0 | 6 | 16S rRNA (adenine(1518)-N(6)/adenine(1519)-N(6))-dimethyltransferase RsmA |
| rsmD | 0 | 1 | 16S rRNA (guanine(966)-N(2))-methyltransferase RsmD |
| rsmH | 0 | 5 | 16S rRNA (cytosine(1402)-N(4))-methyltransferase RsmH |
| rsmI | 0 | 5 | 16S rRNA (cytidine(1402)-2'-O)-methyltransferase |
| trmB | 0 | 4^ | tRNA (guanosine(46)-N7)-methyltransferase TrmB |
| rpsB | 0 | 3 | 30S ribosomal protein S2 |
| rpsC | 2 | 1 | 30S ribosomal protein S3 |
| rpsD | 0 | 2 | 30S ribosomal protein S4 |
| rpsE | 1 | 2 | 30S ribosomal protein S5 |
| rpsH | 0 | 1 | 30S ribosomal protein S8 |
| rpsJ | 1 | 0 | 30S ribosomal protein S10 |
| rpsP | 0 | 2 | 30S ribosomal protein S16 |
| rpsS | 0 | 1 | 30S ribosomal protein S19 |
| rbfA | 0 | 1 | 30S ribosome-binding factor RbfA |
| MBOVPG45_RS00445 | 0 | 1 | 50S ribosomal protein L1 |
| rplB | 0 | 1 | 50S ribosomal protein L2 |
| rplC | 0 | 1 | 50S ribosomal protein L3 |
| rplD | 0 | 10 | 50S ribosomal protein L4 |
| MBOVPG45_RS03525 | 0 | 2 | 50S ribosomal protein L10 |
| rplV | 0 | 1 | 50S ribosomal protein L22 |
| MBOVPG45_RS01360 | 0 | 1 | 50S ribosomal protein L24 |
| rpmE | 0 | 1 | 50S ribosomal protein L31 |
| alaS | 1 | 8^ | alanine--tRNA ligase |
| MBOVPG45_RS01640 | 0 | 5 | arginine--tRNA ligase |
| asnS | 0 | 1 | asparagine--tRNA ligase |
| MBOVPG45_RS00205 | 0 | 9 | class I tRNA ligase family protein |
| MBOVPG45_RS01150 | 0 | 6^ | glutamate--tRNA ligase |
| MBOVPG45_RS02730 | 0 | 2 | glycine--tRNA ligase |
| MBOVPG45_RS02640 | 0 | 1 | histidine--tRNA ligase |
| ileS | 1 | 5^ | isoleucine--tRNA ligase |
| MBOVPG45_RS03145 | 0 | 1 | isoleucine--tRNA ligase |
| MBOVPG45_RS02255 | 0 | 16 | leucine--tRNA ligase |
| lysS | 0 | 3 | lysine--tRNA ligase |
| MBOVPG45_RS03150 | 0 | 10 | methionine--tRNA ligase |
| pheS | 0 | 2 | phenylalanine--tRNA ligase subunit alpha |
| MBOVPG45_RS00380 | 0 | 18 | phenylalanine--tRNA ligase subunit beta |
| serS | 0 | 1 | serine--tRNA ligase |
| MBOVPG45_RS02170 | 1 | 8^ | threonine--tRNA ligase |
| trpS | 0 | 1 | tryptophan--tRNA ligase |
| MBOVPG45_RS04210 | 0 | 9^ | tyrosine--tRNA ligase |
| MBOVPG45_RS00740 | 0 | 7^ | valine--tRNA ligase |
| tilS | 0 | 8 | tRNA lysidine(34) synthetase TilS |
| thiI | 0 | 4 | tRNA 4-thiouridine(8) synthase ThiI |
| mnmA | 0 | 5 | tRNA 2-thiouridine(34) synthase MnmA |
| *Identical NSM. ^Contains a gene-disrupting NSM. |  |  |  |

### 2.3. Core genome multilocus sequencing Typing (cgMLST)

Isolate 2019-043682 had an undescribed cgMLST type (ST10-like) while solate 1982-M6152 had the same cgMLST type (ST17) as PG45.,.

## 3. Discussion

Isolate 2019-043682 had significantly elevated MICs for multiple fluoroquinolones, macrolides and tetracyclines, as well as a lincosamide, a pleuromutilin and two inhibitors of protein synthesis (gentamicin and florfenicol). Increased MIC is also observed for spectinomycin, indicating multi-drug resistant *M. bovis* can emerge in the field.

Of the *M. bovis* genes previously linked by Sulyok *et al* with AMR for various classes of antimicrobial (Table 4), two sites linked with fluoroquinolone resistance (*gyrA* and *gyrB*) display multiple non-synonymous mutations (NSMs) for the high-MIC isolate 2019-043682 and no NSMs for the low-MIC isolate 1982-M6152. *parC*, likewise associated with fluoroquinolone resistance, shows 18 unique NSMs in the isolate 2019-043682, and a single NSM in 1982-M6152, which is shared with the 2019-043682, therefore the shared single NSM is unlikely to be contributory to the elevated MICs.

**Table 4:** Count of non-synonymous mutations by isolate for genes previously identified (Sulyok et al, 2017) as markers for AMR. M. bovis PG45 used as reference genome. *One NSM is identical between isolates.

| Gene: | 1982-M6152 | 2019-043682 | Target Antimicrobials |
| --- | --- | --- | --- |
| *gyrA* | 0 | 8 | fluoroquinolones |
| *gyrB* | 0 | 6 | fluoroquinolones |
| *parC* | 1* | 19* | fluoroquinolones |
| *rrs1-rrs2* | 0 | 0 | tetracyclines, spectinomycin |
| *rrl1-rrl2* | 0 | 0 | macrolides, lincosamides, phenicols, pleuromutilins |

Although genes *rrs1-rrs2* and *rrl1-rrl2* were associated with AMR for tetracyclines, spectinomycin, macrolides, lincosamides and pleuromutilins, there are no NSMs for them in the isolate 2019-043682 despite the elevated MIC values, suggesting additional genetic events may be associated with AMR for these antimicrobials.

For antimicrobials where an observed increase in MIC was not matched with a previously identified *M*. *bovis* resistance-associated mutation, genes identified as AMR-associated in other species, as well as genes within the same functional groups are likely candidates for AMR association. Beyond the genes previously associated with AMR in *M. bovis*, an additional 510 features contain non-synonymous mutations. Assigning these genes to functional groups allowed us to exclude pseudogenes and genes coding for uncharacterized and hypothetical proteins. Additionally excluded were genes whose NSMs were identical between isolates 2019-043682 (high MIC) and 1982-M6152 (low MIC). Of the 149 genes remaining, we focused on a subset of 55 genes with nonsynonymous mutations within three functional roles known to be involved in antimicrobial resistance: protein synthesis, topoisomerases, and efflux pumps, specifically the ATP binding cassette (ABC) transporter system.

### 3.1. Protein Synthesis

Interference with protein synthesis is a primary method of action for the antimicrobials, with different antimicrobials interfering at different stages of synthesis, and at different locations within the ribosome complex.

#### 3.1.1. Methyltransferases

RNA methyltransferases methylate specific bases within ribosomal RNA, altering the physical structure of binding sites and other active sites within the ribosomal subunits [11–13]. Mutations within the 16S RNA methyltransferase family are known to confer aminoglycoside resistance within other bacterial species [13], and five (*rsmA, rsmD, rsmH, rsmH* and MBOVPG45_RS02280) within isolate 2019-043682 contain NSMs not found in isolate 1982-M6152. The 23S methyltransferase *rlmA* has been associated with AMR for tylosin [11–13], and while *rlmA* is wildtype in isolate 2019-043682, the related 23S methyltransferases *rlmB, rlmD* and *rlmH* contain 5, 11 and 1 unique NSMs respectively. For tRNA methyltransferases, *trmD* is implicated in multi-drug resistance [14]: it is wildtype in isolate 2019-043682, but two related tRNA methyltransferases (*trmB*, MBOVPG45_RS00465) contain NSMs, with *trmB* containing a gene disruption.

#### 3.1.2. Ribosomal Proteins

Mutations in *RpsJ*, a component of the 30S ribosomal subunit, are known to confer tetracycline resistance [15], but *RpsJ* is wildtype in the isolate 2019-043682 and contains a single NSM in the isolate 1982-M6152 that is therefore unlikely to influence MIC values. However, eight other 30S ribosomal proteins contain NSMs in the isolate 2019-043682. Of the 50S ribosomal proteins with observed NSMs, *rlpD* and *rplV* have been previously associated with macrolide resistance in *Clostridium perfringens* and two *Campylobacter* species [16, 17] with *rplD* containing ten separate NSMs in the isolate 2019-043682. A single NSM is found in the 50S subunit gene *rplC*, where mutation has been previously associated with pleuromutilin resistance [18]. 50S ribosomal proteins mutations have also been linked with resistance to lincosamides, macrolides and phenicols [19]: an additional five genes coding for 50S ribosomal proteins contain NSMs in isolate 2019-043682 while remaining wildtype in isolate 1982-M6152.

#### 3.1.3. Aminoacyl-tRNA Synthetases

While none of the antimicrobials used in MIC testing in this study target them, aminoacyl-tRNA synthetases, also known as tRNA-ligases, are enzymes which attach individual amino acids to their corresponding tRNAs and are a target of interest for antimicrobial development [20]. Isolate 2019-043682 contains 22 tRNA-ligase genes with NSMs, of which ileS (a known target for pseudomonic acid) [21] is disrupted, as is alaS (a novobiocin target)[22], in addition to a glutamate-tRNA ligase (MBOVPG45_RSO1150) and a methionine-tRNA ligase (MBOVPG45_RS03150). Three of these genes (alaS, ileS and MBOVPG45_RS02170) contain different NSMs in isolate 1982-M6152, illustrating that the presence of an NSM on its own is not sufficient for AMR, and deeper investigation into changes in protein structure and function are required.

### 3.2. Topoisomerases

In addition to *gyrA, gyrB* and *parC* discussed by Sulyok et al, *parE* mutations are also involved in fluoroquinolone resistance [23] and the isolate 2019-043682 contains 4 NSMs within the *parE* gene. While *topA*, a type I DNA topoisomerase, has not been linked with AMR previously, the gene is disrupted in the isolate 2019-043682. As bacterial topoisomerase I is a target of interest for antimicrobial development [24–26], screening for mutations affecting *topA* may be of future value to researchers and clinicians.

### 3.3. Bacterial Efflux Pumps: ABC Transporters

Bacterial efflux pumps are a class of membrane transport proteins whose role is the removal of toxic substances or metabolites from within the bacterial cell: It is estimated that 5-10% of all bacterial genes are involved in transport, with efflux pumps specifically comprising a large proportion of these transporters [27]. Of the two classes of efflux pump, primary and secondary, the primary transporters use ATP hydrolysis as an energy source, and are also known as ATP binding cassette transporters, or ABC transporters [28]. They are more commonly implicated in resistance to a single drug or category of drugs, although instances of multi-drug resistant ABC transporters have been described [13, 28].

22 ABC transporter genes contain NSMs in isolate 2019-043682, one of which contains a gene-disrupting mutation. Although none are previously identified as SDR- or MDR-involved, the wide range of antimicrobials affected by efflux pumps suggests that this may be an area of interest for future research. While 8 non-ABC membrane transport proteins with NSMs were identified in isolate 2019-043682 (Table S2), none has been characterized sufficiently to determine their potential as secondary efflux pumps and have thus been excluded from discussion.

## 4. Materials and Methods

### 4.1 Culture & Isolation of Mycoplasmas

The body of a two week old male Holstein calf was submitted to the Animal Health Lab in July of 2019 for post-mortem examination. Histologically, no lesions indicative of mycoplasma pneumonia were observed within the lungs. Culture and isolation of *M. bovis* AHL# 2019-043682 from the calf lung tissue was conducted as follows: The lung tissue submitted was perforated repeatedly using a sterile dry swab to collect sample material for broth (pig serum, horse serum and ureaplasma broths) and agar plate (pig serum agar, yeastolate agar, ureaplasma agar) culture [29]. Mycoplasma agar plates were incubated at 34-38°C with 5-7% CO_2_ and 80-100% relative humidity. Ureaplasma agar plates ware incubated at 34-38°C anaerobically. All broth cultures were incubated aerobically at 34-38°C. Plates were read at 48-72 hour intervals using a transilluminated stereomicroscope. Broth tubes were visually inspected for growth and pH change at 18-24 hour intervals, and were subcultured twice, at 48-72 hours growth and at 48-72 hours following the first subculture. Agar plates were subcultured if suspicious growth was observed during reading.

Following isolation, species identity as *M bovis* was confirmed using goat anti-rabbit/fluorescein isothiocyanate (GAR/FITC)-labelled antiserum fluorescent antibody staining [29]. Blocks of agar containing pure isolate were cut and stored at −80°C for long term storage.

Isolated 1982-M6152, an isolate of *M. bovis* from 1982 stored at −80°C and identified in a previous study [3] as low MIC for most antimicrobials, was propagated and tested by WGS and MIC retesting.

### 4.2 MIC testing

Minimum inhibitory concentration (MIC) testing was conducted on *M. bovis* isolates 2019-043682, 1982-M6152 and PG45 in triplicates for each isolate using previously described procedures [3].

### 4.3 Nucleic acid extraction

For the isolates 1982-M6152, 2019-043682 and *M. bovis* PG45, 100 μL of a broth culture was extracted on the Applied Biosystems MagMAX 96 automated nucleic acid extraction platform (Applied Biosystems, Foster City, CA) and eluted in a final volume of 90 μL, using the Low Cell Content protocol for the MagMAX Pathogen DNA/RNA kit (Applied Biosystems). Samples were held at −20 °C until prepared for WGS.

### 4.4 Whole Genome Sequencing & Bioinformatics

2×251 paired end sequencing was conducted on the Illumina MiSeq platform (Illumina, San Diego, CA) using a Nextera XT kit (Illumina) and associated protocols for whole genome sequencing (Illumina Custom Protocol Selector, Illumina Inc.)

Quality filtering and assembly of FASTQ files was done on instrument and then uploaded to Illumina’s BaseSpace storage and computing cloud. On BaseSpace, genome assembly was conducted using SPAdes Genome Assembler v3.9.0, and MLST assignment of two isolates using the Bacterial Analysis Pipeline v1.0.4.

The assembled FASTQ files were then downloaded from BaseSpace, the adapter for the Nextera XT Kit (CTGTCTCTTATACACATCT) trimmed, then aligned against and annotated with features from *M. bovis* PG45 as a reference genome using DNAstar’s SeqMan NG v14 and MegAlign v. 17 bioinformatics software (DNASTAR, Madison, WI). SNPs were analyzed with MegAlign, DNASTAR’s ArrayStar v14 (DNASTAR, Madison, WI), and Genious v11 (Auckland, New Zealand) to identify deleted and truncated genes, SNPs and non-synonymous mutations relative to *M. bovis* PG45. Gene features were then further annotated with reference to NCBI’s Gene database to identify their functional roles where possible.

## 5. Conclusions

This study identified 55 genetic events of nonsynonymous mutations and gene disruptions linked to *M. bovis* AMR. Future studies are warranted to further analyze these candidate genes, identifying the effects of the altered amino acids on protein structure and their link to AMR. Additionally, mutated genes identified in this study but currently uncharacterized may be assigned to functional groups in future.

## Supplementary Materials

For preprinting purpose, the following are included at the end of this paper: Table S1: ArrayStar Output of Nonsynonymous Mutations, Table S2: Summary of NSMs, WGS sequence data for 1982-M6152 and 2019-043682 will be made available through GenBank.

## Author Contributions

Contributor roles for this publication are defined as follows: Conceptualization, H.C; methodology, H.C., J.E. and L.L.; software, H.C.; validation, H.C., L.L., J.E.,; formal analysis, H.C. and L.L.; investigation, J.E. and L.L.; resources, H.C.; data curation, H.C and L.L.; writing—original draft preparation, L.L.; writing—review and editing, H.C. and J.E.; visualization, L.L., H.C..; supervision, H.C.; project administration, H.C.; funding acquisition, H.C. All authors have read and agreed to the published version of the manuscript.

## Acknowledgments

We greatly appreciate the technical support and advice on WGS provided by Aparna Krishnamurthy and Dr. Durda Slavic, Animal Health Laboratory, University of Guelph, Canada.

## Supplementary Materials

Table S1: ArrayStar Output of Nonsynonymous Mutations, Table S2: Summary of NSMs, WGS sequence data for 1982-M6152 and 2019-043682 will be made available through GenBank.

