## Supplemental Table 1 for "Identification of antimicrobial resistance-associated genes through whole genome sequencing of high and low MIC isolates of *Mycoplasma bovis*"

### S1_ArrayStar output nonsynonymous mutations

| Ref ID | Ref Pos | Gene Name | Common Name | 1982-M6152 | 19-043682 |
|---|---|---|---|---|---|
| NC_014760 | 375 | dnaA | | | A>G |
| NC_014760 | 1717 | MBOVPG45_RS00010 | | | G>A |
| NC_014760 | 3257 | MBOVPG45_RS00030 | | A>C | |
| NC_014760 | 4439 | MBOVPG45_RS00035 | | | T>A |
| NC_014760 | 5197 | MBOVPG45_RS00040 | | | T>C |
| NC_014760 | 5268 | MBOVPG45_RS00040 | | | A>G |
| NC_014760 | 5880 | MBOVPG45_RS00040 | | | G>C |
| NC_014760 | 6185 | MBOVPG45_RS00045 | | | T>C |
| NC_014760 | 6190 | MBOVPG45_RS00045 | | | T>C |
| NC_014760 | 6203 | MBOVPG45_RS00045 | | | TA>CT |
| NC_014760 | 7253 | MBOVPG45_RS00050 | | | C>T |
| NC_014760 | 11281 | MBOVPG45_RS00070 | | | C>T |
| NC_014760 | 14976 | MBOVPG45_RS00085 | | | G>T |
| NC_014760 | 15908 | MBOVPG45_RS00090 | | | C>A |
| NC_014760 | 22594 | MBOVPG45_RS00115 | | | T > del 1 |
| NC_014760 | 23094 | MBOVPG45_RS00115 | | TG > del 2 | TGTGTG > del 6 |
| NC_014760 | 30658 | MBOVPG45_RS00155 | | | A>T |
| NC_014760 | 36577 | MBOVPG45_RS00180 | | | G>T |
| NC_014760 | 36678 | MBOVPG45_RS00180 | | | T>C |
| NC_014760 | 36735 | MBOVPG45_RS00185 | | | T>C |
| NC_014760 | 37695 | MBOVPG45_RS00185 | | | C>G |

| | | | | | |
|---|---|---|---|---|---|
| NC_014760 | 38123 | MBOVPG45_RS00185 | | | G>A |
| NC_014760 | 38466 | MBOVPG45_RS00185 | | | C>T |
| NC_014760 | 38516 | MBOVPG45_RS00185 | | | T>C |
| NC_014760 | 38799 | MBOVPG45_RS00185 | | | T>C |
| NC_014760 | 38809 | MBOVPG45_RS00185 | | | G>T |
| NC_014760 | 38889 | MBOVPG45_RS00185 | | | C>T |
| NC_014760 | 38952 | MBOVPG45_RS00185 | | | C>T |
| NC_014760 | 41706 | MBOVPG45_RS00190 | | A>G | A>G |
| NC_014760 | 43391 | MBOVPG45_RS00190 | | | A>T |
| NC_014760 | 43393 | MBOVPG45_RS00190 | | | A>T |
| NC_014760 | 44647 | MBOVPG45_RS00190 | | | C>T |
| NC_014760 | 45249 | MBOVPG45_RS00190 | | | T>G |
| NC_014760 | 53148 | MBOVPG45_RS00205 | class I tRNA ligase family | | A>T |
| NC_014760 | 53160 | MBOVPG45_RS00205 | class I tRNA ligase family | | G>A |
| NC_014760 | 53170 | MBOVPG45_RS00205 | class I tRNA ligase family | | GA>AT |
| NC_014760 | 53230 | MBOVPG45_RS00205 | class I tRNA ligase family | | A>G |
| NC_014760 | 53609 | MBOVPG45_RS00205 | class I tRNA ligase family | | G>T |
| NC_014760 | 53716 | MBOVPG45_RS00205 | class I tRNA ligase family | | TG>CA |
| NC_014760 | 53726 | MBOVPG45_RS00205 | class I tRNA ligase family | | G>A |
| NC_014760 | 53916 | MBOVPG45_RS00205 | class I tRNA ligase family | | C>T |
| NC_014760 | 54060 | MBOVPG45_RS00205 | class I tRNA ligase family | | G>A |
| NC_014760 | 54319 | rlmB | | | T>C |
| NC_014760 | 54588 | rlmB | | | A>G |
| NC_014760 | 54593 | rlmB | | | T>A |
| NC_014760 | 54646 | rlmB | | | T>C |
| NC_014760 | 54822 | rlmB | | | A>G |
| NC_014760 | 54964 | MBOVPG45_RS00215 | | | G>A |
| NC_014760 | 55020 | MBOVPG45_RS00215 | | | T>G |
| NC_014760 | 55057 | MBOVPG45_RS00215 | | | T>A |
| NC_014760 | 55111 | MBOVPG45_RS00215 | | | A>G |
| NC_014760 | 55239 | MBOVPG45_RS00215 | | | G>A |
| NC_014760 | 55733 | secE | | | G>A |
| NC_014760 | 60006 | MBOVPG45_RS00235 | | | A>G |
| NC_014760 | 61188 | MBOVPG45_RS00240 | | | G>T |
| NC_014760 | 61321 | MBOVPG45_RS00245 | | | GG>AA |
| NC_014760 | 61648 | MBOVPG45_RS00245 | | | C>T |
| NC_014760 | 61669 | MBOVPG45_RS00245 | | | A>G |

| | | | | | |
|---|---|---|---|---|---|
| NC_014760 | 65553 | MBOVPG45_RS00270 | | | TG>AA |
| NC_014760 | 65629 | MBOVPG45_RS00270 | | | G>T |
| NC_014760 | 66075 | MBOVPG45_RS00275 | | | G>A |
| NC_014760 | 66237 | MBOVPG45_RS00275 | | | T>C |
| NC_014760 | 66243 | MBOVPG45_RS00275 | | | T>C |
| NC_014760 | 66640 | MBOVPG45_RS00275 | | | G>T |
| NC_014760 | 66759 | MBOVPG45_RS00275 | | | T>C |
| NC_014760 | 66765 | MBOVPG45_RS00275 | | | GA>TG |
| NC_014760 | 66811 | MBOVPG45_RS00275 | | | T>A |
| NC_014760 | 67139 | rsmA | | | T>C |
| NC_014760 | 67168 | rsmA | | | T>C |
| NC_014760 | 67188 | rsmA | | | T>C |
| NC_014760 | 67235 | rsmA | | | G>T |
| NC_014760 | 67257 | rsmA | | | AT>GC |
| NC_014760 | 67309 | rsmA | | | A>G |
| NC_014760 | 67846 | MBOVPG45_RS00285 | | | C>A |
| NC_014760 | 68017 | MBOVPG45_RS00285 | | | C>A |
| NC_014760 | 68171 | MBOVPG45_RS00285 | | | T>A |
| NC_014760 | 68424 | MBOVPG45_RS00285 | | | C>T |
| NC_014760 | 68496 | MBOVPG45_RS00285 | | | C>T |
| NC_014760 | 68835 | mnmE | | | C>T |
| NC_014760 | 68913 | mnmE | | | C>T |
| NC_014760 | 68925 | mnmE | | | T>C |
| NC_014760 | 68951 | mnmE | | | G>T |
| NC_014760 | 72682 | serS | | | C>T |
| NC_014760 | 73768 | MBOVPG45_RS00310 | | | A>G |
| NC_014760 | 73855 | MBOVPG45_RS00310 | | | A>G |
| NC_014760 | 74078 | MBOVPG45_RS00310 | | | T>C |
| NC_014760 | 74822 | rnpA | | | C>T |
| NC_014760 | 74964 | rnpA | | | C>T |
| NC_014760 | 75016 | rnpA | | | A>C |
| NC_014760 | 75564 | yidC | | | G>A |
| NC_014760 | 75654 | yidC | | | G>C |
| NC_014760 | 75847 | yidC | | | A>G |
| NC_014760 | 75859 | yidC | | - > ins TGCTTTTATCTAT | |
| NC_014760 | 75862 | yidC | | A>T | |
| NC_014760 | 75910 | yidC | | | A>T |
| NC_014760 | 76057 | yidC | | | A>G |
| NC_014760 | 76069 | yidC | | | C>T |
| NC_014760 | 76075 | yidC | | | T>A |
| NC_014760 | 76341 | yidC | | | A>G |
| NC_014760 | 78149 | MBOVPG45_RS00330 | | | T>A |
| NC_014760 | 78548 | MBOVPG45_RS00335 | | | T>G |
| NC_014760 | 78594 | MBOVPG45_RS00335 | | | A>T |
| NC_014760 | 78647 | MBOVPG45_RS00335 | | | G>A |
| NC_014760 | 78669 | MBOVPG45_RS00335 | | | A>G |

| | | | | | |
|---|---|---|---|---|---|
| NC_014760 | 78915 | MBOVPG45_RS00335 | | A>G | |
| NC_014760 | 78953 | MBOVPG45_RS00335 | | | C>T |
| NC_014760 | 79432 | MBOVPG45_RS00340 | | | C>T |
| NC_014760 | 80472 | MBOVPG45_RS00345 | | | A>G |
| NC_014760 | 80743 | MBOVPG45_RS00345 | | | G>C |
| NC_014760 | 80886 | MBOVPG45_RS00345 | | | G>A |
| NC_014760 | 81331 | MBOVPG45_RS00350 | | | A>T |
| NC_014760 | 81365 | MBOVPG45_RS00350 | | | A>T |
| NC_014760 | 81376 | MBOVPG45_RS00350 | | | A>G |
| NC_014760 | 81645 | MBOVPG45_RS00350 | | | A>G |
| NC_014760 | 81679 | MBOVPG45_RS00350 | | | C>T |
| NC_014760 | 82020 | MBOVPG45_RS00350 | | C>T | |
| NC_014760 | 82431 | MBOVPG45_RS00350 | | | T>C |
| NC_014760 | 82632 | MBOVPG45_RS00350 | | | T>G |
| NC_014760 | 82961 | MBOVPG45_RS00350 | | | T>A |
| NC_014760 | 83290 | MBOVPG45_RS00350 | | | G>A |
| NC_014760 | 83544 | MBOVPG45_RS00350 | | | CT>AC |
| NC_014760 | 83547 | MBOVPG45_RS00350 | | | A>G |
| NC_014760 | 83935 | MBOVPG45_RS00350 | | | TC>CT |
| NC_014760 | 84626 | MBOVPG45_RS00350 | | | A>T |
| NC_014760 | 85191 | MBOVPG45_RS00350 | | | A>C |
| NC_014760 | 86893 | MBOVPG45_RS00365 | | | A>G |
| NC_014760 | 86940 | MBOVPG45_RS00365 | | | A>C |
| NC_014760 | 87140 | MBOVPG45_RS00365 | | | C>A |
| NC_014760 | 87536 | pheS | | | G>T |
| NC_014760 | 88053 | pheS | | | G>A |
| NC_014760 | 88161 | MBOVPG45_RS00375 | | | A>T |
| NC_014760 | 88184 | MBOVPG45_RS00375 | | | T>C |
| NC_014760 | 88230 | MBOVPG45_RS00375 | | | C>A |
| NC_014760 | 88567 | MBOVPG45_RS00375 | | | T>A |
| NC_014760 | 89028 | MBOVPG45_RS00380 | | | A>G |
| NC_014760 | 89069 | MBOVPG45_RS00380 | | | G>A |
| NC_014760 | 89071 | MBOVPG45_RS00380 | | | G>T |
| NC_014760 | 89178 | MBOVPG45_RS00380 | | | A>G |
| NC_014760 | 89419 | MBOVPG45_RS00380 | | | T>A |
| NC_014760 | 89487 | MBOVPG45_RS00380 | | | T>C |
| NC_014760 | 89965 | MBOVPG45_RS00380 | | | A>T |
| NC_014760 | 90304 | MBOVPG45_RS00380 | | | T>G |
| NC_014760 | 90576 | MBOVPG45_RS00380 | | | G>A |
| NC_014760 | 90597 | MBOVPG45_RS00380 | | | T>C |
| NC_014760 | 90710 | MBOVPG45_RS00380 | | | A>G |
| NC_014760 | 90714 | MBOVPG45_RS00380 | | | TG>CA |
| NC_014760 | 90803 | MBOVPG45_RS00380 | | | C>G |
| NC_014760 | 90836 | MBOVPG45_RS00380 | | | G>T |
| NC_014760 | 90866 | MBOVPG45_RS00380 | | | T>A |
| NC_014760 | 90883 | MBOVPG45_RS00380 | | | A>T |
| NC_014760 | 90915 | MBOVPG45_RS00380 | | | C>T |

| | | | | | |
|---|---|---|---|---|---|
| NC_014760 | 90917 | MBOVPG45_RS00380 | | | A>G |
| NC_014760 | 91227 | MBOVPG45_RS00385 | | | G>A |
| NC_014760 | 91874 | MBOVPG45_RS00385 | | | G>A |
| NC_014760 | 92015 | MBOVPG45_RS00385 | | | A>G |
| NC_014760 | 92082 | MBOVPG45_RS00385 | | | C>A |
| NC_014760 | 92091 | MBOVPG45_RS00385 | | | A>G |
| NC_014760 | 92255 | pip | | | A>G |
| NC_014760 | 92270 | pip | | | C>T |
| NC_014760 | 92524 | pip | | | T>G |
| NC_014760 | 92566 | pip | | | A>G |
| NC_014760 | 92568 | pip | | | C>T |
| NC_014760 | 92613 | pip | | | C>T |
| NC_014760 | 93133 | pip | | | G>A |
| NC_014760 | 93312 | MBOVPG45_RS00395 | | | G>A |
| NC_014760 | 93414 | MBOVPG45_RS00395 | | | A>G |
| NC_014760 | 93573 | MBOVPG45_RS00395 | | | A>G |
| NC_014760 | 93616 | MBOVPG45_RS00395 | | | CC>TT |
| NC_014760 | 93898 | MBOVPG45_RS00395 | | | AT>TC |
| NC_014760 | 93952 | MBOVPG45_RS00395 | | | A>G |
| NC_014760 | 94105 | MBOVPG45_RS00395 | | | T>C |
| NC_014760 | 94135 | MBOVPG45_RS00395 | | | AG>GA |
| NC_014760 | 94168 | MBOVPG45_RS00395 | | | A>G |
| NC_014760 | 94170 | MBOVPG45_RS00395 | | | A>G |
| NC_014760 | 94484 | MBOVPG45_RS00400 | | | A>T |
| NC_014760 | 94555 | MBOVPG45_RS00400 | | | A>G |
| NC_014760 | 94589 | MBOVPG45_RS00400 | | | A>G |
| NC_014760 | 94603 | MBOVPG45_RS00400 | | | A>T |
| NC_014760 | 94639 | MBOVPG45_RS00400 | | | G>A |
| NC_014760 | 94673 | MBOVPG45_RS00400 | | | A>C |
| NC_014760 | 94738 | MBOVPG45_RS00400 | | | A>G |
| NC_014760 | 94954 | MBOVPG45_RS00405 | | | A>G |
| NC_014760 | 95286 | MBOVPG45_RS00405 | | | A>C |
| NC_014760 | 95430 | MBOVPG45_RS00405 | | | T>C |
| NC_014760 | 95724 | MBOVPG45_RS00405 | | | T>C |
| NC_014760 | 95798 | MBOVPG45_RS00405 | | | G>T |
| NC_014760 | 95873 | MBOVPG45_RS00405 | | | C>A |
| NC_014760 | 96345 | MBOVPG45_RS00410 | | | A>G |
| NC_014760 | 96681 | MBOVPG45_RS00410 | | | TG>CA |
| NC_014760 | 97237 | MBOVPG45_RS00415 | | | T>G |
| NC_014760 | 97602 | MBOVPG45_RS00415 | | A>G | A>G |
| NC_014760 | 97890 | MBOVPG45_RS00420 | | | C>T |
| NC_014760 | 98154 | MBOVPG45_RS00420 | | | C>T |
| NC_014760 | 98204 | MBOVPG45_RS00420 | | | G>A |
| NC_014760 | 98207 | MBOVPG45_RS00420 | | | AT>GC |
| NC_014760 | 98415 | MBOVPG45_RS00420 | | | GC>AT |
| NC_014760 | 98477 | MBOVPG45_RS00420 | | | C>T |
| NC_014760 | 98682 | MBOVPG45_RS00420 | | | T>C |

| | | | | | |
|---|---|---|---|---|---|
| NC_014760 | 98687 | MBOVPG45_RS00420 | | | G>A |
| NC_014760 | 98697 | MBOVPG45_RS00420 | | | AC>GT |
| NC_014760 | 98761 | MBOVPG45_RS00420 | | | A>T |
| NC_014760 | 98822 | MBOVPG45_RS00420 | | | G>A |
| NC_014760 | 98831 | MBOVPG45_RS00420 | | | G>A |
| NC_014760 | 98853 | MBOVPG45_RS00420 | | | A>T |
| NC_014760 | 99221 | MBOVPG45_RS00425 | | | A>G |
| NC_014760 | 99224 | MBOVPG45_RS00425 | | | C>A |
| NC_014760 | 99358 | fba | | | T>C |
| NC_014760 | 99432 | fba | | | TA>CG |
| NC_014760 | 99917 | fba | | | C>T |
| NC_014760 | 100528 | MBOVPG45_RS00435 | | | T>A |
| NC_014760 | 100586 | MBOVPG45_RS00435 | | | G>A |
| NC_014760 | 100588 | MBOVPG45_RS00435 | | | TT>CC |
| NC_014760 | 102115 | MBOVPG45_RS00445 | | | T>C |
| NC_014760 | 102344 | MBOVPG45_RS00450 | | | A>G |
| NC_014760 | 102351 | MBOVPG45_RS00450 | | | A>T |
| NC_014760 | 102358 | MBOVPG45_RS00450 | | | A>G |
| NC_014760 | 104072 | MBOVPG45_RS00460 | | | AAT > del 3 |
| NC_014760 | 104112 | MBOVPG45_RS00460 | | | C>T |
| NC_014760 | 104133 | MBOVPG45_RS00460 | | | G>A |
| NC_014760 | 104148 | MBOVPG45_RS00460 | | | A>G |
| NC_014760 | 104430 | MBOVPG45_RS00460 | | | G>A |
| NC_014760 | 104443 | MBOVPG45_RS00460 | | | C>T |
| NC_014760 | 104541 | MBOVPG45_RS00460 | | | T>G |
| NC_014760 | 104555 | MBOVPG45_RS00460 | | | A>C |
| NC_014760 | 104568 | MBOVPG45_RS00460 | | | T>C |
| NC_014760 | 104581 | MBOVPG45_RS00460 | | | G>A |
| NC_014760 | 104591 | MBOVPG45_RS00460 | | | T>G |
| NC_014760 | 104611 | MBOVPG45_RS00460 | | | T>A |
| NC_014760 | 104616 | MBOVPG45_RS00460 | | | T>C |
| NC_014760 | 104626 | MBOVPG45_RS00460 | | | T>C |
| NC_014760 | 104635 | MBOVPG45_RS00460 | | | TG>GA |
| NC_014760 | 104650 | MBOVPG45_RS00460 | | | T>C |
| NC_014760 | 105222 | MBOVPG45_RS00465 | | | C>T |
| NC_014760 | 105388 | MBOVPG45_RS00465 | | | A>G |
| NC_014760 | 105633 | MBOVPG45_RS00470 | | | A>G |
| NC_014760 | 105688 | MBOVPG45_RS00470 | | | A>C |
| NC_014760 | 105782 | MBOVPG45_RS00470 | | | A>G |
| NC_014760 | 105858 | MBOVPG45_RS00470 | | | C>T |
| NC_014760 | 105968 | MBOVPG45_RS00470 | | | G>A |
| NC_014760 | 106091 | MBOVPG45_RS00470 | | | G>T |
| NC_014760 | 106097 | MBOVPG45_RS00470 | | | A>G |
| NC_014760 | 106274 | MBOVPG45_RS00475 | | | G>A |
| NC_014760 | 106355 | MBOVPG45_RS00475 | | | A>G |
| NC_014760 | 106387 | MBOVPG45_RS00475 | | | T>A |
| NC_014760 | 106703 | ylqF | | | T>C |

| | | | | | |
|---|---|---|---|---|---|
| NC_014760 | 106787 | ylqF | | | T>C |
| NC_014760 | 107167 | ylqF | | | G>A |
| NC_014760 | 107419 | MBOVPG45_RS00485 | | | G>A |
| NC_014760 | 107455 | MBOVPG45_RS00485 | | | A>G |
| NC_014760 | 107518 | MBOVPG45_RS00485 | | G>T | G>T |
| NC_014760 | 107531 | MBOVPG45_RS00485 | | | C>T |
| NC_014760 | 107716 | MBOVPG45_RS00485 | | | T>G |
| NC_014760 | 107976 | MBOVPG45_RS00490 | | | G>C |
| NC_014760 | 108118 | MBOVPG45_RS00490 | | | A>G |
| NC_014760 | 108131 | MBOVPG45_RS00490 | | | T>C |
| NC_014760 | 108358 | MBOVPG45_RS00490 | | | G>A |
| NC_014760 | 108881 | MBOVPG45_RS00495 | | | G>A |
| NC_014760 | 109283 | MBOVPG45_RS00495 | | | A>G |
| NC_014760 | 109687 | MBOVPG45_RS00495 | | | C>A |
| NC_014760 | 109760 | MBOVPG45_RS00495 | | | C>T |
| NC_014760 | 110067 | MBOVPG45_RS00500 | | | A>G |
| NC_014760 | 110917 | MBOVPG45_RS00505 | | | TC>CT |
| NC_014760 | 110986 | MBOVPG45_RS00505 | | | T>G |
| NC_014760 | 111027 | MBOVPG45_RS00505 | | | C>T |
| NC_014760 | 111031 | MBOVPG45_RS00505 | | | C>T |
| NC_014760 | 111128 | MBOVPG45_RS00505 | | | A>T |
| NC_014760 | 111274 | MBOVPG45_RS00505 | | | A>T |
| NC_014760 | 111288 | MBOVPG45_RS00505 | | | AT>GC |
| NC_014760 | 111369 | MBOVPG45_RS00505 | | | G>A |
| NC_014760 | 111454 | MBOVPG45_RS00505 | | | C>T |
| NC_014760 | 111456 | MBOVPG45_RS00505 | | | C>T |
| NC_014760 | 115075 | lpdA | | C>T | |
| NC_014760 | 116070 | lpdA | | | C>T |
| NC_014760 | 117143 | MBOVPG45_RS00540 | | | G>C |
| NC_014760 | 117345 | MBOVPG45_RS00540 | | | C>T |
| NC_014760 | 117352 | MBOVPG45_RS00540 | | | T>A |
| NC_014760 | 118227 | MBOVPG45_RS00545 | | | A>G |
| NC_014760 | 119112 | MBOVPG45_RS00550 | | | A>T |
| NC_014760 | 119162 | MBOVPG45_RS00550 | | | G>A |
| NC_014760 | 119276 | MBOVPG45_RS00550 | | | A>C |
| NC_014760 | 119321 | MBOVPG45_RS00550 | | | A>T |
| NC_014760 | 119337 | MBOVPG45_RS00550 | | A>C | A>C |
| NC_014760 | 119368 | MBOVPG45_RS00550 | | | A>G |
| NC_014760 | 119481 | MBOVPG45_RS00550 | | | TG>CA |
| NC_014760 | 119491 | MBOVPG45_RS00550 | | | A>G |
| NC_014760 | 119496 | MBOVPG45_RS00550 | | | TGG>GAA |
| NC_014760 | 119500 | MBOVPG45_RS00550 | | | A>C |
| NC_014760 | 119529 | MBOVPG45_RS00550 | | | CA>TG |
| NC_014760 | 119623 | MBOVPG45_RS00550 | | | G>A |
| NC_014760 | 119730 | MBOVPG45_RS00550 | | | AG>GC |
| NC_014760 | 119854 | MBOVPG45_RS00550 | | | C>G |
| NC_014760 | 119864 | MBOVPG45_RS00550 | | | A>G |

| | | | | | |
|---|---|---|---|---|---|
| NC_014760 | 119913 | MBOVPG45_RS00550 | | | CT>TG |
| NC_014760 | 119980 | MBOVPG45_RS00550 | | | A>C |
| NC_014760 | 120041 | MBOVPG45_RS00550 | | | A>T |
| NC_014760 | 120062 | MBOVPG45_RS00550 | | | A>G |
| NC_014760 | 120126 | MBOVPG45_RS00550 | | | G>T |
| NC_014760 | 120130 | MBOVPG45_RS00550 | | | C>T |
| NC_014760 | 120198 | MBOVPG45_RS00550 | | | T>A |
| NC_014760 | 120206 | MBOVPG45_RS00550 | | | G>A |
| NC_014760 | 120274 | MBOVPG45_RS00550 | | | G>C |
| NC_014760 | 120279 | MBOVPG45_RS00550 | | | T>A |
| NC_014760 | 120283 | MBOVPG45_RS00550 | | | G>A |
| NC_014760 | 120381 | MBOVPG45_RS00550 | | | G>T |
| NC_014760 | 120431 | MBOVPG45_RS00550 | | | C>T |
| NC_014760 | 120610 | MBOVPG45_RS00550 | | | A>G |
| NC_014760 | 120657 | MBOVPG45_RS00550 | | | A>T |
| NC_014760 | 120669 | MBOVPG45_RS00550 | | | A>T |
| NC_014760 | 120752 | MBOVPG45_RS00550 | | | C>T |
| NC_014760 | 120905 | MBOVPG45_RS00550 | | | G>A |
| NC_014760 | 120935 | MBOVPG45_RS00550 | | | C>A |
| NC_014760 | 121116 | MBOVPG45_RS00550 | | | TA>CC |
| NC_014760 | 121128 | MBOVPG45_RS00550 | | | T>A |
| NC_014760 | 121158 | MBOVPG45_RS00550 | | | A>G |
| NC_014760 | 121309 | MBOVPG45_RS00550 | | | G>T |
| NC_014760 | 121454 | MBOVPG45_RS00550 | | | G>A |
| NC_014760 | 121490 | MBOVPG45_RS00550 | | | A>C |
| NC_014760 | 121582 | MBOVPG45_RS00550 | | | G>A |
| NC_014760 | 121584 | MBOVPG45_RS00550 | | | T>A |
| NC_014760 | 121622 | MBOVPG45_RS00550 | | | G>A |
| NC_014760 | 121624 | MBOVPG45_RS00550 | | | G>A |
| NC_014760 | 121640 | MBOVPG45_RS00550 | | | G>A |
| NC_014760 | 121655 | MBOVPG45_RS00550 | | | A>C |
| NC_014760 | 121714 | MBOVPG45_RS00550 | | | A>G |
| NC_014760 | 121720 | MBOVPG45_RS00550 | | | C>T |
| NC_014760 | 122978 | MBOVPG45_RS00555 | | | A>G |
| NC_014760 | 122991 | MBOVPG45_RS00555 | | | G>A |
| NC_014760 | 126802 | MBOVPG45_RS00570 | | | G>A |
| NC_014760 | 126809 | MBOVPG45_RS00570 | | | A>G |
| NC_014760 | 126829 | MBOVPG45_RS00570 | | | C>G |
| NC_014760 | 127076 | MBOVPG45_RS00570 | | | A>T |
| NC_014760 | 128082 | MBOVPG45_RS00575 | | T>A | |
| NC_014760 | 128470 | MBOVPG45_RS00575 | | | G>A |
| NC_014760 | 129778 | MBOVPG45_RS00585 | | | C>A |
| NC_014760 | 132243 | MBOVPG45_RS00590 | | | C>T |
| NC_014760 | 132696 | MBOVPG45_RS00590 | | | C>T |
| NC_014760 | 132891 | MBOVPG45_RS00595 | | | T>C |
| NC_014760 | 133930 | MBOVPG45_RS00600 | | | C>T |
| NC_014760 | 141369 | MBOVPG45_RS00640 | | | G>A |

| | | | | | |
|---|---|---|---|---|---|
| NC_014760 | 142341 | MBOVPG45_RS00640 | | A>G | |
| NC_014760 | 144605 | MBOVPG45_RS00650 | | | G>A |
| NC_014760 | 144619 | MBOVPG45_RS00650 | | | A>G |
| NC_014760 | 144920 | MBOVPG45_RS00650 | | | C>T |
| NC_014760 | 145304 | MBOVPG45_RS00650 | | | C>T |
| NC_014760 | 145513 | MBOVPG45_RS00650 | | | C>T |
| NC_014760 | 145685 | MBOVPG45_RS00650 | | | T>C |
| NC_014760 | 145789 | MBOVPG45_RS00650 | | | G>A |
| NC_014760 | 145813 | MBOVPG45_RS00650 | | | GT>TC |
| NC_014760 | 145896 | MBOVPG45_RS00650 | | A>T | |
| NC_014760 | 146413 | pepF | | | T>C |
| NC_014760 | 146541 | pepF | | | T>C |
| NC_014760 | 147487 | pepF | | | T>A |
| NC_014760 | 147489 | pepF | | | C>T |
| NC_014760 | 154022 | MBOVPG45_RS00690 | | | G>A |
| NC_014760 | 154045 | MBOVPG45_RS00690 | | | C>T |
| NC_014760 | 154170 | MBOVPG45_RS00690 | | | T>A |
| NC_014760 | 154333 | MBOVPG45_RS00690 | | | C>T |
| NC_014760 | 154382 | MBOVPG45_RS00690 | | | G>A |
| NC_014760 | 154405 | MBOVPG45_RS00690 | | | G>A |
| NC_014760 | 154409 | MBOVPG45_RS00690 | | | - > ins A |
| NC_014760 | 154597 | MBOVPG45_RS00690 | | | T>A |
| NC_014760 | 154739 | MBOVPG45_RS00690 | | | G>A |
| NC_014760 | 154975 | MBOVPG45_RS00690 | | | G>A |
| NC_014760 | 155075 | MBOVPG45_RS00690 | | | C>T |
| NC_014760 | 155102 | MBOVPG45_RS00690 | | | C>T |
| NC_014760 | 155399 | thiI | | | G>A |
| NC_014760 | 155411 | thiI | | | A>C |
| NC_014760 | 155489 | thiI | | | A>G |
| NC_014760 | 156302 | thiI | | | G>A |
| NC_014760 | 156567 | MBOVPG45_RS00700 | | A>T | |
| NC_014760 | 157477 | MBOVPG45_RS00705 | | | G>A |
| NC_014760 | 157703 | MBOVPG45_RS00705 | | | T>A |
| NC_014760 | 158334 | MBOVPG45_RS00705 | | | T>A |
| NC_014760 | 158416 | MBOVPG45_RS00705 | | | A>G |
| NC_014760 | 158505 | MBOVPG45_RS00710 | | A>C | |
| NC_014760 | 158545 | MBOVPG45_RS00710 | | | C>G |
| NC_014760 | 158547 | MBOVPG45_RS00710 | | | GG>TC |
| NC_014760 | 158619 | MBOVPG45_RS00710 | | | GC>TT |
| NC_014760 | 158646 | MBOVPG45_RS00710 | | | G>T |
| NC_014760 | 158687 | MBOVPG45_RS00710 | | | C>T |
| NC_014760 | 158695 | MBOVPG45_RS00710 | | | TA>CT |
| NC_014760 | 158723 | MBOVPG45_RS00710 | | | C>T |
| NC_014760 | 158738 | MBOVPG45_RS00710 | | | T>G |
| NC_014760 | 158746 | MBOVPG45_RS00710 | | | A>T |
| NC_014760 | 158768 | MBOVPG45_RS00710 | | | C>A |
| NC_014760 | 158815 | MBOVPG45_RS00710 | | | C>T |

| | | | | | |
|---|---|---|---|---|---|
| NC_014760 | 158826 | MBOVPG45_RS00710 | | | CC>TT |
| NC_014760 | 158912 | MBOVPG45_RS00710 | | | G>C |
| NC_014760 | 158932 | MBOVPG45_RS00710 | | | AT>GA |
| NC_014760 | 159038 | MBOVPG45_RS00710 | | | T>C |
| NC_014760 | 159083 | MBOVPG45_RS00710 | | | C>T |
| NC_014760 | 159088 | MBOVPG45_RS00710 | | | T>C |
| NC_014760 | 159092 | MBOVPG45_RS00710 | | | C>T |
| NC_014760 | 159115 | MBOVPG45_RS00710 | | | T>A |
| NC_014760 | 159117 | MBOVPG45_RS00710 | | | GC>AT |
| NC_014760 | 159131 | MBOVPG45_RS00710 | | | C>T |
| NC_014760 | 159136 | MBOVPG45_RS00710 | | | A>T |
| NC_014760 | 159220 | MBOVPG45_RS00710 | | | T>A |
| NC_014760 | 162561 | MBOVPG45_RS00725 | | | C>G |
| NC_014760 | 162573 | MBOVPG45_RS00725 | | | A>T |
| NC_014760 | 162661 | MBOVPG45_RS00725 | | | T>A |
| NC_014760 | 162852 | MBOVPG45_RS00725 | | | CGA>GAG |
| NC_014760 | 163046 | MBOVPG45_RS00725 | | | C>T |
| NC_014760 | 163071 | MBOVPG45_RS00725 | | | C>T |
| NC_014760 | 163084 | MBOVPG45_RS00725 | | | C>A |
| NC_014760 | 163100 | MBOVPG45_RS00725 | | | T>C |
| NC_014760 | 163136 | MBOVPG45_RS00725 | | | T>C |
| NC_014760 | 163148 | MBOVPG45_RS00725 | | | C>G |
| NC_014760 | 163155 | MBOVPG45_RS00725 | | | T>C |
| NC_014760 | 163167 | MBOVPG45_RS00725 | | | C>T |
| NC_014760 | 163183 | MBOVPG45_RS00725 | | | G>T |
| NC_014760 | 163287 | MBOVPG45_RS00725 | | | T>C |
| NC_014760 | 163345 | MBOVPG45_RS00725 | | A>C | A>C |
| NC_014760 | 163427 | MBOVPG45_RS00725 | | | - > ins CT |
| NC_014760 | 163428 | MBOVPG45_RS00725 | | - > ins C | A>C |
| NC_014760 | 164311 | MBOVPG45_RS00730 | | | C>G |
| NC_014760 | 164323 | MBOVPG45_RS00730 | | | A>T |
| NC_014760 | 164411 | MBOVPG45_RS00730 | | | T>A |
| NC_014760 | 164602 | MBOVPG45_RS00730 | | | CGA>GAG |
| NC_014760 | 164796 | MBOVPG45_RS00730 | | | C>T |
| NC_014760 | 164821 | MBOVPG45_RS00730 | | | C>T |
| NC_014760 | 164834 | MBOVPG45_RS00730 | | | C>A |
| NC_014760 | 164850 | MBOVPG45_RS00730 | | | T>C |
| NC_014760 | 164886 | MBOVPG45_RS00730 | | | T>C |
| NC_014760 | 164898 | MBOVPG45_RS00730 | | | C>G |
| NC_014760 | 164905 | MBOVPG45_RS00730 | | | T>C |
| NC_014760 | 164917 | MBOVPG45_RS00730 | | | C>T |
| NC_014760 | 164933 | MBOVPG45_RS00730 | | | G>T |
| NC_014760 | 165037 | MBOVPG45_RS00730 | | | T>C |
| NC_014760 | 165095 | MBOVPG45_RS00730 | | A>C | A>C |
| NC_014760 | 165178 | MBOVPG45_RS00730 | | | A > del 1 |
| NC_014760 | 165866 | MBOVPG45_RS00735 | | | C>T |
| NC_014760 | 165923 | MBOVPG45_RS00735 | | | G>A |

| | | | | | |
|---|---|---|---|---|---|
| NC_014760 | 165934 | MBOVPG45_RS00735 | | | C>T |
| NC_014760 | 167558 | MBOVPG45_RS00735 | | | A>G |
| NC_014760 | 167648 | MBOVPG45_RS00735 | | | C>A |
| NC_014760 | 167717 | MBOVPG45_RS00735 | | | C>T |
| NC_014760 | 170114 | MBOVPG45_RS00740 | | | A>G |
| NC_014760 | 170155 | MBOVPG45_RS00740 | | | G>A |
| NC_014760 | 170211 | MBOVPG45_RS00740 | | | CG>TA |
| NC_014760 | 170358 | MBOVPG45_RS00740 | | | A>G |
| NC_014760 | 170626 | MBOVPG45_RS00740 | | | G>T |
| NC_014760 | 170808 | MBOVPG45_RS00740 | | | G>A |
| NC_014760 | 170867 | MBOVPG45_RS00740 | | | G>A |
| NC_014760 | 170942 | MBOVPG45_RS00745 | | | C>T |
| NC_014760 | 171669 | MBOVPG45_RS00745 | | | GT>AA |
| NC_014760 | 172051 | MBOVPG45_RS00750 | | | C>G |
| NC_014760 | 172067 | MBOVPG45_RS00750 | | | G>A |
| NC_014760 | 172129 | MBOVPG45_RS00750 | | | G>T |
| NC_014760 | 172843 | MBOVPG45_RS00750 | | | G>A |
| NC_014760 | 173363 | MBOVPG45_RS00755 | | | A>G |
| NC_014760 | 174043 | MBOVPG45_RS00755 | | | - > ins ATAA |
| NC_014760 | 174264 | MBOVPG45_RS00760 | | | A>G |
| NC_014760 | 174840 | MBOVPG45_RS00765 | | | C>T |
| NC_014760 | 174980 | MBOVPG45_RS00765 | | | A>T |
| NC_014760 | 175008 | MBOVPG45_RS00765 | | | C>T |
| NC_014760 | 175061 | MBOVPG45_RS00765 | | | G>A |
| NC_014760 | 175241 | MBOVPG45_RS00770 | | | C>T |
| NC_014760 | 175259 | MBOVPG45_RS00770 | | | C>T |
| NC_014760 | 175673 | MBOVPG45_RS00770 | | | GA>AC |
| NC_014760 | 176049 | MBOVPG45_RS00770 | | | A>G |
| NC_014760 | 176128 | MBOVPG45_RS00770 | | | A>T |
| NC_014760 | 176136 | MBOVPG45_RS00770 | | | G>A |
| NC_014760 | 176156 | MBOVPG45_RS00770 | | | G>A |
| NC_014760 | 176162 | MBOVPG45_RS00770 | | | A>G |
| NC_014760 | 176282 | MBOVPG45_RS00770 | | | G>A |
| NC_014760 | 176291 | MBOVPG45_RS00770 | | | A>G |
| NC_014760 | 176293 | MBOVPG45_RS00770 | | | T>A |
| NC_014760 | 176378 | MBOVPG45_RS00770 | | | A>T |
| NC_014760 | 176402 | MBOVPG45_RS00770 | | | A>G |
| NC_014760 | 176413 | MBOVPG45_RS00770 | | | T>A |
| NC_014760 | 176418 | MBOVPG45_RS00770 | | | CT>AG |
| NC_014760 | 176458 | MBOVPG45_RS00770 | | | A>T |
| NC_014760 | 176531 | MBOVPG45_RS00770 | | | A>G |
| NC_014760 | 176536 | MBOVPG45_RS00770 | | | T>A |
| NC_014760 | 176544 | MBOVPG45_RS00770 | | | G>A |
| NC_014760 | 176556 | MBOVPG45_RS00770 | | | G>A |
| NC_014760 | 176561 | MBOVPG45_RS00770 | | | C>T |
| NC_014760 | 176577 | MBOVPG45_RS00770 | | | CATCAT > del 6 |
| NC_014760 | 178094 | pyk | | | A>T |

| | | | | | |
|---|---|---|---|---|---|
| NC_014760 | 178282 | MBOVPG45_RS00780 | | | CG>TA |
| NC_014760 | 178366 | MBOVPG45_RS00780 | | | A>G |
| NC_014760 | 178534 | MBOVPG45_RS00780 | | | T>C |
| NC_014760 | 178566 | MBOVPG45_RS00780 | | | A>G |
| NC_014760 | 178723 | MBOVPG45_RS00780 | | | TA>CT |
| NC_014760 | 178734 | MBOVPG45_RS00780 | | | A>T |
| NC_014760 | 178742 | MBOVPG45_RS00780 | | | A>T |
| NC_014760 | 178744 | MBOVPG45_RS00780 | | | T>G |
| NC_014760 | 178788 | MBOVPG45_RS00780 | | | T>A |
| NC_014760 | 178794 | MBOVPG45_RS00780 | | | T>G |
| NC_014760 | 178804 | MBOVPG45_RS00780 | | | T>C |
| NC_014760 | 178930 | MBOVPG45_RS00780 | | | C>T |
| NC_014760 | 178967 | MBOVPG45_RS00780 | | | C>A |
| NC_014760 | 178970 | MBOVPG45_RS00780 | | | C>A |
| NC_014760 | 179088 | MBOVPG45_RS00780 | | | G>A |
| NC_014760 | 179091 | MBOVPG45_RS00780 | | | G>A |
| NC_014760 | 179120 | MBOVPG45_RS00780 | | | GC>AT |
| NC_014760 | 179140 | MBOVPG45_RS00780 | | | C>T |
| NC_014760 | 179169 | MBOVPG45_RS00780 | | | A>G |
| NC_014760 | 180543 | dnaK | | | A>T |
| NC_014760 | 181213 | dnaK | | | - > ins ATTCATCAAATA ATA |
| NC_014760 | 181437 | MBOVPG45_RS00790 | | | T>C |
| NC_014760 | 181543 | MBOVPG45_RS00790 | | | A>C |
| NC_014760 | 181664 | MBOVPG45_RS00790 | | | T>C |
| NC_014760 | 181738 | MBOVPG45_RS00790 | | | C>T |
| NC_014760 | 182080 | MBOVPG45_RS00790 | | | T>C |
| NC_014760 | 182093 | MBOVPG45_RS00790 | | | C>A |
| NC_014760 | 182535 | MBOVPG45_RS00790 | | | T>A |
| NC_014760 | 182781 | MBOVPG45_RS00795 | | | C>T |
| NC_014760 | 182786 | MBOVPG45_RS00795 | | | G>A |
| NC_014760 | 182922 | MBOVPG45_RS00795 | | | A>G |
| NC_014760 | 183309 | MBOVPG45_RS00800 | | C>T | |
| NC_014760 | 183325 | MBOVPG45_RS00800 | | | C>G |
| NC_014760 | 183406 | MBOVPG45_RS00800 | | | T>C |
| NC_014760 | 183452 | MBOVPG45_RS00800 | | | A>T |
| NC_014760 | 183483 | MBOVPG45_RS00800 | | | T>G |
| NC_014760 | 183733 | MBOVPG45_RS00800 | | | A>T |
| NC_014760 | 184166 | MBOVPG45_RS00800 | | | AC>GT |
| NC_014760 | 184839 | MBOVPG45_RS00805 | | | T>C |
| NC_014760 | 184992 | MBOVPG45_RS00805 | | | C>T |
| NC_014760 | 185171 | MBOVPG45_RS00805 | | | T>C |
| NC_014760 | 185373 | MBOVPG45_RS00805 | | | A>G |
| NC_014760 | 185377 | MBOVPG45_RS00805 | | | G>T |
| NC_014760 | 185391 | MBOVPG45_RS00805 | | | GA>TG |
| NC_014760 | 185839 | MBOVPG45_RS04405 | | | T>A |

| | | | | | |
|---|---|---|---|---|---|
| NC_014760 | 185853 | MBOVPG45_RS04405 | | | T>C |
| NC_014760 | 185879 | MBOVPG45_RS04405 | | | C>G |
| NC_014760 | 185887 | MBOVPG45_RS04405 | | | C>T |
| NC_014760 | 185914 | MBOVPG45_RS04405 | | | G>A |
| NC_014760 | 186667 | MBOVPG45_RS00810 | | | AA>GC |
| NC_014760 | 187768 | MBOVPG45_RS00815 | | | T>C |
| NC_014760 | 187777 | MBOVPG45_RS00815 | | | C>A |
| NC_014760 | 187833 | MBOVPG45_RS00815 | | | T>C |
| NC_014760 | 188284 | MBOVPG45_RS00815 | | | G>C |
| NC_014760 | 188493 | MBOVPG45_RS00815 | | | G>A |
| NC_014760 | 189057 | MBOVPG45_RS00815 | | | G>A |
| NC_014760 | 189144 | MBOVPG45_RS00815 | | | T>A |
| NC_014760 | 189232 | MBOVPG45_RS00815 | | | C>T |
| NC_014760 | 189991 | MBOVPG45_RS00815 | | | C>T |
| NC_014760 | 195409 | tig | | | T>C |
| NC_014760 | 195949 | tig | | | T>C |
| NC_014760 | 196340 | tig | | | A>T |
| NC_014760 | 200561 | MBOVPG45_RS00865 | | | A>T |
| NC_014760 | 201033 | MBOVPG45_RS00865 | | | T>A |
| NC_014760 | 201195 | MBOVPG45_RS00870 | | | C>T |
| NC_014760 | 202195 | MBOVPG45_RS00870 | | | T>G |
| NC_014760 | 202199 | MBOVPG45_RS00870 | | | C>T |
| NC_014760 | 202367 | MBOVPG45_RS00870 | | | C>A |
| NC_014760 | 202399 | MBOVPG45_RS00870 | | | T>C |
| NC_014760 | 202533 | MBOVPG45_RS00870 | | | C>T |
| NC_014760 | 202750 | MBOVPG45_RS00870 | | | G>T |
| NC_014760 | 202925 | MBOVPG45_RS00870 | | | C>T |
| NC_014760 | 203126 | MBOVPG45_RS00870 | | | C>T |
| NC_014760 | 203140 | MBOVPG45_RS00870 | | | C>T |
| NC_014760 | 203247 | MBOVPG45_RS00870 | | | C>T |
| NC_014760 | 203249 | MBOVPG45_RS00870 | | | T>C |
| NC_014760 | 203387 | MBOVPG45_RS00870 | | | C>T |
| NC_014760 | 218020 | MBOVPG45_RS00935 | | | TC>CT |
| NC_014760 | 218129 | MBOVPG45_RS00935 | | | C>T |
| NC_014760 | 218227 | MBOVPG45_RS00935 | | | T>G |
| NC_014760 | 222893 | MBOVPG45_RS00955 | | | C>T |
| NC_014760 | 224139 | MBOVPG45_RS00960 | | | G>A |
| NC_014760 | 230209 | MBOVPG45_RS00975 | | | A>T |
| NC_014760 | 231066 | MBOVPG45_RS00975 | | | C>T |
| NC_014760 | 235507 | MBOVPG45_RS01005 | | A>T | |
| NC_014760 | 235934 | MBOVPG45_RS01005 | | T > del 1 | |
| NC_014760 | 239758 | MBOVPG45_RS01020 | | | C>T |
| NC_014760 | 241469 | MBOVPG45_RS01030 | | | A>G |
| NC_014760 | 241882 | MBOVPG45_RS01030 | | | - > ins T |
| NC_014760 | 242137 | MBOVPG45_RS01035 | | | T>A |
| NC_014760 | 242155 | MBOVPG45_RS01035 | | | TTCTAGTTC > del 9 |

| | | | | | |
|---|---|---|---|---|---|
| NC_014760 | 242169 | MBOVPG45_RS01035 | | | C>T |
| NC_014760 | 242172 | MBOVPG45_RS01035 | | | T>C |
| NC_014760 | 242181 | MBOVPG45_RS01035 | | | T>C |
| NC_014760 | 242183 | MBOVPG45_RS01035 | | | A>T |
| NC_014760 | 242197 | MBOVPG45_RS01035 | | | T>G |
| NC_014760 | 242209 | MBOVPG45_RS01035 | | | T>A |
| NC_014760 | 242630 | MBOVPG45_RS01035 | | | G>A |
| NC_014760 | 242831 | MBOVPG45_RS01040 | | | A>T |
| NC_014760 | 242888 | MBOVPG45_RS01040 | | | G>A |
| NC_014760 | 242916 | MBOVPG45_RS01040 | | | G>T |
| NC_014760 | 243092 | MBOVPG45_RS01040 | | - > ins T | |
| NC_014760 | 243287 | MBOVPG45_RS01040 | | | G>A |
| NC_014760 | 243294 | MBOVPG45_RS01040 | | | T>C |
| NC_014760 | 243333 | MBOVPG45_RS01040 | | | C>A |
| NC_014760 | 248926 | MBOVPG45_RS01075 | | | A>T |
| NC_014760 | 248930 | MBOVPG45_RS01075 | | | C>A |
| NC_014760 | 248942 | MBOVPG45_RS01075 | | | T>G |
| NC_014760 | 248946 | MBOVPG45_RS01075 | | | CG>TA |
| NC_014760 | 248951 | MBOVPG45_RS01075 | | | C>T |
| NC_014760 | 248962 | MBOVPG45_RS01075 | | | T>A |
| NC_014760 | 248988 | MBOVPG45_RS01075 | | | C>T |
| NC_014760 | 249000 | MBOVPG45_RS01075 | | | T>A |
| NC_014760 | 249123 | MBOVPG45_RS01075 | | | CA>TG |
| NC_014760 | 249129 | MBOVPG45_RS01075 | | | TAT>ATG |
| NC_014760 | 249346 | MBOVPG45_RS01075 | | | A>T |
| NC_014760 | 249348 | MBOVPG45_RS01075 | | | TG>CA |
| NC_014760 | 249351 | MBOVPG45_RS01075 | | | C>A |
| NC_014760 | 249354 | MBOVPG45_RS01075 | | | C>A |
| NC_014760 | 249367 | MBOVPG45_RS01075 | | | - > ins CTG |
| NC_014760 | 249434 | MBOVPG45_RS01075 | | | C>T |
| NC_014760 | 249489 | MBOVPG45_RS01075 | | | C>T |
| NC_014760 | 249606 | MBOVPG45_RS01075 | | | T>C |
| NC_014760 | 249610 | MBOVPG45_RS01075 | | | T>A |
| NC_014760 | 249612 | MBOVPG45_RS01075 | | | T>C |
| NC_014760 | 249614 | MBOVPG45_RS01075 | | | T>C |
| NC_014760 | 249637 | MBOVPG45_RS01075 | | | T>C |
| NC_014760 | 249690 | MBOVPG45_RS01075 | | | C>G |
| NC_014760 | 249731 | MBOVPG45_RS01075 | | | T>G |
| NC_014760 | 249827 | MBOVPG45_RS01075 | | | T>C |
| NC_014760 | 249892 | MBOVPG45_RS01075 | | T > del 1 | |
| NC_014760 | 249903 | MBOVPG45_RS01075 | | | T>A |
| NC_014760 | 249927 | MBOVPG45_RS01075 | | | T>A |
| NC_014760 | 249939 | MBOVPG45_RS01075 | | | T>C |
| NC_014760 | 249969 | MBOVPG45_RS01075 | | | T>G |
| NC_014760 | 250004 | MBOVPG45_RS01075 | | | C>T |
| NC_014760 | 250079 | MBOVPG45_RS01075 | | | G>T |
| NC_014760 | 250127 | MBOVPG45_RS01075 | | - > ins TT | G>A |

| | | | | | |
|---|---|---|---|---|---|
| NC_014760 | 252748 | MBOVPG45_RS01085 | | | C>T |
| NC_014760 | 253208 | MBOVPG45_RS01090 | | | A>G |
| NC_014760 | 256515 | pgk | | | A>T |
| NC_014760 | 256560 | pgk | | | T>C |
| NC_014760 | 257653 | MBOVPG45_RS01110 | | | C>A |
| NC_014760 | 257746 | MBOVPG45_RS01110 | | | A>G |
| NC_014760 | 264264 | MBOVPG45_RS01150 | | | T>C |
| NC_014760 | 264564 | MBOVPG45_RS01150 | | | C>T |
| NC_014760 | 264721 | MBOVPG45_RS01150 | | | G>C |
| NC_014760 | 264760 | MBOVPG45_RS01150 | | | C>T |
| NC_014760 | 264794 | MBOVPG45_RS01150 | | | T>C |
| NC_014760 | 264809 | MBOVPG45_RS01150 | | | T>G |
| NC_014760 | 265487 | MBOVPG45_RS01155 | | | A>G |
| NC_014760 | 265755 | MBOVPG45_RS01155 | | | C>A |
| NC_014760 | 265903 | MBOVPG45_RS01155 | | | T>A |
| NC_014760 | 265916 | MBOVPG45_RS01155 | | | C>G |
| NC_014760 | 265924 | MBOVPG45_RS01155 | | | G>T |
| NC_014760 | 265928 | MBOVPG45_RS01155 | | | C>T |
| NC_014760 | 266106 | MBOVPG45_RS01155 | | | G>A |
| NC_014760 | 266116 | MBOVPG45_RS01155 | | | A>T |
| NC_014760 | 266123 | MBOVPG45_RS01155 | | | C>T |
| NC_014760 | 266165 | MBOVPG45_RS01155 | | | - > ins AAA |
| NC_014760 | 266267 | MBOVPG45_RS01155 | | | C>T |
| NC_014760 | 266484 | MBOVPG45_RS01155 | | | G>C |
| NC_014760 | 266489 | MBOVPG45_RS01155 | | | AA>GT |
| NC_014760 | 266500 | MBOVPG45_RS01155 | | | T>A |
| NC_014760 | 266512 | MBOVPG45_RS01155 | | | A>T |
| NC_014760 | 266553 | MBOVPG45_RS01155 | | | A>G |
| NC_014760 | 266609 | MBOVPG45_RS01155 | | | T>A |
| NC_014760 | 266623 | MBOVPG45_RS01155 | | | G>T |
| NC_014760 | 266780 | MBOVPG45_RS01155 | | | C>T |
| NC_014760 | 266782 | MBOVPG45_RS01155 | | | A>T |
| NC_014760 | 266936 | MBOVPG45_RS01155 | | | T>A |
| NC_014760 | 266983 | MBOVPG45_RS01155 | | | G>T |
| NC_014760 | 267082 | MBOVPG45_RS01155 | | | TA>AG |
| NC_014760 | 267087 | MBOVPG45_RS01155 | | | A>G |
| NC_014760 | 267144 | MBOVPG45_RS01155 | | | A>T |
| NC_014760 | 267313 | MBOVPG45_RS01155 | | | GT>AC |
| NC_014760 | 267362 | MBOVPG45_RS01155 | | | T>A |
| NC_014760 | 267374 | MBOVPG45_RS01155 | | | A>T |
| NC_014760 | 267392 | MBOVPG45_RS01155 | | | A>G |
| NC_014760 | 267529 | MBOVPG45_RS01160 | | | AAG>GGA |
| NC_014760 | 267553 | MBOVPG45_RS01160 | | | G>A |
| NC_014760 | 267568 | MBOVPG45_RS01160 | | | G>T |
| NC_014760 | 267583 | MBOVPG45_RS01160 | | | T>A |
| NC_014760 | 267602 | MBOVPG45_RS01160 | | | T>G |
| NC_014760 | 267614 | MBOVPG45_RS01160 | | | CAAGTG > del 6 |

| | | | | | |
|---|---|---|---|---|---|
| NC_014760 | 267644 | MBOVPG45_RS01160 | | | A>G |
| NC_014760 | 267830 | MBOVPG45_RS01160 | | | C>G |
| NC_014760 | 267858 | MBOVPG45_RS01160 | | | A>T |
| NC_014760 | 267870 | MBOVPG45_RS01160 | | | CAC>TGG |
| NC_014760 | 267949 | MBOVPG45_RS01160 | | | C>T |
| NC_014760 | 268567 | MBOVPG45_RS01160 | | | A>G |
| NC_014760 | 268616 | MBOVPG45_RS01160 | | | T>C |
| NC_014760 | 268752 | MBOVPG45_RS01160 | | | A>T |
| NC_014760 | 268814 | MBOVPG45_RS01160 | | | C>A |
| NC_014760 | 268822 | MBOVPG45_RS01160 | | | A>G |
| NC_014760 | 269127 | MBOVPG45_RS01160 | | | A>T |
| NC_014760 | 269149 | MBOVPG45_RS01160 | | | G>A |
| NC_014760 | 269155 | MBOVPG45_RS01160 | | | C>T |
| NC_014760 | 269285 | MBOVPG45_RS01160 | | | C>T |
| NC_014760 | 269288 | MBOVPG45_RS01160 | | | T>A |
| NC_014760 | 269296 | MBOVPG45_RS01160 | | | G>A |
| NC_014760 | 269347 | MBOVPG45_RS01160 | | | A>G |
| NC_014760 | 269387 | MBOVPG45_RS01160 | | | T>C |
| NC_014760 | 269411 | MBOVPG45_RS01160 | | | C>T |
| NC_014760 | 269648 | MBOVPG45_RS01165 | | | A>G |
| NC_014760 | 269683 | MBOVPG45_RS01165 | | | G>A |
| NC_014760 | 269727 | MBOVPG45_RS01165 | | | A>G |
| NC_014760 | 269839 | MBOVPG45_RS01165 | | | G>A |
| NC_014760 | 271059 | MBOVPG45_RS01175 | | | C>T |
| NC_014760 | 271071 | MBOVPG45_RS01175 | | | T>C |
| NC_014760 | 271128 | MBOVPG45_RS01175 | | | A>G |
| NC_014760 | 271130 | MBOVPG45_RS01175 | | | AC>GT |
| NC_014760 | 271134 | MBOVPG45_RS01175 | | | CC>TT |
| NC_014760 | 271212 | MBOVPG45_RS01175 | | | C>T |
| NC_014760 | 271217 | MBOVPG45_RS01175 | | | C>T |
| NC_014760 | 271286 | MBOVPG45_RS01175 | | | T>C |
| NC_014760 | 271296 | MBOVPG45_RS01175 | | | T>A |
| NC_014760 | 271299 | MBOVPG45_RS01175 | | | T>C |
| NC_014760 | 271302 | MBOVPG45_RS01175 | | | C>T |
| NC_014760 | 271328 | MBOVPG45_RS01175 | | | C>T |
| NC_014760 | 271341 | MBOVPG45_RS01175 | | | A>C |
| NC_014760 | 271353 | MBOVPG45_RS01175 | | | G>A |
| NC_014760 | 271359 | MBOVPG45_RS01175 | | | T>C |
| NC_014760 | 271367 | MBOVPG45_RS01175 | | | TT>AC |
| NC_014760 | 271428 | MBOVPG45_RS01175 | | | G>C |
| NC_014760 | 271430 | MBOVPG45_RS01175 | | | AC>GT |
| NC_014760 | 271481 | MBOVPG45_RS01175 | | | C>T |
| NC_014760 | 271504 | MBOVPG45_RS01175 | | | - > ins A |
| NC_014760 | 271513 | MBOVPG45_RS01175 | | | A>C |
| NC_014760 | 273077 | MBOVPG45_RS01185 | | | A>T |
| NC_014760 | 273150 | MBOVPG45_RS01185 | | | G>A |
| NC_014760 | 273189 | MBOVPG45_RS01185 | | | - > ins A |

| | | | | | |
|---|---|---|---|---|---|
| NC_014760 | 273560 | MBOVPG45_RS01185 | | | A>G |
| NC_014760 | 273673 | MBOVPG45_RS01185 | | | T>A |
| NC_014760 | 273698 | MBOVPG45_RS01185 | | | G>A |
| NC_014760 | 273974 | MBOVPG45_RS01185 | | | G>A |
| NC_014760 | 274146 | MBOVPG45_RS01185 | | | T>C |
| NC_014760 | 274167 | MBOVPG45_RS01185 | | | A>G |
| NC_014760 | 274254 | MBOVPG45_RS01185 | | | G>A |
| NC_014760 | 274257 | MBOVPG45_RS01185 | | | A>G |
| NC_014760 | 274264 | MBOVPG45_RS01185 | | | A>T |
| NC_014760 | 274268 | MBOVPG45_RS01185 | | | G>A |
| NC_014760 | 274273 | MBOVPG45_RS01185 | | | CA>TT |
| NC_014760 | 274286 | MBOVPG45_RS01185 | | | G>A |
| NC_014760 | 274325 | MBOVPG45_RS01185 | | | A>T |
| NC_014760 | 274468 | MBOVPG45_RS01185 | | | A>T |
| NC_014760 | 274497 | MBOVPG45_RS01185 | | | G>A |
| NC_014760 | 274644 | MBOVPG45_RS01185 | | | A>T |
| NC_014760 | 274700 | MBOVPG45_RS01185 | | | G>A |
| NC_014760 | 274776 | MBOVPG45_RS01185 | | | C>T |
| NC_014760 | 274853 | MBOVPG45_RS01185 | | | CAA>TCT |
| NC_014760 | 274908 | MBOVPG45_RS01185 | | | A>C |
| NC_014760 | 275207 | MBOVPG45_RS01185 | | | T>C |
| NC_014760 | 275216 | MBOVPG45_RS01185 | | | A>G |
| NC_014760 | 275280 | MBOVPG45_RS01185 | | | A>G |
| NC_014760 | 275322 | MBOVPG45_RS01185 | | | A>C |
| NC_014760 | 275327 | MBOVPG45_RS01185 | | | A>G |
| NC_014760 | 275391 | MBOVPG45_RS01185 | | | C>T |
| NC_014760 | 275486 | MBOVPG45_RS01185 | | | AG>GA |
| NC_014760 | 275504 | MBOVPG45_RS01185 | | | A>G |
| NC_014760 | 275538 | MBOVPG45_RS01185 | | | A>G |
| NC_014760 | 275567 | MBOVPG45_RS01185 | | | A>G |
| NC_014760 | 275617 | MBOVPG45_RS04625 | | | A>G |
| NC_014760 | 275643 | MBOVPG45_RS04625 | | G>A | |
| NC_014760 | 275691 | MBOVPG45_RS04625 | | | G>A |
| NC_014760 | 275940 | MBOVPG45_RS04625 | | | G>T |
| NC_014760 | 276019 | MBOVPG45_RS04625 | | | C>T |
| NC_014760 | 276189 | MBOVPG45_RS04625 | | | A>G |
| NC_014760 | 276761 | MBOVPG45_RS01195 | | | G>C |
| NC_014760 | 276765 | MBOVPG45_RS01195 | | | T>A |
| NC_014760 | 276769 | MBOVPG45_RS01195 | | | C>A |
| NC_014760 | 276810 | MBOVPG45_RS01195 | | | TT>GC |
| NC_014760 | 276890 | MBOVPG45_RS01195 | | | G>A |
| NC_014760 | 276924 | MBOVPG45_RS01195 | | | T>A |
| NC_014760 | 277137 | MBOVPG45_RS01195 | | | C>T |
| NC_014760 | 277470 | MBOVPG45_RS01195 | | | A>G |
| NC_014760 | 277571 | MBOVPG45_RS01195 | | | A>G |
| NC_014760 | 277633 | MBOVPG45_RS01195 | | | A>T |
| NC_014760 | 278565 | MBOVPG45_RS01195 | | | A>G |

| | | | | | |
|---|---|---|---|---|---|
| NC_014760 | 279751 | MBOVPG45_RS01195 | | | A > del 1 |
| NC_014760 | 280080 | MBOVPG45_RS01200 | | | T>C |
| NC_014760 | 281463 | MBOVPG45_RS01205 | | | C>T |
| NC_014760 | 281610 | MBOVPG45_RS01205 | | | G>A |
| NC_014760 | 281632 | MBOVPG45_RS01205 | | | GG>CA |
| NC_014760 | 281786 | MBOVPG45_RS01205 | | | T>C |
| NC_014760 | 281897 | MBOVPG45_RS01205 | | | A>C |
| NC_014760 | 282020 | MBOVPG45_RS01205 | | | C>T |
| NC_014760 | 282122 | MBOVPG45_RS01205 | | | G>A |
| NC_014760 | 282460 | MBOVPG45_RS01210 | | | G>A |
| NC_014760 | 282463 | MBOVPG45_RS01210 | | | G>C |
| NC_014760 | 283862 | MBOVPG45_RS01215 | | | T>C |
| NC_014760 | 284090 | MBOVPG45_RS01215 | | | A>G |
| NC_014760 | 284185 | MBOVPG45_RS01215 | | | A>T |
| NC_014760 | 284324 | MBOVPG45_RS01215 | | | A>G |
| NC_014760 | 284693 | MBOVPG45_RS01215 | | | C>T |
| NC_014760 | 285240 | MBOVPG45_RS01215 | | | C>T |
| NC_014760 | 285764 | MBOVPG45_RS01215 | | | C>T |
| NC_014760 | 286046 | MBOVPG45_RS01215 | | | T>C |
| NC_014760 | 286089 | MBOVPG45_RS01215 | | | C>T |
| NC_014760 | 286167 | MBOVPG45_RS01215 | | | A>C |
| NC_014760 | 286175 | MBOVPG45_RS01215 | | | A>C |
| NC_014760 | 286340 | MBOVPG45_RS01215 | | | C>T |
| NC_014760 | 286434 | MBOVPG45_RS01215 | | | A>T |
| NC_014760 | 286439 | MBOVPG45_RS01215 | | | C>T |
| NC_014760 | 287082 | MBOVPG45_RS01215 | | | G>A |
| NC_014760 | 287154 | MBOVPG45_RS01215 | | | T>C |
| NC_014760 | 287203 | MBOVPG45_RS01215 | | | GA>AG |
| NC_014760 | 287221 | MBOVPG45_RS01215 | | | C>A |
| NC_014760 | 287282 | MBOVPG45_RS01215 | | | T>C |
| NC_014760 | 287454 | MBOVPG45_RS01215 | | | C>T |
| NC_014760 | 287766 | MBOVPG45_RS01215 | | | T>C |
| NC_014760 | 288107 | MBOVPG45_RS01215 | | | T>C |
| NC_014760 | 288159 | MBOVPG45_RS01215 | | | T>G |
| NC_014760 | 288176 | MBOVPG45_RS01215 | | | T>C |
| NC_014760 | 288194 | MBOVPG45_RS01215 | | | C>T |
| NC_014760 | 288398 | MBOVPG45_RS01215 | | | T>C |
| NC_014760 | 289078 | gyrA | | | A>G |
| NC_014760 | 289477 | gyrA | | | C>T |
| NC_014760 | 290091 | gyrA | | | G>A |
| NC_014760 | 290265 | gyrA | | | A>G |
| NC_014760 | 290293 | gyrA | | | AG>GA |
| NC_014760 | 290460 | gyrA | | | G>A |
| NC_014760 | 290552 | gyrA | | | A>T |
| NC_014760 | 290620 | gyrA | | | C>T |
| NC_014760 | 291937 | MBOVPG45_RS01225 | | | A>T |
| NC_014760 | 292121 | MBOVPG45_RS01225 | | | G>C |

| | | | | | |
|---|---|---|---|---|---|
| NC_014760 | 292503 | MBOVPG45_RS01225 | | | A>G |
| NC_014760 | 292515 | MBOVPG45_RS01225 | | | C>A |
| NC_014760 | 293006 | MBOVPG45_RS01225 | | | A>T |
| NC_014760 | 293078 | MBOVPG45_RS01225 | | | C>A |
| NC_014760 | 293219 | MBOVPG45_RS01225 | | | G>A |
| NC_014760 | 293726 | rpsD | | | G>A |
| NC_014760 | 294141 | rpsD | | | T>C |
| NC_014760 | 294448 | rpmE | | | A>G |
| NC_014760 | 294901 | hrcA | | | C>T |
| NC_014760 | 295867 | grpE | | | G>C |
| NC_014760 | 295952 | grpE | | | C>T |
| NC_014760 | 295974 | grpE | | | T>A |
| NC_014760 | 296555 | grpE | | | T>C |
| NC_014760 | 298696 | MBOVPG45_RS01270 | rRNA pseudouridine | | C>T |
| NC_014760 | 299077 | MBOVPG45_RS01270 | rRNA pseudouridine | | G>A |
| NC_014760 | 299555 | MBOVPG45_RS01275 | | | G>A |
| NC_014760 | 299714 | MBOVPG45_RS01275 | | | C>T |
| NC_014760 | 300028 | MBOVPG45_RS01275 | | | A>T |
| NC_014760 | 300225 | MBOVPG45_RS01275 | | | G>A |
| NC_014760 | 300470 | MBOVPG45_RS01275 | | | C>G |
| NC_014760 | 300473 | MBOVPG45_RS01275 | | | A>G |
| NC_014760 | 302153 | recA | | | TA>CG |
| NC_014760 | 302392 | recA | | | T>C |
| NC_014760 | 303010 | recU | | | T>A |
| NC_014760 | 304123 | rpsJ | | C>T | |
| NC_014760 | 305116 | rplC | | | C>T |
| NC_014760 | 305204 | rplD | | | T>A |
| NC_014760 | 305279 | rplD | | | A>G |
| NC_014760 | 305303 | rplD | | | G>A |
| NC_014760 | 305358 | rplD | | | T>C |
| NC_014760 | 305360 | rplD | | | G>A |
| NC_014760 | 305376 | rplD | | | A>C |
| NC_014760 | 305408 | rplD | | | G>A |
| NC_014760 | 305453 | rplD | | | G>A |
| NC_014760 | 305460 | rplD | | | - > ins AGCTAAAGA |
| NC_014760 | 305704 | rplD | | | AG>GA |
| NC_014760 | 306754 | rplB | | | C>A |
| NC_014760 | 307518 | rpsS | | | T>C |
| NC_014760 | 308038 | rplV | | | A>T |
| NC_014760 | 308343 | rpsC | | | TA>CG |
| NC_014760 | 308344 | rpsC | | A>G | |
| NC_014760 | 308498 | rpsC | | G>A | |
| NC_014760 | 310225 | MBOVPG45_RS01360 | | | A>G |
| NC_014760 | 311251 | rpsH | | | A>G |

| | | | | | |
|---|---|---|---|---|---|
| NC_014760 | 312554 | rpsE | | | G>A |
| NC_014760 | 313031 | rpsE | | | C>T |
| NC_014760 | 313084 | rpsE | | C>A | |
| NC_014760 | 313704 | MBOVPG45_RS01400 | | | A>T |
| NC_014760 | 313732 | MBOVPG45_RS01400 | | | A>G |
| NC_014760 | 313826 | MBOVPG45_RS01400 | | | C>A |
| NC_014760 | 313860 | MBOVPG45_RS01400 | | | A>G |
| NC_014760 | 313862 | MBOVPG45_RS01400 | | | T>A |
| NC_014760 | 313937 | MBOVPG45_RS01400 | | | T>A |
| NC_014760 | 314031 | MBOVPG45_RS01400 | | | A>G |
| NC_014760 | 314040 | MBOVPG45_RS01400 | | | G > del 1 |
| NC_014760 | 314346 | MBOVPG45_RS01400 | | | AC>GT |
| NC_014760 | 314538 | MBOVPG45_RS01400 | | | T > del 1 |
| NC_014760 | 314552 | MBOVPG45_RS01400 | | | TGAAAATAATA > del 11 |
| NC_014760 | 314670 | MBOVPG45_RS01400 | | | A>G |
| NC_014760 | 314743 | MBOVPG45_RS01400 | | | TA>CT |
| NC_014760 | 314969 | MBOVPG45_RS01405 | | | T>C |
| NC_014760 | 315035 | MBOVPG45_RS01405 | | | T>C |
| NC_014760 | 315073 | MBOVPG45_RS01405 | | | T>A |
| NC_014760 | 315435 | MBOVPG45_RS01405 | | | G>A |
| NC_014760 | 315484 | MBOVPG45_RS01405 | | | C>G |
| NC_014760 | 315494 | MBOVPG45_RS01405 | | | T>C |
| NC_014760 | 315760 | MBOVPG45_RS01405 | | | C>T |
| NC_014760 | 333960 | MBOVPG45_RS01485 | | | A>T |
| NC_014760 | 334353 | MBOVPG45_RS01485 | | | A>G |
| NC_014760 | 334428 | MBOVPG45_RS01490 | | | T>C |
| NC_014760 | 334526 | MBOVPG45_RS01490 | | | TT>CC |
| NC_014760 | 334561 | MBOVPG45_RS01490 | | | T>A |
| NC_014760 | 334564 | MBOVPG45_RS01490 | | | C>T |
| NC_014760 | 334577 | MBOVPG45_RS01490 | | | G>C |
| NC_014760 | 334660 | MBOVPG45_RS01490 | | | C>T |
| NC_014760 | 334669 | MBOVPG45_RS01490 | | | G>C |
| NC_014760 | 334702 | MBOVPG45_RS01490 | | | T>C |
| NC_014760 | 334761 | MBOVPG45_RS01490 | | | C>T |
| NC_014760 | 334843 | MBOVPG45_RS01490 | | | T>C |
| NC_014760 | 334882 | MBOVPG45_RS01490 | | | C>T |
| NC_014760 | 335077 | MBOVPG45_RS01490 | | | C>T |
| NC_014760 | 335091 | MBOVPG45_RS01490 | | | T>G |
| NC_014760 | 335141 | MBOVPG45_RS01490 | | | A>T |
| NC_014760 | 335180 | MBOVPG45_RS01490 | | | T>C |
| NC_014760 | 335239 | MBOVPG45_RS01490 | | | C>T |
| NC_014760 | 335389 | MBOVPG45_RS01490 | | | T>C |
| NC_014760 | 335446 | MBOVPG45_RS01490 | | | T>G |
| NC_014760 | 335706 | tpiA | | | C>T |
| NC_014760 | 336212 | tpiA | | | T>A |
| NC_014760 | 336442 | deoC | | | T>C |

| | | | | | |
|---|---|---|---|---|---|
| NC_014760 | 336667 | deoC | | | C>T |
| NC_014760 | 336727 | deoC | | | C>T |
| NC_014760 | 336853 | deoC | | | A>T |
| NC_014760 | 337141 | MBOVPG45_RS01505 | | | CC>TT |
| NC_014760 | 337535 | MBOVPG45_RS01505 | | | C>A |
| NC_014760 | 338666 | deoD | | | C>T |
| NC_014760 | 338748 | deoD | | | T>A |
| NC_014760 | 338938 | deoD | | | T>C |
| NC_014760 | 339998 | MBOVPG45_RS01515 | | | C>A |
| NC_014760 | 340222 | MBOVPG45_RS01515 | | | A>T |
| NC_014760 | 341084 | obgE | | | G>A |
| NC_014760 | 341199 | obgE | | | T>A |
| NC_014760 | 341776 | MBOVPG45_RS01525 | | | T>C |
| NC_014760 | 341780 | MBOVPG45_RS01525 | | | C>A |
| NC_014760 | 341787 | MBOVPG45_RS01525 | | | T>G |
| NC_014760 | 341790 | MBOVPG45_RS01525 | | | T>G |
| NC_014760 | 341812 | MBOVPG45_RS01525 | | | T>C |
| NC_014760 | 341832 | MBOVPG45_RS01525 | | | C>T |
| NC_014760 | 341839 | MBOVPG45_RS01525 | | | C>T |
| NC_014760 | 341869 | MBOVPG45_RS01525 | | | T>C |
| NC_014760 | 343030 | MBOVPG45_RS01525 | | | A>G |
| NC_014760 | 345146 | MBOVPG45_RS01540 | | | A>G |
| NC_014760 | 345580 | MBOVPG45_RS01545 | | | C>T |
| NC_014760 | 345591 | MBOVPG45_RS01545 | | | T>A |
| NC_014760 | 345634 | MBOVPG45_RS01545 | | | A>G |
| NC_014760 | 345777 | MBOVPG45_RS01545 | | | T>G |
| NC_014760 | 347859 | MBOVPG45_RS01550 | | | C>T |
| NC_014760 | 348078 | MBOVPG45_RS01550 | | | C>T |
| NC_014760 | 348474 | MBOVPG45_RS01550 | | | C>T |
| NC_014760 | 348617 | MBOVPG45_RS01550 | | | T>C |
| NC_014760 | 348750 | MBOVPG45_RS01550 | | | G>T |
| NC_014760 | 348758 | MBOVPG45_RS01550 | | | T>C |
| NC_014760 | 349035 | MBOVPG45_RS01555 | | | G>T |
| NC_014760 | 349063 | MBOVPG45_RS01555 | | | G>T |
| NC_014760 | 349169 | MBOVPG45_RS01555 | | | C>T |
| NC_014760 | 349178 | MBOVPG45_RS01555 | | | C>T |
| NC_014760 | 349292 | MBOVPG45_RS01555 | | | T>C |
| NC_014760 | 349339 | MBOVPG45_RS01555 | | | TG>GA |
| NC_014760 | 349399 | MBOVPG45_RS01555 | | | T>G |
| NC_014760 | 349419 | MBOVPG45_RS01555 | | | A>C |
| NC_014760 | 349421 | MBOVPG45_RS01555 | | | T>C |
| NC_014760 | 349516 | MBOVPG45_RS01555 | | | C>T |
| NC_014760 | 349522 | MBOVPG45_RS01555 | | | G>A |
| NC_014760 | 349526 | MBOVPG45_RS01555 | | | C>T |
| NC_014760 | 349541 | MBOVPG45_RS01555 | | | C>G |
| NC_014760 | 349572 | MBOVPG45_RS01555 | | | G>T |
| NC_014760 | 349625 | MBOVPG45_RS01555 | | | A>C |

| | | | | | |
|---|---|---|---|---|---|
| NC_014760 | 349705 | MBOVPG45_RS01555 | | | A>G |
| NC_014760 | 349757 | MBOVPG45_RS01555 | | | G>A |
| NC_014760 | 349781 | MBOVPG45_RS01555 | | T>C | |
| NC_014760 | 349785 | MBOVPG45_RS01555 | | | - > ins TTA |
| NC_014760 | 349829 | MBOVPG45_RS01555 | | | C>T |
| NC_014760 | 349863 | MBOVPG45_RS01555 | | | T>A |
| NC_014760 | 349865 | MBOVPG45_RS01555 | | | T>C |
| NC_014760 | 349871 | MBOVPG45_RS01555 | | | GA>AG |
| NC_014760 | 349962 | MBOVPG45_RS01555 | | | T>A |
| NC_014760 | 349984 | MBOVPG45_RS01555 | | | AT>GC |
| NC_014760 | 349992 | MBOVPG45_RS01555 | | | G>T |
| NC_014760 | 350199 | MBOVPG45_RS01555 | | | CT>TG |
| NC_014760 | 350256 | MBOVPG45_RS01555 | | | C>A |
| NC_014760 | 350432 | MBOVPG45_RS01555 | | | C>T |
| NC_014760 | 350435 | MBOVPG45_RS01555 | | | T>C |
| NC_014760 | 350564 | MBOVPG45_RS01555 | | | G>T |
| NC_014760 | 351154 | MBOVPG45_RS01560 | | | T>C |
| NC_014760 | 351197 | MBOVPG45_RS01560 | | | T>C |
| NC_014760 | 351279 | MBOVPG45_RS01560 | | | A>T |
| NC_014760 | 351910 | MBOVPG45_RS01560 | | | A>T |
| NC_014760 | 352136 | MBOVPG45_RS01560 | | | C>T |
| NC_014760 | 352974 | MBOVPG45_RS01565 | | | A>C |
| NC_014760 | 353206 | MBOVPG45_RS01565 | | | A>G |
| NC_014760 | 353654 | MBOVPG45_RS01570 | | | TT>GA |
| NC_014760 | 353787 | MBOVPG45_RS01570 | | | T>C |
| NC_014760 | 354028 | MBOVPG45_RS01570 | | | A>T |
| NC_014760 | 354038 | MBOVPG45_RS01570 | | | T>C |
| NC_014760 | 354232 | MBOVPG45_RS01570 | | | T>G |
| NC_014760 | 354270 | MBOVPG45_RS01570 | | | C>G |
| NC_014760 | 356763 | MBOVPG45_RS01580 | | | A>G |
| NC_014760 | 357311 | MBOVPG45_RS01585 | | | A>G |
| NC_014760 | 357983 | MBOVPG45_RS01585 | | | C>T |
| NC_014760 | 358227 | MBOVPG45_RS01590 | | | T>C |
| NC_014760 | 358379 | MBOVPG45_RS01590 | | | T>C |
| NC_014760 | 358613 | MBOVPG45_RS01590 | | | G>T |
| NC_014760 | 358731 | MBOVPG45_RS01590 | | A>G | |
| NC_014760 | 358749 | MBOVPG45_RS01590 | | | T>A |
| NC_014760 | 359126 | MBOVPG45_RS01595 | | | A>G |
| NC_014760 | 361083 | MBOVPG45_RS01600 | | | C>A |
| NC_014760 | 361279 | MBOVPG45_RS01600 | | | A>G |
| NC_014760 | 361294 | MBOVPG45_RS01600 | | | - > ins TAT |
| NC_014760 | 361298 | MBOVPG45_RS01600 | | | T>C |
| NC_014760 | 361391 | MBOVPG45_RS01600 | | | C>T |
| NC_014760 | 361394 | MBOVPG45_RS01600 | | | TG>CA |
| NC_014760 | 361399 | MBOVPG45_RS01600 | | | C>T |
| NC_014760 | 361431 | MBOVPG45_RS01600 | | | T>C |
| NC_014760 | 361591 | MBOVPG45_RS01600 | | | T>A |

| | | | | | |
|---|---|---|---|---|---|
| NC_014760 | 361594 | MBOVPG45_RS01600 | | | T>C |
| NC_014760 | 361600 | MBOVPG45_RS01600 | | | T>A |
| NC_014760 | 361877 | MBOVPG45_RS01600 | | | - > ins TTCACT |
| NC_014760 | 362024 | MBOVPG45_RS01600 | | | C>T |
| NC_014760 | 362026 | MBOVPG45_RS01600 | | | C>T |
| NC_014760 | 362101 | MBOVPG45_RS01600 | | | T>C |
| NC_014760 | 362107 | MBOVPG45_RS01600 | | | CT>TC |
| NC_014760 | 362123 | MBOVPG45_RS01600 | | | C>T |
| NC_014760 | 362149 | MBOVPG45_RS01600 | | | T>C |
| NC_014760 | 362159 | MBOVPG45_RS01600 | | | A>T |
| NC_014760 | 362243 | MBOVPG45_RS01600 | | | A>T |
| NC_014760 | 362246 | MBOVPG45_RS01600 | | | C>T |
| NC_014760 | 362262 | MBOVPG45_RS01600 | | | T>G |
| NC_014760 | 362457 | MBOVPG45_RS01600 | | | A>T |
| NC_014760 | 362785 | MBOVPG45_RS01600 | | | T>A |
| NC_014760 | 362788 | MBOVPG45_RS01600 | | | GA>TT |
| NC_014760 | 362798 | MBOVPG45_RS01600 | | | T>C |
| NC_014760 | 362875 | MBOVPG45_RS01600 | | | G>T |
| NC_014760 | 362981 | MBOVPG45_RS01600 | | | C>T |
| NC_014760 | 363231 | MBOVPG45_RS01605 | | | T>A |
| NC_014760 | 363262 | MBOVPG45_RS01605 | | | C>T |
| NC_014760 | 363424 | MBOVPG45_RS01605 | | | C>T |
| NC_014760 | 363477 | MBOVPG45_RS01605 | | | G>A |
| NC_014760 | 363482 | MBOVPG45_RS01605 | | | T>A |
| NC_014760 | 364058 | MBOVPG45_RS01610 | | | A>T |
| NC_014760 | 364134 | MBOVPG45_RS01610 | | | C>T |
| NC_014760 | 364679 | MBOVPG45_RS01610 | | | C>T |
| NC_014760 | 365021 | MBOVPG45_RS01610 | | | C>T |
| NC_014760 | 365106 | MBOVPG45_RS01610 | | | G>T |
| NC_014760 | 365190 | MBOVPG45_RS01610 | | | T>C |
| NC_014760 | 365312 | MBOVPG45_RS01615 | | | T>C |
| NC_014760 | 365314 | MBOVPG45_RS01615 | | | T>C |
| NC_014760 | 365416 | MBOVPG45_RS01615 | | | T>C |
| NC_014760 | 365529 | MBOVPG45_RS01615 | | | G>A |
| NC_014760 | 365644 | MBOVPG45_RS01615 | | | TT>CA |
| NC_014760 | 365670 | MBOVPG45_RS01615 | | | T>C |
| NC_014760 | 365944 | MBOVPG45_RS01615 | | | T>C |
| NC_014760 | 366042 | MBOVPG45_RS01615 | | | T>C |
| NC_014760 | 369370 | MBOVPG45_RS01630 | | | C>T |
| NC_014760 | 369380 | MBOVPG45_RS01630 | | | A>T |
| NC_014760 | 369441 | MBOVPG45_RS01630 | | | G>A |
| NC_014760 | 370053 | MBOVPG45_RS01635 | | | AT>TG |
| NC_014760 | 370073 | MBOVPG45_RS01635 | | | GCA>AGC |
| NC_014760 | 370107 | MBOVPG45_RS01635 | | | T>A |
| NC_014760 | 370779 | MBOVPG45_RS01635 | | | T>C |
| NC_014760 | 371209 | MBOVPG45_RS01640 | | | G>C |
| NC_014760 | 371242 | MBOVPG45_RS01640 | | | T>A |

| | | | | | |
|---|---|---|---|---|---|
| NC_014760 | 371577 | MBOVPG45_RS01640 | | | T>C |
| NC_014760 | 371649 | MBOVPG45_RS01640 | | | G>C |
| NC_014760 | 372376 | MBOVPG45_RS01640 | | | G>A |
| NC_014760 | 373493 | MBOVPG45_RS01645 | | | C>G |
| NC_014760 | 373586 | MBOVPG45_RS01645 | | | T>C |
| NC_014760 | 373874 | MBOVPG45_RS01645 | | | T>C |
| NC_014760 | 373922 | MBOVPG45_RS01645 | | | G>A |
| NC_014760 | 374153 | MBOVPG45_RS01645 | | | C>A |
| NC_014760 | 374177 | MBOVPG45_RS01645 | | | G>A |
| NC_014760 | 374281 | MBOVPG45_RS01645 | | | C>T |
| NC_014760 | 374344 | MBOVPG45_RS01650 | | | T>C |
| NC_014760 | 374387 | MBOVPG45_RS01650 | | | T>C |
| NC_014760 | 374469 | MBOVPG45_RS01650 | | | A>T |
| NC_014760 | 375084 | MBOVPG45_RS01650 | | | C>A |
| NC_014760 | 375100 | MBOVPG45_RS01650 | | | A>T |
| NC_014760 | 375326 | MBOVPG45_RS01650 | | | C>T |
| NC_014760 | 376115 | MBOVPG45_RS01655 | | | C>T |
| NC_014760 | 376551 | MBOVPG45_RS01655 | | | C>T |
| NC_014760 | 376633 | MBOVPG45_RS01655 | | | C>T |
| NC_014760 | 376680 | MBOVPG45_RS01655 | | | T>C |
| NC_014760 | 376692 | MBOVPG45_RS01655 | | | AT>GC |
| NC_014760 | 377305 | MBOVPG45_RS01660 | | | C>T |
| NC_014760 | 377784 | MBOVPG45_RS01665 | | | T>C |
| NC_014760 | 377815 | MBOVPG45_RS01665 | | | T>C |
| NC_014760 | 377817 | MBOVPG45_RS01665 | | | T>C |
| NC_014760 | 377825 | MBOVPG45_RS01665 | | | G>A |
| NC_014760 | 377963 | MBOVPG45_RS01665 | | | G>T |
| NC_014760 | 377967 | MBOVPG45_RS01665 | | | T>C |
| NC_014760 | 377979 | MBOVPG45_RS01665 | | | T>C |
| NC_014760 | 378171 | MBOVPG45_RS01665 | | | C>T |
| NC_014760 | 378220 | MBOVPG45_RS01665 | | | C>G |
| NC_014760 | 378314 | MBOVPG45_RS01665 | | | T>C |
| NC_014760 | 378595 | MBOVPG45_RS01665 | | | A>T |
| NC_014760 | 378597 | MBOVPG45_RS01665 | | | C>G |
| NC_014760 | 378604 | MBOVPG45_RS01665 | | | C>A |
| NC_014760 | 378644 | MBOVPG45_RS01665 | | | A>G |
| NC_014760 | 378750 | MBOVPG45_RS01665 | | | C>T |
| NC_014760 | 378791 | MBOVPG45_RS01665 | | | C>G |
| NC_014760 | 378807 | MBOVPG45_RS01665 | | | G>T |
| NC_014760 | 379068 | MBOVPG45_RS01665 | | | T>C |
| NC_014760 | 380382 | lepA | | | C>T |
| NC_014760 | 380385 | lepA | | | C>G |
| NC_014760 | 380396 | lepA | | | G>T |
| NC_014760 | 380514 | lepA | | | G>A |
| NC_014760 | 381462 | MBOVPG45_RS01675 | | | C>A |
| NC_014760 | 381595 | MBOVPG45_RS01675 | | | G>A |
| NC_014760 | 392689 | MBOVPG45_RS01720 | | | G>A |

| | | | | | |
|---|---|---|---|---|---|
| NC_014760 | 397770 | MBOVPG45_RS01750 | | | G>A |
| NC_014760 | 397791 | MBOVPG45_RS01750 | | | C>G |
| NC_014760 | 397884 | MBOVPG45_RS01750 | | | G>T |
| NC_014760 | 398022 | MBOVPG45_RS01750 | | | CA>TG |
| NC_014760 | 398025 | MBOVPG45_RS01750 | | | T>A |
| NC_014760 | 398036 | MBOVPG45_RS01750 | | | C>A |
| NC_014760 | 398393 | MBOVPG45_RS01755 | | | TG>CT |
| NC_014760 | 398400 | MBOVPG45_RS01755 | | | C>T |
| NC_014760 | 398404 | MBOVPG45_RS01755 | | | A>T |
| NC_014760 | 398406 | MBOVPG45_RS01755 | | | G>C |
| NC_014760 | 398409 | MBOVPG45_RS01755 | | | G>A |
| NC_014760 | 398412 | MBOVPG45_RS01755 | | | C>T |
| NC_014760 | 398426 | MBOVPG45_RS01755 | | | G>A |
| NC_014760 | 398489 | MBOVPG45_RS01755 | | | A>T |
| NC_014760 | 398533 | MBOVPG45_RS01755 | | | T>A |
| NC_014760 | 398538 | MBOVPG45_RS01755 | | | C>T |
| NC_014760 | 398623 | MBOVPG45_RS01755 | | | GG>AC |
| NC_014760 | 398713 | MBOVPG45_RS01755 | | | A>C |
| NC_014760 | 398726 | MBOVPG45_RS01755 | | | C>T |
| NC_014760 | 398744 | MBOVPG45_RS01755 | | | T>C |
| NC_014760 | 398769 | MBOVPG45_RS01755 | | | C>T |
| NC_014760 | 398831 | MBOVPG45_RS01755 | | | T>A |
| NC_014760 | 398835 | MBOVPG45_RS01755 | | | C>T |
| NC_014760 | 398850 | MBOVPG45_RS01755 | | | T>C |
| NC_014760 | 398855 | MBOVPG45_RS01755 | | | T>C |
| NC_014760 | 398861 | MBOVPG45_RS01755 | | | T>A |
| NC_014760 | 398865 | MBOVPG45_RS01755 | | | C>T |
| NC_014760 | 398876 | MBOVPG45_RS01755 | | | G>C |
| NC_014760 | 398886 | MBOVPG45_RS01755 | | | T>C |
| NC_014760 | 399083 | MBOVPG45_RS01755 | | | G>A |
| NC_014760 | 399090 | MBOVPG45_RS01755 | | | C>T |
| NC_014760 | 399131 | MBOVPG45_RS01755 | | | A>G |
| NC_014760 | 399162 | MBOVPG45_RS01755 | | | C>T |
| NC_014760 | 399190 | MBOVPG45_RS01760 | | | T>C |
| NC_014760 | 399195 | MBOVPG45_RS01760 | | | C>T |
| NC_014760 | 399237 | MBOVPG45_RS01760 | | | C>T |
| NC_014760 | 399241 | MBOVPG45_RS01760 | | | CA>TG |
| NC_014760 | 399253 | MBOVPG45_RS01760 | | | T>C |
| NC_014760 | 399264 | MBOVPG45_RS01760 | | | A>T |
| NC_014760 | 399343 | MBOVPG45_RS01760 | | | T>G |
| NC_014760 | 399379 | MBOVPG45_RS01760 | | | T>C |
| NC_014760 | 399388 | MBOVPG45_RS01760 | | | C>T |
| NC_014760 | 399392 | MBOVPG45_RS01760 | | | G>T |
| NC_014760 | 399394 | MBOVPG45_RS01760 | | | C>G |
| NC_014760 | 399549 | MBOVPG45_RS01760 | | | C>G |
| NC_014760 | 399585 | MBOVPG45_RS01760 | | | T>C |
| NC_014760 | 399591 | MBOVPG45_RS01760 | | | TT>GG |

| | | | | | |
|---|---|---|---|---|---|
| NC_014760 | 399688 | MBOVPG45_RS01760 | | | C>T |
| NC_014760 | 399695 | MBOVPG45_RS01760 | | | T>G |
| NC_014760 | 399705 | MBOVPG45_RS01760 | | | A>T |
| NC_014760 | 399717 | MBOVPG45_RS01760 | | | T>C |
| NC_014760 | 399726 | MBOVPG45_RS01760 | | | G>A |
| NC_014760 | 399735 | MBOVPG45_RS01760 | | | TC>CT |
| NC_014760 | 399743 | MBOVPG45_RS01760 | | | C>A |
| NC_014760 | 399808 | MBOVPG45_RS01760 | | | T>C |
| NC_014760 | 400121 | MBOVPG45_RS01765 | | | C>T |
| NC_014760 | 400128 | MBOVPG45_RS01765 | | | A>T |
| NC_014760 | 400138 | MBOVPG45_RS01765 | | | T>A |
| NC_014760 | 400142 | MBOVPG45_RS01765 | | | G>C |
| NC_014760 | 400154 | MBOVPG45_RS01765 | | | GTG>AAT |
| NC_014760 | 400169 | MBOVPG45_RS01765 | | | C>T |
| NC_014760 | 400208 | MBOVPG45_RS01765 | | | C>T |
| NC_014760 | 400223 | MBOVPG45_RS01765 | | | GA>AG |
| NC_014760 | 400349 | MBOVPG45_RS01765 | | | TT>CA |
| NC_014760 | 400370 | MBOVPG45_RS01765 | | | C>T |
| NC_014760 | 400405 | MBOVPG45_RS01765 | | | T>C |
| NC_014760 | 400484 | MBOVPG45_RS01765 | | | T>C |
| NC_014760 | 400577 | MBOVPG45_RS01765 | | | C>T |
| NC_014760 | 400580 | MBOVPG45_RS01765 | | | G>A |
| NC_014760 | 400585 | MBOVPG45_RS01765 | | | T>A |
| NC_014760 | 400598 | MBOVPG45_RS01765 | | | C>T |
| NC_014760 | 400631 | MBOVPG45_RS01765 | | | T>A |
| NC_014760 | 400634 | MBOVPG45_RS01765 | | | CT>TG |
| NC_014760 | 400637 | MBOVPG45_RS01765 | | | T>C |
| NC_014760 | 400724 | MBOVPG45_RS01765 | | | G>T |
| NC_014760 | 400751 | MBOVPG45_RS01765 | | | T>A |
| NC_014760 | 400762 | MBOVPG45_RS01765 | | | AA>CG |
| NC_014760 | 400797 | MBOVPG45_RS01765 | | | G>T |
| NC_014760 | 400826 | MBOVPG45_RS01765 | | C>T | |
| NC_014760 | 400853 | MBOVPG45_RS01765 | | | C>T |
| NC_014760 | 400873 | MBOVPG45_RS01765 | | | G>C |
| NC_014760 | 400876 | MBOVPG45_RS01765 | | | TG>GA |
| NC_014760 | 400885 | MBOVPG45_RS01765 | | | - > ins GATTTCC |
| NC_014760 | 400891 | MBOVPG45_RS01765 | | | TTATTTA > del 7 |
| NC_014760 | 400903 | MBOVPG45_RS01765 | | | C>T |
| NC_014760 | 400961 | MBOVPG45_RS01765 | | | T>C |
| NC_014760 | 401042 | MBOVPG45_RS01765 | | | T>C |
| NC_014760 | 401047 | MBOVPG45_RS01765 | | | T>A |
| NC_014760 | 401050 | MBOVPG45_RS01765 | | | T>G |
| NC_014760 | 401159 | MBOVPG45_RS01765 | | | C>T |
| NC_014760 | 401183 | MBOVPG45_RS01770, MBOVPG45_RS01765 | | | C>T |
| NC_014760 | 401939 | MBOVPG45_RS01775 | | | T>C |
| NC_014760 | 402085 | MBOVPG45_RS01775 | | | C>T |

| | | | | | |
|---|---|---|---|---|---|
| NC_014760 | 402098 | MBOVPG45_RS01775 | | | C>T |
| NC_014760 | 402427 | MBOVPG45_RS01775 | | | C>T |
| NC_014760 | 402430 | MBOVPG45_RS01775 | | C>T | |
| NC_014760 | 402621 | MBOVPG45_RS01775 | | | T>G |
| NC_014760 | 403360 | MBOVPG45_RS01775 | | | C>T |
| NC_014760 | 403646 | MBOVPG45_RS01775 | | | T>C |
| NC_014760 | 404108 | MBOVPG45_RS01780 | | | G>T |
| NC_014760 | 404123 | MBOVPG45_RS01780 | | | G>C |
| NC_014760 | 404249 | MBOVPG45_RS01780 | | | C>T |
| NC_014760 | 404262 | MBOVPG45_RS01780 | | | GT>AC |
| NC_014760 | 404385 | MBOVPG45_RS01780 | | | GG>AC |
| NC_014760 | 404507 | MBOVPG45_RS01780 | | | C>T |
| NC_014760 | 404565 | MBOVPG45_RS01780 | | | T>C |
| NC_014760 | 404801 | MBOVPG45_RS01780 | | | C>T |
| NC_014760 | 404938 | MBOVPG45_RS01780 | | | C>T |
| NC_014760 | 405317 | MBOVPG45_RS01780 | | | C>T |
| NC_014760 | 405341 | MBOVPG45_RS01780 | | | T>C |
| NC_014760 | 405383 | MBOVPG45_RS01780 | | | CGC>TAT |
| NC_014760 | 405618 | MBOVPG45_RS01780 | | | T>G |
| NC_014760 | 406387 | MBOVPG45_RS01780 | | | T>C |
| NC_014760 | 406559 | MBOVPG45_RS01780 | | | T>C |
| NC_014760 | 410113 | MBOVPG45_RS01815 | | | AA>GC |
| NC_014760 | 411488 | MBOVPG45_RS01820 | | | T>C |
| NC_014760 | 411500 | MBOVPG45_RS01820 | | | T>C |
| NC_014760 | 411717 | MBOVPG45_RS01820 | | | TGCA>CATG |
| NC_014760 | 411730 | MBOVPG45_RS01820 | | | G>A |
| NC_014760 | 412080 | MBOVPG45_RS01820 | | | T>C |
| NC_014760 | 412228 | MBOVPG45_RS01820 | | | GC>CT |
| NC_014760 | 412250 | MBOVPG45_RS01820 | | | A>C |
| NC_014760 | 414758 | MBOVPG45_RS01835 | | | C>T |
| NC_014760 | 414862 | MBOVPG45_RS01835 | | | A>G |
| NC_014760 | 414883 | MBOVPG45_RS01835 | | | A>C |
| NC_014760 | 414988 | MBOVPG45_RS01835 | | | G>A |
| NC_014760 | 415004 | MBOVPG45_RS01835 | | | A>T |
| NC_014760 | 415006 | MBOVPG45_RS01835 | | | C>T |
| NC_014760 | 415076 | MBOVPG45_RS01835 | | | T>A |
| NC_014760 | 415078 | MBOVPG45_RS01835 | | | GC>AT |
| NC_014760 | 415099 | MBOVPG45_RS01835 | | | G>A |
| NC_014760 | 415197 | MBOVPG45_RS01835 | | | A>T |
| NC_014760 | 415200 | MBOVPG45_RS01835 | | | T>A |
| NC_014760 | 415241 | MBOVPG45_RS01835 | | | G>A |
| NC_014760 | 415247 | MBOVPG45_RS01835 | | | AA>GT |
| NC_014760 | 415346 | MBOVPG45_RS01835 | | | A>G |
| NC_014760 | 415349 | MBOVPG45_RS01835 | | | A>G |
| NC_014760 | 415464 | MBOVPG45_RS01835 | | | A>T |
| NC_014760 | 415468 | MBOVPG45_RS01835 | | | A>G |
| NC_014760 | 415478 | MBOVPG45_RS01835 | | | C>G |

| | | | | | |
|---|---|---|---|---|---|
| NC_014760 | 415486 | MBOVPG45_RS01835 | | | AC>CA |
| NC_014760 | 415490 | MBOVPG45_RS01835 | | | G>C |
| NC_014760 | 415506 | MBOVPG45_RS01835 | | | CC>AG |
| NC_014760 | 415509 | MBOVPG45_RS01835 | | | C>A |
| NC_014760 | 415519 | MBOVPG45_RS01835 | | | A>C |
| NC_014760 | 415532 | MBOVPG45_RS01835 | | | TG>CA |
| NC_014760 | 416161 | whiA | | | C>T |
| NC_014760 | 416257 | whiA | | | C>T |
| NC_014760 | 416834 | whiA | | | GC>AT |
| NC_014760 | 417290 | MBOVPG45_RS01845 | | | C>T |
| NC_014760 | 417510 | MBOVPG45_RS01845 | | | T>A |
| NC_014760 | 417550 | MBOVPG45_RS01845 | | | G>T |
| NC_014760 | 417836 | MBOVPG45_RS01845 | | | T>C |
| NC_014760 | 418892 | MBOVPG45_RS01850 | | | C>T |
| NC_014760 | 419355 | MBOVPG45_RS01850 | | | T>C |
| NC_014760 | 419406 | MBOVPG45_RS01850 | | | T>C |
| NC_014760 | 419734 | ligA | | | G>C |
| NC_014760 | 420001 | ligA | | | G>A |
| NC_014760 | 424904 | MBOVPG45_RS01865 | | | GC>AA |
| NC_014760 | 424907 | MBOVPG45_RS01865 | | | C>T |
| NC_014760 | 425072 | MBOVPG45_RS01865 | | | G>A |
| NC_014760 | 425081 | MBOVPG45_RS01865 | | | A>G |
| NC_014760 | 425095 | MBOVPG45_RS01865 | | | T>G |
| NC_014760 | 425104 | MBOVPG45_RS01865 | | | A>T |
| NC_014760 | 425110 | MBOVPG45_RS01865 | | | TA>AC |
| NC_014760 | 425117 | MBOVPG45_RS01865 | | | C>G |
| NC_014760 | 425126 | MBOVPG45_RS01865 | | | C>A |
| NC_014760 | 425173 | MBOVPG45_RS01865 | | | CA>TC |
| NC_014760 | 425176 | MBOVPG45_RS01865 | | | C>G |
| NC_014760 | 425183 | MBOVPG45_RS01865 | | | A>C |
| NC_014760 | 425192 | MBOVPG45_RS01865 | | | C>G |
| NC_014760 | 425281 | MBOVPG45_RS01865 | | | A>T |
| NC_014760 | 425300 | MBOVPG45_RS01865 | | | G>C |
| NC_014760 | 425309 | MBOVPG45_RS01865 | | | G>A |
| NC_014760 | 425438 | MBOVPG45_RS01865 | | | A>G |
| NC_014760 | 425441 | MBOVPG45_RS01865 | | | GGT>AAA |
| NC_014760 | 425500 | MBOVPG45_RS01865 | | | CA>TG |
| NC_014760 | 425543 | MBOVPG45_RS01865 | | | G>A |
| NC_014760 | 425797 | MBOVPG45_RS01865 | | | A>G |
| NC_014760 | 425809 | MBOVPG45_RS01865 | | | A>T |
| NC_014760 | 425814 | MBOVPG45_RS01865 | | | A>T |
| NC_014760 | 425829 | MBOVPG45_RS01865 | | | T>C |
| NC_014760 | 425835 | MBOVPG45_RS01865 | | | A>T |
| NC_014760 | 425859 | MBOVPG45_RS01865 | | | T>C |
| NC_014760 | 425914 | MBOVPG45_RS01865 | | | A>T |
| NC_014760 | 426032 | MBOVPG45_RS01865 | | | G>T |
| NC_014760 | 426146 | MBOVPG45_RS01865 | | | A>G |

| | | | | | |
|---|---|---|---|---|---|
| NC_014760 | 426284 | MBOVPG45_RS01865 | | | G>A |
| NC_014760 | 426289 | MBOVPG45_RS01865 | | | A>C |
| NC_014760 | 426542 | MBOVPG45_RS01865 | | | T>C |
| NC_014760 | 426653 | MBOVPG45_RS01865 | | | TC>CA |
| NC_014760 | 426693 | MBOVPG45_RS01865 | | | C>A |
| NC_014760 | 426836 | MBOVPG45_RS01870 | | | G>A |
| NC_014760 | 427161 | MBOVPG45_RS01870 | | | G>A |
| NC_014760 | 427188 | MBOVPG45_RS01870 | | | C>T |
| NC_014760 | 427222 | MBOVPG45_RS01870 | | | C>G |
| NC_014760 | 427307 | MBOVPG45_RS01870 | | | G>C |
| NC_014760 | 427341 | MBOVPG45_RS01870 | | | C>T |
| NC_014760 | 428749 | MBOVPG45_RS01870 | | G>A | |
| NC_014760 | 430161 | MBOVPG45_RS01875 | | | G>A |
| NC_014760 | 433117 | MBOVPG45_RS01880 | | | G>A |
| NC_014760 | 434748 | MBOVPG45_RS01885 | | | A>T |
| NC_014760 | 437928 | MBOVPG45_RS01905 | | | G>A |
| NC_014760 | 438288 | MBOVPG45_RS01905 | | | A>C |
| NC_014760 | 439044 | MBOVPG45_RS01905 | | | A>G |
| NC_014760 | 443328 | MBOVPG45_RS01920 | | | G>T |
| NC_014760 | 443538 | MBOVPG45_RS01920 | | | G>T |
| NC_014760 | 444935 | MBOVPG45_RS01925 | | | C>T |
| NC_014760 | 447801 | MBOVPG45_RS01935 | | C>T | |
| NC_014760 | 449716 | MBOVPG45_RS01940 | | A>G | |
| NC_014760 | 450306 | MBOVPG45_RS01940 | | | T>C |
| NC_014760 | 450464 | MBOVPG45_RS01940 | | | T>G |
| NC_014760 | 450657 | MBOVPG45_RS01940 | | | C>T |
| NC_014760 | 451398 | MBOVPG45_RS01945 | | | T>G |
| NC_014760 | 451663 | MBOVPG45_RS01945 | | | A>G |
| NC_014760 | 454173 | asnS | | | T>C |
| NC_014760 | 455097 | MBOVPG45_RS01960 | | | A>T |
| NC_014760 | 456260 | MBOVPG45_RS01970 | | | T>A |
| NC_014760 | 462062 | MBOVPG45_RS02005 | | | C>T |
| NC_014760 | 463061 | MBOVPG45_RS02005 | | | T>C |
| NC_014760 | 463304 | MBOVPG45_RS02005 | | | C>G |
| NC_014760 | 463373 | MBOVPG45_RS02005 | | | C>T |
| NC_014760 | 463676 | MBOVPG45_RS02005 | | | T>C |
| NC_014760 | 463976 | MBOVPG45_RS02010 | | | T>A |
| NC_014760 | 463997 | MBOVPG45_RS02010 | | | G>T |
| NC_014760 | 464003 | MBOVPG45_RS02010 | | | AA>TG |
| NC_014760 | 464069 | MBOVPG45_RS02010 | | | G>T |
| NC_014760 | 464129 | MBOVPG45_RS02010 | | C>A | |
| NC_014760 | 464204 | MBOVPG45_RS02010 | | | T>A |
| NC_014760 | 464350 | MBOVPG45_RS02010 | | | G>T |
| NC_014760 | 464429 | MBOVPG45_RS02010 | | | C>T |
| NC_014760 | 464497 | MBOVPG45_RS02010 | | | TT>CC |
| NC_014760 | 464560 | MBOVPG45_RS02010 | | | C>T |
| NC_014760 | 464642 | MBOVPG45_RS02010 | | | C>T |

| | | | | | |
|---|---|---|---|---|---|
| NC_014760 | 464645 | MBOVPG45_RS02010 | | | T>C |
| NC_014760 | 464654 | MBOVPG45_RS02010 | | | T>C |
| NC_014760 | 464663 | MBOVPG45_RS02010 | | | C>T |
| NC_014760 | 464695 | MBOVPG45_RS02010 | | | C>T |
| NC_014760 | 464801 | MBOVPG45_RS02010 | | | C>T |
| NC_014760 | 464947 | MBOVPG45_RS02010 | | | T>C |
| NC_014760 | 464953 | MBOVPG45_RS02010 | | | AGA>TCC |
| NC_014760 | 464958 | MBOVPG45_RS02010 | | | CAT>AGC |
| NC_014760 | 464963 | MBOVPG45_RS02010 | | | G>A |
| NC_014760 | 464970 | MBOVPG45_RS02010 | | | AACA > del 4 |
| NC_014760 | 464974 | MBOVPG45_RS02010 | | | T>G |
| NC_014760 | 464977 | MBOVPG45_RS02010 | | | - > ins CTCA |
| NC_014760 | 465022 | MBOVPG45_RS02010 | | | A>G |
| NC_014760 | 465056 | MBOVPG45_RS02010 | | | C>T |
| NC_014760 | 465107 | MBOVPG45_RS02010 | | | C>T |
| NC_014760 | 465118 | MBOVPG45_RS02010 | | | TT>CC |
| NC_014760 | 465286 | MBOVPG45_RS02010 | | | A>G |
| NC_014760 | 465289 | MBOVPG45_RS02010 | | | C>T |
| NC_014760 | 465296 | MBOVPG45_RS02010 | | | C>T |
| NC_014760 | 465374 | MBOVPG45_RS02010 | | | G>T |
| NC_014760 | 465497 | MBOVPG45_RS02010 | | | T>C |
| NC_014760 | 465500 | MBOVPG45_RS02010 | | | G>A |
| NC_014760 | 465526 | MBOVPG45_RS02010 | | | A>C |
| NC_014760 | 465565 | MBOVPG45_RS02010 | | | A>G |
| NC_014760 | 465586 | MBOVPG45_RS02010 | | | T>C |
| NC_014760 | 465684 | MBOVPG45_RS02010 | | | T>A |
| NC_014760 | 465839 | MBOVPG45_RS02010 | | | T>C |
| NC_014760 | 466370 | msrB | | | T>C |
| NC_014760 | 466619 | msrB | | | C>T |
| NC_014760 | 466982 | MBOVPG45_RS02020 | | | C>A |
| NC_014760 | 467001 | MBOVPG45_RS02020 | | | G>T |
| NC_014760 | 467128 | MBOVPG45_RS02020 | | C > del 1 | |
| NC_014760 | 467173 | MBOVPG45_RS02020 | | | C>T |
| NC_014760 | 467471 | mutM | | | C>G |
| NC_014760 | 467519 | mutM | | | T>C |
| NC_014760 | 467630 | mutM | | | G>A |
| NC_014760 | 467996 | mutM | | | TT>CC |
| NC_014760 | 468171 | MBOVPG45_RS02030 | | | C>T |
| NC_014760 | 469103 | MBOVPG45_RS02030 | | | A>C |
| NC_014760 | 469244 | MBOVPG45_RS02030 | | | T>C |
| NC_014760 | 469302 | MBOVPG45_RS02030 | | | G>A |
| NC_014760 | 469380 | MBOVPG45_RS02035 | | | C>T |
| NC_014760 | 469412 | MBOVPG45_RS02035 | | | AA>GC |
| NC_014760 | 469531 | MBOVPG45_RS02035 | | | T>C |
| NC_014760 | 469537 | MBOVPG45_RS02035 | | | C>T |
| NC_014760 | 469660 | MBOVPG45_RS02035 | | | T>G |
| NC_014760 | 469729 | MBOVPG45_RS02035 | | | T>C |

| | | | | | |
|---|---|---|---|---|---|
| NC_014760 | 469811 | MBOVPG45_RS02035 | | | A>C |
| NC_014760 | 470276 | MBOVPG45_RS02035 | | | T>A |
| NC_014760 | 470412 | MBOVPG45_RS02035 | | | G>A |
| NC_014760 | 470536 | MBOVPG45_RS02035 | | | G>C |
| NC_014760 | 470542 | MBOVPG45_RS02035 | | | T>A |
| NC_014760 | 471148 | MBOVPG45_RS02045 | | | C>A |
| NC_014760 | 471469 | MBOVPG45_RS02045 | | | T>C |
| NC_014760 | 472081 | MBOVPG45_RS02045 | | | T>C |
| NC_014760 | 472165 | MBOVPG45_RS02045 | | | C>T |
| NC_014760 | 473268 | MBOVPG45_RS02050 | | | G>A |
| NC_014760 | 473494 | MBOVPG45_RS02050 | | | T>A |
| NC_014760 | 474125 | MBOVPG45_RS02050 | | | T>A |
| NC_014760 | 474207 | MBOVPG45_RS02050 | | | A>G |
| NC_014760 | 474250 | MBOVPG45_RS02050 | | | A>G |
| NC_014760 | 476136 | MBOVPG45_RS02060 | | | G>A |
| NC_014760 | 477575 | tsaE | | | G>A |
| NC_014760 | 477606 | tsaE | | | A>T |
| NC_014760 | 477995 | MBOVPG45_RS02070 | | | G>A |
| NC_014760 | 478198 | MBOVPG45_RS02070 | | | C>A |
| NC_014760 | 478207 | MBOVPG45_RS02070 | | | A>C |
| NC_014760 | 478218 | MBOVPG45_RS02070 | | | GC>TA |
| NC_014760 | 478888 | tsaD | | | A>G |
| NC_014760 | 479040 | tsaD | | | AG>GC |
| NC_014760 | 479074 | tsaD | | | G>T |
| NC_014760 | 479290 | MBOVPG45_RS02080 | | | C>T |
| NC_014760 | 479431 | MBOVPG45_RS02080 | | | A>G |
| NC_014760 | 479441 | MBOVPG45_RS02080 | | | G>A |
| NC_014760 | 479443 | MBOVPG45_RS02080 | | | A>G |
| NC_014760 | 479527 | MBOVPG45_RS02080 | | | G>A |
| NC_014760 | 479581 | MBOVPG45_RS02080 | | | A>C |
| NC_014760 | 479698 | MBOVPG45_RS02080 | | | G>A |
| NC_014760 | 479770 | MBOVPG45_RS02080 | | | C>A |
| NC_014760 | 479778 | MBOVPG45_RS02080 | | | T>G |
| NC_014760 | 480149 | MBOVPG45_RS02080 | | | C>A |
| NC_014760 | 480232 | MBOVPG45_RS02080 | | | A>T |
| NC_014760 | 480271 | MBOVPG45_RS02080 | | | A>C |
| NC_014760 | 480280 | MBOVPG45_RS02080 | | | AT>GC |
| NC_014760 | 480355 | MBOVPG45_RS02080 | | | A>G |
| NC_014760 | 480580 | MBOVPG45_RS02080 | | | G>A |
| NC_014760 | 480587 | MBOVPG45_RS02080 | | | GA>AC |
| NC_014760 | 480613 | MBOVPG45_RS02080 | | | G>A |
| NC_014760 | 480621 | MBOVPG45_RS02080 | | | A>T |
| NC_014760 | 480864 | MBOVPG45_RS02080 | | | A>T |
| NC_014760 | 480967 | MBOVPG45_RS02080 | | | G>A |
| NC_014760 | 480971 | MBOVPG45_RS02080 | | | A>C |
| NC_014760 | 487031 | MBOVPG45_RS02105 | | | A>T |
| NC_014760 | 488484 | MBOVPG45_RS02110 | | | A>G |

| | | | | | |
|---|---|---|---|---|---|
| NC_014760 | 491269 | MBOVPG45_RS02120 | | | T>G |
| NC_014760 | 491437 | MBOVPG45_RS02120 | | | T>C |
| NC_014760 | 491637 | MBOVPG45_RS02120 | | | T>C |
| NC_014760 | 491801 | MBOVPG45_RS02120 | | | T>G |
| NC_014760 | 491930 | MBOVPG45_RS02120 | | | A>T |
| NC_014760 | 491958 | MBOVPG45_RS02120 | | | C>T |
| NC_014760 | 492094 | MBOVPG45_RS02120 | | | T>G |
| NC_014760 | 492174 | MBOVPG45_RS02120 | | | C>T |
| NC_014760 | 492196 | MBOVPG45_RS02120 | | | T>C |
| NC_014760 | 492244 | MBOVPG45_RS02120 | | | G>C |
| NC_014760 | 492271 | MBOVPG45_RS02120 | | | T>C |
| NC_014760 | 492422 | MBOVPG45_RS02120 | | | T>C |
| NC_014760 | 492526 | MBOVPG45_RS02120 | | | C>T |
| NC_014760 | 492609 | MBOVPG45_RS02120 | | | T>C |
| NC_014760 | 492624 | MBOVPG45_RS02120 | | | A>C |
| NC_014760 | 492658 | MBOVPG45_RS02120 | | | T>C |
| NC_014760 | 492723 | MBOVPG45_RS02120 | | | T>C |
| NC_014760 | 493605 | MBOVPG45_RS02125 | | - > ins T | |
| NC_014760 | 494007 | MBOVPG45_RS02125 | | | G>T |
| NC_014760 | 494230 | MBOVPG45_RS02125 | | | T>A |
| NC_014760 | 494385 | MBOVPG45_RS02125 | | | T>C |
| NC_014760 | 494529 | MBOVPG45_RS02125 | | | A>G |
| NC_014760 | 494778 | MBOVPG45_RS02125 | | | C>T |
| NC_014760 | 494903 | MBOVPG45_RS02125 | | | A>G |
| NC_014760 | 495020 | MBOVPG45_RS02125 | | | A>G |
| NC_014760 | 495033 | MBOVPG45_RS02125 | | | C>T |
| NC_014760 | 495053 | MBOVPG45_RS02125 | | | C>A |
| NC_014760 | 495056 | MBOVPG45_RS02125 | | | T>C |
| NC_014760 | 496543 | MBOVPG45_RS02130 | | | T>C |
| NC_014760 | 496546 | MBOVPG45_RS02130 | | | C>T |
| NC_014760 | 496872 | MBOVPG45_RS02130 | | | T > del 1 |
| NC_014760 | 496897 | MBOVPG45_RS02130 | | | C>T |
| NC_014760 | 497123 | MBOVPG45_RS02130 | | | T>C |
| NC_014760 | 497281 | MBOVPG45_RS02130 | | | T>G |
| NC_014760 | 497287 | MBOVPG45_RS02130 | | | A>G |
| NC_014760 | 497318 | MBOVPG45_RS02130 | | | C>A |
| NC_014760 | 497823 | MBOVPG45_RS02135 | | | A>G |
| NC_014760 | 497826 | MBOVPG45_RS02135 | | | G>A |
| NC_014760 | 497839 | MBOVPG45_RS02135 | | | T>C |
| NC_014760 | 497946 | MBOVPG45_RS02135 | | | A>G |
| NC_014760 | 498112 | MBOVPG45_RS02135 | | | G>A |
| NC_014760 | 498724 | MBOVPG45_RS02140 | | | G>A |
| NC_014760 | 500449 | MBOVPG45_RS02150 | | | GGG > del 3 |
| NC_014760 | 500489 | MBOVPG45_RS02150 | | | C>T |
| NC_014760 | 500534 | MBOVPG45_RS02150 | | | G>A |
| NC_014760 | 500539 | MBOVPG45_RS02150 | | | A>G |
| NC_014760 | 500549 | MBOVPG45_RS02150 | | | G>A |

| | | | | | |
|---|---|---|---|---|---|
| NC_014760 | 500605 | MBOVPG45_RS02150 | | | C>G |
| NC_014760 | 500613 | MBOVPG45_RS02150 | | | A>C |
| NC_014760 | 500623 | MBOVPG45_RS02150 | | | G>C |
| NC_014760 | 500652 | MBOVPG45_RS02150 | | | A>C |
| NC_014760 | 500657 | MBOVPG45_RS02150 | | | A>G |
| NC_014760 | 500710 | MBOVPG45_RS02150 | | | G>A |
| NC_014760 | 500735 | MBOVPG45_RS02150 | | | T>C |
| NC_014760 | 500741 | MBOVPG45_RS02150 | | | A>G |
| NC_014760 | 500754 | MBOVPG45_RS02150 | | | G>C |
| NC_014760 | 500763 | MBOVPG45_RS02150 | | | T>A |
| NC_014760 | 500786 | MBOVPG45_RS02150 | | G>T | G>A |
| NC_014760 | 500813 | MBOVPG45_RS02150 | | | T>A |
| NC_014760 | 501101 | MBOVPG45_RS02150 | | | GAC > del 3 |
| NC_014760 | 501954 | MBOVPG45_RS02155 | | | A>G |
| NC_014760 | 502829 | MBOVPG45_RS02155 | | | A>G |
| NC_014760 | 502830 | MBOVPG45_RS02155 | | | A>G |
| NC_014760 | 503162 | MBOVPG45_RS02160 | | | TA>GT |
| NC_014760 | 503660 | MBOVPG45_RS02160 | | | C>T |
| NC_014760 | 504209 | trpS | | | T>A |
| NC_014760 | 505264 | MBOVPG45_RS02170 | | | C>A |
| NC_014760 | 506113 | MBOVPG45_RS02170 | | | T>C |
| NC_014760 | 506166 | MBOVPG45_RS02170 | | | GT>AC |
| NC_014760 | 506367 | MBOVPG45_RS02170 | | | A>T |
| NC_014760 | 506466 | MBOVPG45_RS02170 | | C>T | |
| NC_014760 | 506476 | MBOVPG45_RS02170 | | | C>T |
| NC_014760 | 506547 | MBOVPG45_RS02170 | | | A>G |
| NC_014760 | 506551 | MBOVPG45_RS02170 | | | A>G |
| NC_014760 | 506659 | MBOVPG45_RS02170 | | | A>G |
| NC_014760 | 506905 | MBOVPG45_RS04420 | | | T>A |
| NC_014760 | 506916 | MBOVPG45_RS04420 | | | G>T |
| NC_014760 | 507065 | MBOVPG45_RS04420 | | | A>C |
| NC_014760 | 507142 | MBOVPG45_RS04420 | | | G>A |
| NC_014760 | 507923 | MBOVPG45_RS02180 | | | A>T |
| NC_014760 | 507930 | MBOVPG45_RS02180 | | | T>C |
| NC_014760 | 507933 | MBOVPG45_RS02180 | | | C>G |
| NC_014760 | 508164 | MBOVPG45_RS02180 | | | T>C |
| NC_014760 | 508231 | MBOVPG45_RS02180 | | | T>A |
| NC_014760 | 508247 | MBOVPG45_RS02180 | | | T>G |
| NC_014760 | 508281 | MBOVPG45_RS02180 | | | T>C |
| NC_014760 | 508294 | MBOVPG45_RS02180 | | | AT>GC |
| NC_014760 | 508304 | MBOVPG45_RS02180 | | | TT>CC |
| NC_014760 | 508329 | MBOVPG45_RS02180 | | | A>C |
| NC_014760 | 508356 | MBOVPG45_RS02180 | | | G>C |
| NC_014760 | 508422 | MBOVPG45_RS02180 | | | T>C |
| NC_014760 | 508424 | MBOVPG45_RS02180 | | | C>T |
| NC_014760 | 508440 | MBOVPG45_RS02180 | | | T>C |
| NC_014760 | 508825 | MBOVPG45_RS02185 | | | T>C |

| | | | | | |
|---|---|---|---|---|---|
| NC_014760 | 508827 | MBOVPG45_RS02185 | | | G>A |
| NC_014760 | 508854 | MBOVPG45_RS02185 | | | A>G |
| NC_014760 | 508863 | MBOVPG45_RS02185 | | | T>C |
| NC_014760 | 508932 | MBOVPG45_RS02185 | | | G>A |
| NC_014760 | 508953 | MBOVPG45_RS02185 | | | T>G |
| NC_014760 | 508970 | MBOVPG45_RS02185 | | | T>C |
| NC_014760 | 509112 | MBOVPG45_RS02185 | | | G>A |
| NC_014760 | 509161 | MBOVPG45_RS02185 | | | C>T |
| NC_014760 | 509214 | MBOVPG45_RS02185 | | | A>G |
| NC_014760 | 509350 | MBOVPG45_RS02185 | | | A>G |
| NC_014760 | 509357 | MBOVPG45_RS02185 | | | A>C |
| NC_014760 | 509770 | MBOVPG45_RS02185 | | | T>C |
| NC_014760 | 509794 | MBOVPG45_RS02185 | | | T>A |
| NC_014760 | 509813 | MBOVPG45_RS02185 | | | A>C |
| NC_014760 | 509852 | MBOVPG45_RS02185 | | | T>C |
| NC_014760 | 509917 | MBOVPG45_RS02185 | | | C>T |
| NC_014760 | 509946 | MBOVPG45_RS02185 | | | G>T |
| NC_014760 | 510067 | MBOVPG45_RS02185 | | | TA>CG |
| NC_014760 | 510171 | MBOVPG45_RS02185 | | | A>T |
| NC_014760 | 510240 | MBOVPG45_RS02185 | | | G>A |
| NC_014760 | 510784 | MBOVPG45_RS02190 | | | G>A |
| NC_014760 | 510933 | MBOVPG45_RS02190 | | | C>T |
| NC_014760 | 511038 | MBOVPG45_RS02190 | | | G>C |
| NC_014760 | 511107 | MBOVPG45_RS02190 | | | A > del 1 |
| NC_014760 | 512844 | atpH | | | T>A |
| NC_014760 | 512945 | atpH | | | G>A |
| NC_014760 | 512953 | atpH | | | G>A |
| NC_014760 | 512956 | atpH | | | A>T |
| NC_014760 | 513447 | MBOVPG45_RS02215 | | | A>G |
| NC_014760 | 515730 | MBOVPG45_RS02220 | | A>G | |
| NC_014760 | 517631 | MBOVPG45_RS02230 | | | G>T |
| NC_014760 | 519897 | MBOVPG45_RS02245 | | | A>G |
| NC_014760 | 521292 | lon | | | A>C |
| NC_014760 | 521294 | lon | | C>T | |
| NC_014760 | 522302 | lon | | | A>C |
| NC_014760 | 522518 | lon | | | A>G |
| NC_014760 | 522537 | lon | | | A>T |
| NC_014760 | 523679 | MBOVPG45_RS02255 | | | T>A |
| NC_014760 | 524097 | MBOVPG45_RS02255 | | | C>T |
| NC_014760 | 524121 | MBOVPG45_RS02255 | | | C>T |
| NC_014760 | 524509 | MBOVPG45_RS02255 | | | C>T |
| NC_014760 | 524586 | MBOVPG45_RS02255 | | | A>G |
| NC_014760 | 524612 | MBOVPG45_RS02255 | | | T>A |
| NC_014760 | 524777 | MBOVPG45_RS02255 | | | T>C |
| NC_014760 | 524798 | MBOVPG45_RS02255 | | | A>T |
| NC_014760 | 525121 | MBOVPG45_RS02255 | | | T>C |
| NC_014760 | 525136 | MBOVPG45_RS02255 | | | T>C |

| | | | | | |
|---|---|---|---|---|---|
| NC_014760 | 525219 | MBOVPG45_RS02255 | | | C>T |
| NC_014760 | 525250 | MBOVPG45_RS02255 | | | T>C |
| NC_014760 | 525372 | MBOVPG45_RS02255 | | | T>A |
| NC_014760 | 525867 | MBOVPG45_RS02255 | | | C>T |
| NC_014760 | 525904 | MBOVPG45_RS02255 | | | T>A |
| NC_014760 | 525907 | MBOVPG45_RS02255 | | | A>G |
| NC_014760 | 526035 | MBOVPG45_RS02260 | | | C>T |
| NC_014760 | 526112 | MBOVPG45_RS02260 | | | T>A |
| NC_014760 | 526134 | MBOVPG45_RS02260 | | | T>C |
| NC_014760 | 526152 | MBOVPG45_RS02260 | | | TG>CA |
| NC_014760 | 526268 | MBOVPG45_RS02260 | | | G>A |
| NC_014760 | 526274 | MBOVPG45_RS02260 | | | C>T |
| NC_014760 | 526509 | MBOVPG45_RS02260 | | | C>T |
| NC_014760 | 526560 | MBOVPG45_RS02260 | | | C>T |
| NC_014760 | 526621 | MBOVPG45_RS02260 | | | T>A |
| NC_014760 | 527473 | ssb | | | A>G |
| NC_014760 | 527506 | ssb | | | C>T |
| NC_014760 | 527527 | ssb | | | C>G |
| NC_014760 | 527533 | ssb | | | A>G |
| NC_014760 | 527720 | ssb | | C>T | |
| NC_014760 | 528299 | MBOVPG45_RS02275 | | | C>T |
| NC_014760 | 528754 | MBOVPG45_RS02275 | | | A>G |
| NC_014760 | 528974 | MBOVPG45_RS02275 | | | T>C |
| NC_014760 | 529217 | MBOVPG45_RS02275 | | | G>A |
| NC_014760 | 529460 | MBOVPG45_RS02275 | | | T>A |
| NC_014760 | 529465 | MBOVPG45_RS02275 | | | C>T |
| NC_014760 | 529601 | MBOVPG45_RS02275 | | | A>C |
| NC_014760 | 529642 | MBOVPG45_RS02275 | | | T>A |
| NC_014760 | 529772 | MBOVPG45_RS02275 | | | T>C |
| NC_014760 | 529969 | MBOVPG45_RS02280 | (uracil(1498)-N(3))-methyltransferas | | C>T |
| NC_014760 | 530197 | MBOVPG45_RS02280 | (uracil(1498)-N(3))-methyltransferas | | G>A |
| NC_014760 | 530519 | MBOVPG45_RS02280 | (uracil(1498)-N(3))-methyltransferas | | A>C |
| NC_014760 | 530549 | MBOVPG45_RS02280 | (uracil(1498)-N(3))-methyltransferas | | T>C |
| NC_014760 | 530918 | MBOVPG45_RS02285 | | | G>A |
| NC_014760 | 530923 | MBOVPG45_RS02285 | | | A>T |
| NC_014760 | 531011 | MBOVPG45_RS02285 | | | C>T |
| NC_014760 | 531106 | MBOVPG45_RS02285 | | | G>A |
| NC_014760 | 531127 | MBOVPG45_RS02285 | | | A>T |
| NC_014760 | 531421 | MBOVPG45_RS02290 | | | T>A |

| | | | | | |
|---|---|---|---|---|---|
| NC_014760 | 531438 | MBOVPG45_RS02290 | | | G>A |
| NC_014760 | 532709 | MBOVPG45_RS02295 | | | A>C |
| NC_014760 | 532945 | MBOVPG45_RS02295 | | | T>C |
| NC_014760 | 533326 | MBOVPG45_RS02295 | | | G>A |
| NC_014760 | 533495 | MBOVPG45_RS02295 | | | C>A |
| NC_014760 | 533589 | MBOVPG45_RS02295 | | A>C | A>C |
| NC_014760 | 534120 | rsmH | | | A>C |
| NC_014760 | 534155 | rsmH | | | G>A |
| NC_014760 | 534221 | rsmH | | | T>C |
| NC_014760 | 534449 | rsmH | | | A>G |
| NC_014760 | 534654 | rsmH | | | T>C |
| NC_014760 | 535888 | MBOVPG45_RS02310 | | | A>C |
| NC_014760 | 536751 | MBOVPG45_RS02315 | | | G>A |
| NC_014760 | 537640 | MBOVPG45_RS02315 | | | A>G |
| NC_014760 | 538024 | MBOVPG45_RS02320 | | | A>G |
| NC_014760 | 538752 | dcm | | | T>C |
| NC_014760 | 539019 | dcm | | | C>T |
| NC_014760 | 539235 | dcm | | | T>C |
| NC_014760 | 539370 | dcm | | | C>A |
| NC_014760 | 540693 | uvrB | | | A>G |
| NC_014760 | 540926 | uvrB | | | AA>GG |
| NC_014760 | 541115 | uvrB | | | G>A |
| NC_014760 | 541358 | uvrB | | | A>T |
| NC_014760 | 541422 | uvrB | | | G>T |
| NC_014760 | 541772 | uvrB | | | T>A |
| NC_014760 | 541893 | uvrB | | | A>G |
| NC_014760 | 542036 | uvrB | | | G>A |
| NC_014760 | 542088 | uvrB | | | C>T |
| NC_014760 | 542155 | uvrA | | | G>A |
| NC_014760 | 542169 | uvrA | | | CC>TA |
| NC_014760 | 542566 | uvrA | | | A>G |
| NC_014760 | 542813 | uvrA | | | C>T |
| NC_014760 | 542824 | uvrA | | | A>G |
| NC_014760 | 543869 | uvrA | | | A>G |
| NC_014760 | 543905 | uvrA | | | C>T |
| NC_014760 | 543989 | uvrA | | | C>T |
| NC_014760 | 544264 | uvrA | | | G>A |
| NC_014760 | 544453 | uvrA | | | G>T |
| NC_014760 | 545064 | MBOVPG45_RS02345 | | | A>C |
| NC_014760 | 545201 | MBOVPG45_RS02345 | | | G>T |
| NC_014760 | 545347 | MBOVPG45_RS02345 | | | A>G |
| NC_014760 | 545389 | MBOVPG45_RS02345 | | | A>G |
| NC_014760 | 545563 | MBOVPG45_RS02345 | | | G>A |
| NC_014760 | 545610 | MBOVPG45_RS02350, MBOVPG45_RS02345 | | | G>A |
| NC_014760 | 545712 | MBOVPG45_RS02350 | | | A>G |
| NC_014760 | 545757 | MBOVPG45_RS02350 | | | C>T |

| | | | | | |
|---|---|---|---|---|---|
| NC_014760 | 545823 | MBOVPG45_RS02350 | | | G>T |
| NC_014760 | 545944 | MBOVPG45_RS02350 | | | G>T |
| NC_014760 | 545975 | MBOVPG45_RS02350 | | | G>A |
| NC_014760 | 551171 | MBOVPG45_RS02380 | | | T>A |
| NC_014760 | 552200 | MBOVPG45_RS02385 | | | C>T |
| NC_014760 | 553973 | MBOVPG45_RS02390 | | | A>T |
| NC_014760 | 553975 | MBOVPG45_RS02390 | | | T>C |
| NC_014760 | 553981 | MBOVPG45_RS02390 | | | C>T |
| NC_014760 | 553984 | MBOVPG45_RS02390 | | | CT>TG |
| NC_014760 | 553992 | MBOVPG45_RS02390 | | | C>T |
| NC_014760 | 555460 | MBOVPG45_RS02390 | | | T>G |
| NC_014760 | 555720 | MBOVPG45_RS02390 | | | TG>GT |
| NC_014760 | 555738 | MBOVPG45_RS02390 | | | C>T |
| NC_014760 | 555798 | MBOVPG45_RS02390 | | | A>G |
| NC_014760 | 555820 | MBOVPG45_RS02390 | | | T>C |
| NC_014760 | 555826 | MBOVPG45_RS02390 | | | C>T |
| NC_014760 | 555866 | MBOVPG45_RS02390 | | | A>T |
| NC_014760 | 555880 | MBOVPG45_RS02390 | | | C>T |
| NC_014760 | 556007 | MBOVPG45_RS02390 | | | A>T |
| NC_014760 | 556010 | MBOVPG45_RS02390 | | | TGG>GTT |
| NC_014760 | 556024 | MBOVPG45_RS02390 | | | T>C |
| NC_014760 | 556028 | MBOVPG45_RS02390 | | | CGGC>TTCT |
| NC_014760 | 556047 | MBOVPG45_RS02390 | | | T>C |
| NC_014760 | 556135 | MBOVPG45_RS02390 | | | C>T |
| NC_014760 | 556137 | MBOVPG45_RS02390 | | | T>C |
| NC_014760 | 556153 | MBOVPG45_RS02390 | | | C>T |
| NC_014760 | 556161 | MBOVPG45_RS02390 | | | GT>AC |
| NC_014760 | 556466 | MBOVPG45_RS02390 | | | AC>TG |
| NC_014760 | 556473 | MBOVPG45_RS02390 | | | GG>AT |
| NC_014760 | 556478 | MBOVPG45_RS02390 | | | AG>TT |
| NC_014760 | 557052 | MBOVPG45_RS02390 | | | A>G |
| NC_014760 | 557067 | MBOVPG45_RS02390 | | | G>A |
| NC_014760 | 557345 | MBOVPG45_RS02395 | | | G>C |
| NC_014760 | 557376 | MBOVPG45_RS02395 | | | A>T |
| NC_014760 | 557481 | MBOVPG45_RS02395 | | | A>T |
| NC_014760 | 557763 | MBOVPG45_RS02395 | | | A>T |
| NC_014760 | 557979 | MBOVPG45_RS02400 | | C>T | |
| NC_014760 | 559822 | MBOVPG45_RS02400 | | T>C | T>C |
| NC_014760 | 559972 | MBOVPG45_RS02400 | | | A>G |
| NC_014760 | 561474 | MBOVPG45_RS02405 | | | C>T |
| NC_014760 | 561488 | MBOVPG45_RS02405 | | | A>G |
| NC_014760 | 561570 | MBOVPG45_RS02405 | | | A>T |
| NC_014760 | 561944 | MBOVPG45_RS02405 | | | G>A |
| NC_014760 | 562005 | MBOVPG45_RS02405 | | | TT>CA |
| NC_014760 | 563125 | MBOVPG45_RS02410 | | | C>T |
| NC_014760 | 564824 | MBOVPG45_RS02420 | | | A>G |
| NC_014760 | 565237 | MBOVPG45_RS02420 | | | - > ins T |

| | | | | | |
|---|---|---|---|---|---|
| NC_014760 | 565492 | MBOVPG45_RS02425 | | | T>A |
| NC_014760 | 565510 | MBOVPG45_RS02425 | | | TTCTAGTTC > del 9 |
| NC_014760 | 565524 | MBOVPG45_RS02425 | | | C>T |
| NC_014760 | 565527 | MBOVPG45_RS02425 | | | T>C |
| NC_014760 | 565536 | MBOVPG45_RS02425 | | | T>C |
| NC_014760 | 565538 | MBOVPG45_RS02425 | | | A>T |
| NC_014760 | 565552 | MBOVPG45_RS02425 | | | T>G |
| NC_014760 | 565564 | MBOVPG45_RS02425 | | | T>A |
| NC_014760 | 566186 | MBOVPG45_RS02430 | | | A>T |
| NC_014760 | 566238 | MBOVPG45_RS02430 | | | A>C |
| NC_014760 | 566243 | MBOVPG45_RS02430 | | | G>A |
| NC_014760 | 566271 | MBOVPG45_RS02430 | | | G>T |
| NC_014760 | 566794 | MBOVPG45_RS02430 | | | A>T |
| NC_014760 | 566912 | MBOVPG45_RS02435 | | | A>G |
| NC_014760 | 567177 | MBOVPG45_RS02435 | | | A>C |
| NC_014760 | 567250 | MBOVPG45_RS02440 | | | C>T |
| NC_014760 | 567268 | MBOVPG45_RS02440 | | | C>A |
| NC_014760 | 567301 | MBOVPG45_RS02440 | | | T>C |
| NC_014760 | 567327 | MBOVPG45_RS02440 | | | T>C |
| NC_014760 | 567427 | MBOVPG45_RS02440 | | | A>T |
| NC_014760 | 567440 | MBOVPG45_RS02440 | | | AG>GA |
| NC_014760 | 567514 | MBOVPG45_RS02445 | | | TC>CT |
| NC_014760 | 568134 | MBOVPG45_RS02445 | | | T>A |
| NC_014760 | 568142 | MBOVPG45_RS02445 | | | C>G |
| NC_014760 | 569339 | MBOVPG45_RS02455 | | | T>C |
| NC_014760 | 569367 | MBOVPG45_RS02455 | | | T>A |
| NC_014760 | 569845 | MBOVPG45_RS02455 | | | C>G |
| NC_014760 | 569979 | MBOVPG45_RS02455 | | | C>T |
| NC_014760 | 570373 | MBOVPG45_RS02455 | | | T>C |
| NC_014760 | 570980 | MBOVPG45_RS02460 | | | T>A |
| NC_014760 | 571002 | MBOVPG45_RS02460 | | | T>C |
| NC_014760 | 571014 | MBOVPG45_RS02460 | | | T>C |
| NC_014760 | 571077 | MBOVPG45_RS02460 | | | T>G |
| NC_014760 | 571239 | MBOVPG45_RS02460 | | | T>C |
| NC_014760 | 571267 | MBOVPG45_RS02460 | | | A>T |
| NC_014760 | 571686 | MBOVPG45_RS02460 | | | C>T |
| NC_014760 | 571695 | MBOVPG45_RS02460 | | | T>C |
| NC_014760 | 571787 | MBOVPG45_RS02460 | | | GG>AA |
| NC_014760 | 571791 | MBOVPG45_RS02460 | | | T>C |
| NC_014760 | 572379 | MBOVPG45_RS04635 | | | G>A |
| NC_014760 | 572397 | MBOVPG45_RS04635 | | | T>C |
| NC_014760 | 572449 | MBOVPG45_RS04635 | | | C>T |
| NC_014760 | 572471 | MBOVPG45_RS04635 | | | A > del 1 |
| NC_014760 | 572484 | MBOVPG45_RS04635 | | | C>T |
| NC_014760 | 572490 | MBOVPG45_RS04635 | | | - > ins T |
| NC_014760 | 573609 | MBOVPG45_RS02470 | | | T>G |

| | | | | | |
|---|---|---|---|---|---|
| NC_014760 | 573793 | MBOVPG45_RS02470 | | | C>T |
| NC_014760 | 573795 | MBOVPG45_RS02470 | | | T>A |
| NC_014760 | 573840 | MBOVPG45_RS02470 | | | T>G |
| NC_014760 | 573886 | MBOVPG45_RS02470 | | | T>C |
| NC_014760 | 573955 | MBOVPG45_RS02470 | | | T>C |
| NC_014760 | 573989 | MBOVPG45_RS02470 | | | T>A |
| NC_014760 | 574024 | MBOVPG45_RS02470 | | | G>A |
| NC_014760 | 574026 | MBOVPG45_RS02470 | | | A>G |
| NC_014760 | 574046 | MBOVPG45_RS02470 | | | A>C |
| NC_014760 | 575129 | MBOVPG45_RS02480 | | | A>T |
| NC_014760 | 575865 | MBOVPG45_RS02480 | | A > del 1 | |
| NC_014760 | 575881 | MBOVPG45_RS02480 | | | A>G |
| NC_014760 | 575893 | MBOVPG45_RS02480 | | | G>C |
| NC_014760 | 576731 | MBOVPG45_RS02485 | | | A>G |
| NC_014760 | 576734 | MBOVPG45_RS02485 | | | T>C |
| NC_014760 | 576925 | MBOVPG45_RS02485 | | | - > ins T |
| NC_014760 | 577264 | MBOVPG45_RS02490 | | | TCC>CTT |
| NC_014760 | 577275 | MBOVPG45_RS02490 | | | AA>GG |
| NC_014760 | 577278 | MBOVPG45_RS02490 | | | C>T |
| NC_014760 | 577312 | MBOVPG45_RS02490 | | | C>T |
| NC_014760 | 578011 | MBOVPG45_RS02495 | | | C>T |
| NC_014760 | 578792 | MBOVPG45_RS02500 | | T>C | |
| NC_014760 | 578841 | MBOVPG45_RS02500 | | | GT>AC |
| NC_014760 | 578874 | MBOVPG45_RS02500 | | | T>G |
| NC_014760 | 579359 | MBOVPG45_RS02500 | | | T>C |
| NC_014760 | 579372 | MBOVPG45_RS02500 | | | - > ins GAT |
| NC_014760 | 579400 | MBOVPG45_RS02500 | | | T>C |
| NC_014760 | 579404 | MBOVPG45_RS02500 | | | G>A |
| NC_014760 | 579484 | MBOVPG45_RS02500 | | | G>T |
| NC_014760 | 579487 | MBOVPG45_RS02500 | | | C>T |
| NC_014760 | 579518 | MBOVPG45_RS02500 | | | TT>CC |
| NC_014760 | 579529 | MBOVPG45_RS02500 | | | TC>GT |
| NC_014760 | 579617 | MBOVPG45_RS02500 | | | G>A |
| NC_014760 | 579619 | MBOVPG45_RS02500 | | | A>T |
| NC_014760 | 579727 | MBOVPG45_RS02500 | | | C>T |
| NC_014760 | 579805 | MBOVPG45_RS02500 | | | CC>TT |
| NC_014760 | 579830 | MBOVPG45_RS02500 | | | T>C |
| NC_014760 | 579857 | MBOVPG45_RS02500 | | | C>T |
| NC_014760 | 579901 | MBOVPG45_RS02500 | | | TG>CC |
| NC_014760 | 579904 | MBOVPG45_RS02500 | | | T>A |
| NC_014760 | 579911 | MBOVPG45_RS02500 | | | T>C |
| NC_014760 | 579925 | MBOVPG45_RS02500 | | | C>T |
| NC_014760 | 579928 | MBOVPG45_RS02500 | | | T>A |
| NC_014760 | 579931 | MBOVPG45_RS02500 | | | TAT > del 3 |
| NC_014760 | 579971 | MBOVPG45_RS02500 | | | C>T |
| NC_014760 | 580022 | MBOVPG45_RS02500 | | | A>G |
| NC_014760 | 580026 | MBOVPG45_RS02500 | | | T>A |

| | | | | | | |
|---|---|---|---|---|---|---|
| NC_014760 | 580032 | MBOVPG45_RS02500 | | | | - > ins TTT |
| NC_014760 | 580084 | MBOVPG45_RS02500 | | | | C>T |
| NC_014760 | 580095 | MBOVPG45_RS02500 | | | | TGATATATT > del 9 |
| NC_014760 | 580303 | MBOVPG45_RS02500 | | | | C>G |
| NC_014760 | 580438 | MBOVPG45_RS02500 | | | | G>T |
| NC_014760 | 580594 | MBOVPG45_RS02500 | | | | G>C |
| NC_014760 | 580619 | MBOVPG45_RS02500 | | | | T>C |
| NC_014760 | 580685 | MBOVPG45_RS02500 | | | | T>C |
| NC_014760 | 580780 | MBOVPG45_RS02500 | | | | T>C |
| NC_014760 | 580897 | MBOVPG45_RS02500 | | | | A>G |
| NC_014760 | 580900 | MBOVPG45_RS02500 | | | | G>A |
| NC_014760 | 581268 | MBOVPG45_RS02505 | | | | A>T |
| NC_014760 | 581494 | MBOVPG45_RS02505 | | | | T>A |
| NC_014760 | 581561 | MBOVPG45_RS02505 | | | | G>A |
| NC_014760 | 581582 | MBOVPG45_RS02505 | | | | A>G |
| NC_014760 | 581597 | MBOVPG45_RS02505 | | | | T>C |
| NC_014760 | 581712 | MBOVPG45_RS02505 | | | | A>G |
| NC_014760 | 581846 | MBOVPG45_RS02505 | | | | G>A |
| NC_014760 | 582294 | ruvX | | | | T>C |
| NC_014760 | 582733 | alaS | | | | G>C |
| NC_014760 | 583598 | alaS | | | | T>C |
| NC_014760 | 583645 | alaS | | | | G>A |
| NC_014760 | 583801 | alaS | | | | C>T |
| NC_014760 | 584000 | alaS | | | | C>T |
| NC_014760 | 584052 | alaS | | | | T>A |
| NC_014760 | 584117 | alaS | | | G>A | |
| NC_014760 | 584246 | alaS | | | | T>C |
| NC_014760 | 584257 | alaS | | | | C>G |
| NC_014760 | 585222 | mnmA | | | | T>C |
| NC_014760 | 585273 | mnmA | | | | T>C |
| NC_014760 | 585314 | mnmA | | | | G>A |
| NC_014760 | 585459 | mnmA | | | | T>C |
| NC_014760 | 585469 | mnmA | | | | A>T |
| NC_014760 | 586464 | MBOVPG45_RS02525 | | | | T>C |
| NC_014760 | 586640 | MBOVPG45_RS02525 | | | | G>A |
| NC_014760 | 587053 | MBOVPG45_RS02525 | | | | GC>AT |
| NC_014760 | 587200 | MBOVPG45_RS02530 | | | | A>G |
| NC_014760 | 587419 | MBOVPG45_RS02530 | | | | A>G |
| NC_014760 | 587434 | MBOVPG45_RS02530 | | | | A>T |
| NC_014760 | 587476 | MBOVPG45_RS02530 | | | | G>A |
| NC_014760 | 587523 | MBOVPG45_RS02530 | | | | A>C |
| NC_014760 | 587536 | MBOVPG45_RS02530 | | | | G>A |
| NC_014760 | 587647 | MBOVPG45_RS02530 | | | | C>A |
| NC_014760 | 587660 | MBOVPG45_RS02530 | | | | TC>CT |
| NC_014760 | 587663 | MBOVPG45_RS02530 | | | | G>A |
| NC_014760 | 587683 | MBOVPG45_RS02530 | | | | GAG>AGC |

| | | | | | |
|---|---|---|---|---|---|
| NC_014760 | 591273 | MBOVPG45_RS02550 | | | G>T |
| NC_014760 | 591350 | MBOVPG45_RS02550 | | | C>A |
| NC_014760 | 591570 | MBOVPG45_RS02550 | | | T>A |
| NC_014760 | 592036 | MBOVPG45_RS02550 | | | C>T |
| NC_014760 | 592099 | MBOVPG45_RS02550 | | | C>T |
| NC_014760 | 592126 | MBOVPG45_RS02550 | | | C>T |
| NC_014760 | 592432 | MBOVPG45_RS02555 | | | A>T |
| NC_014760 | 592516 | MBOVPG45_RS02555 | | | G>A |
| NC_014760 | 592526 | MBOVPG45_RS02555 | | | T>C |
| NC_014760 | 592580 | MBOVPG45_RS02555 | | | T>C |
| NC_014760 | 592587 | MBOVPG45_RS02555 | | | - > ins G |
| NC_014760 | 592588 | MBOVPG45_RS02555 | | | A>T |
| NC_014760 | 592642 | MBOVPG45_RS02555 | | | G>A |
| NC_014760 | 592645 | MBOVPG45_RS02555 | | | T>C |
| NC_014760 | 592676 | MBOVPG45_RS02555 | | | C>T |
| NC_014760 | 592985 | MBOVPG45_RS02555 | | | T>A |
| NC_014760 | 593925 | MBOVPG45_RS02560 | | | C>T |
| NC_014760 | 594709 | MBOVPG45_RS02565 | | | G>A |
| NC_014760 | 594792 | MBOVPG45_RS02565 | | | G>C |
| NC_014760 | 594891 | MBOVPG45_RS02565 | | | CG>TA |
| NC_014760 | 595603 | MBOVPG45_RS02570 | | | T>C |
| NC_014760 | 595806 | MBOVPG45_RS02570 | | | T>A |
| NC_014760 | 595822 | MBOVPG45_RS02570 | | | G>T |
| NC_014760 | 596087 | MBOVPG45_RS02570 | | | T>G |
| NC_014760 | 596888 | MBOVPG45_RS02575 | | | CA>AG |
| NC_014760 | 596973 | MBOVPG45_RS02575 | | | G>A |
| NC_014760 | 597195 | MBOVPG45_RS02575 | | | A>G |
| NC_014760 | 597675 | MBOVPG45_RS02580 | | | T>C |
| NC_014760 | 597818 | MBOVPG45_RS02580 | | | C>T |
| NC_014760 | 598367 | MBOVPG45_RS02580 | | | G>A |
| NC_014760 | 598389 | MBOVPG45_RS02580 | | | AC>GT |
| NC_014760 | 600549 | MBOVPG45_RS02580 | | | C>T |
| NC_014760 | 606351 | MBOVPG45_RS02615 | | | G>A |
| NC_014760 | 609179 | glpK | | | G>A |
| NC_014760 | 609190 | glpK | | | G>A |
| NC_014760 | 610095 | glpK | | | A>T |
| NC_014760 | 613725 | MBOVPG45_RS02640 | | | T>C |
| NC_014760 | 615303 | MBOVPG45_RS02645 | | | C>T |
| NC_014760 | 615566 | MBOVPG45_RS02645 | | | G>A |
| NC_014760 | 620753 | MBOVPG45_RS02665 | | | C>G |
| NC_014760 | 620756 | MBOVPG45_RS02665 | | | C>T |
| NC_014760 | 624992 | topA | | | - > ins A |
| NC_014760 | 631133 | MBOVPG45_RS02715, MBOVPG45_RS02710 | | | C>T |
| NC_014760 | 631191 | MBOVPG45_RS02715 | | | G>A |
| NC_014760 | 631213 | MBOVPG45_RS02715 | | | T>C |
| NC_014760 | 633285 | MBOVPG45_RS02720 | | | C>T |

| | | | | | |
|---|---|---|---|---|---|
| NC_014760 | 634259 | MBOVPG45_RS02725 | | | C>T |
| NC_014760 | 634299 | MBOVPG45_RS02725 | | | A>G |
| NC_014760 | 634813 | MBOVPG45_RS02725 | | | A>G |
| NC_014760 | 635026 | MBOVPG45_RS02725 | | | G>A |
| NC_014760 | 635117 | MBOVPG45_RS02725 | | | C>T |
| NC_014760 | 635135 | MBOVPG45_RS02725 | | | CTGAGC > del 6 |
| NC_014760 | 635152 | MBOVPG45_RS02725 | | | A>G |
| NC_014760 | 635163 | MBOVPG45_RS02725 | | | T>A |
| NC_014760 | 635185 | MBOVPG45_RS02725 | | | G>C |
| NC_014760 | 635197 | MBOVPG45_RS02725 | | | T>C |
| NC_014760 | 635200 | MBOVPG45_RS02725 | | | A>G |
| NC_014760 | 635267 | MBOVPG45_RS02725 | | | T>A |
| NC_014760 | 635282 | MBOVPG45_RS02725 | | | C>T |
| NC_014760 | 635285 | MBOVPG45_RS02725 | | | C>T |
| NC_014760 | 635295 | MBOVPG45_RS02725 | | | - > ins GAT |
| NC_014760 | 635342 | MBOVPG45_RS02725 | | | C>T |
| NC_014760 | 635463 | MBOVPG45_RS02725 | | | GC>AT |
| NC_014760 | 635566 | MBOVPG45_RS02725 | | | C>A |
| NC_014760 | 635575 | MBOVPG45_RS02725 | | | G>A |
| NC_014760 | 636688 | MBOVPG45_RS02730 | | | G>A |
| NC_014760 | 636732 | MBOVPG45_RS02730 | | | A>G |
| NC_014760 | 637373 | dnaG | | | G>T |
| NC_014760 | 637455 | dnaG | | | A>G |
| NC_014760 | 638540 | dnaG | | | G>A |
| NC_014760 | 638893 | dnaG | | | G>A |
| NC_014760 | 639516 | MBOVPG45_RS02740 | | | T>A |
| NC_014760 | 641166 | MBOVPG45_RS02745 | | | T>C |
| NC_014760 | 641193 | MBOVPG45_RS02745 | | | A>G |
| NC_014760 | 641354 | MBOVPG45_RS02745 | | | AC>CA |
| NC_014760 | 641390 | MBOVPG45_RS02745 | | | T>G |
| NC_014760 | 641833 | MBOVPG45_RS02750 | | | A>G |
| NC_014760 | 641914 | MBOVPG45_RS02750 | | | G>A |
| NC_014760 | 642412 | MBOVPG45_RS02750 | | | A>G |
| NC_014760 | 643726 | MBOVPG45_RS02755 | | | A>G |
| NC_014760 | 643759 | MBOVPG45_RS02760 | | | A>G |
| NC_014760 | 643790 | MBOVPG45_RS02760 | | | A>T |
| NC_014760 | 644166 | MBOVPG45_RS02760 | | | G>A |
| NC_014760 | 644211 | MBOVPG45_RS02760 | | | G>A |
| NC_014760 | 644240 | MBOVPG45_RS02760 | | | G>A |
| NC_014760 | 644253 | MBOVPG45_RS02760 | | | A>G |
| NC_014760 | 644774 | MBOVPG45_RS02760 | | | A>G |
| NC_014760 | 644777 | MBOVPG45_RS02760 | | | T>C |
| NC_014760 | 644786 | MBOVPG45_RS02760 | | | G>A |
| NC_014760 | 644808 | MBOVPG45_RS02760 | | | A>G |
| NC_014760 | 644819 | MBOVPG45_RS02760 | | | T>A |
| NC_014760 | 644849 | MBOVPG45_RS02760 | | | G>A |
| NC_014760 | 644862 | MBOVPG45_RS02760 | | | G>A |

| | | | | | |
|---|---|---|---|---|---|
| NC_014760 | 644865 | MBOVPG45_RS02760 | | | A>T |
| NC_014760 | 644909 | MBOVPG45_RS02760 | | | T>C |
| NC_014760 | 644959 | MBOVPG45_RS02760 | | | G>T |
| NC_014760 | 645099 | MBOVPG45_RS02760 | | | T>A |
| NC_014760 | 645310 | MBOVPG45_RS02760 | | | A>G |
| NC_014760 | 645973 | MBOVPG45_RS02765 | | | C>T |
| NC_014760 | 646189 | MBOVPG45_RS02765 | | | T>G |
| NC_014760 | 646219 | MBOVPG45_RS02765 | | | C>T |
| NC_014760 | 646365 | MBOVPG45_RS02765 | | | C>T |
| NC_014760 | 646369 | MBOVPG45_RS02765 | | | T>C |
| NC_014760 | 646646 | rpsB | | | T>C |
| NC_014760 | 647105 | rpsB | | | A>C |
| NC_014760 | 647511 | rpsB | | | C>T |
| NC_014760 | 647839 | MBOVPG45_RS02775 | | | T>C |
| NC_014760 | 648784 | MBOVPG45_RS02780 | | | T>A |
| NC_014760 | 648875 | MBOVPG45_RS02780 | | | C>A |
| NC_014760 | 648893 | MBOVPG45_RS02780 | | | T>C |
| NC_014760 | 649000 | MBOVPG45_RS02780 | | | A>G |
| NC_014760 | 649011 | MBOVPG45_RS02780 | | | TG>CA |
| NC_014760 | 649015 | MBOVPG45_RS02780 | | | C>T |
| NC_014760 | 649103 | MBOVPG45_RS02780 | | | G>A |
| NC_014760 | 649213 | MBOVPG45_RS02780 | | | T>A |
| NC_014760 | 649254 | MBOVPG45_RS02780 | | | AA>GG |
| NC_014760 | 649358 | MBOVPG45_RS02780 | | | T>C |
| NC_014760 | 649427 | MBOVPG45_RS02780 | | | C>T |
| NC_014760 | 649462 | MBOVPG45_RS02780 | | | T>G |
| NC_014760 | 649516 | MBOVPG45_RS02780 | | | G>A |
| NC_014760 | 650147 | MBOVPG45_RS02785 | | | TAATAGTAGTAG > del 12 |
| NC_014760 | 650149 | MBOVPG45_RS02785 | | - > ins TAGx3 | |
| NC_014760 | 650172 | MBOVPG45_RS02785 | | | T>C |
| NC_014760 | 650184 | MBOVPG45_RS02785 | | | A>G |
| NC_014760 | 650209 | MBOVPG45_RS02785 | | | GC>AT |
| NC_014760 | 650212 | MBOVPG45_RS02785 | | | C>T |
| NC_014760 | 650235 | MBOVPG45_RS02785 | | | G>A |
| NC_014760 | 650298 | MBOVPG45_RS02785 | | | C>G |
| NC_014760 | 650424 | MBOVPG45_RS02785 | | | T>C |
| NC_014760 | 650433 | MBOVPG45_RS02785 | | | A>G |
| NC_014760 | 650481 | MBOVPG45_RS02785 | | | A>T |
| NC_014760 | 650512 | MBOVPG45_RS02785 | | | C>T |
| NC_014760 | 650561 | MBOVPG45_RS02785 | | | CG>TT |
| NC_014760 | 650886 | MBOVPG45_RS02785 | | | A>G |
| NC_014760 | 651207 | MBOVPG45_RS02785 | | | A>G |
| NC_014760 | 651209 | MBOVPG45_RS02785 | | | AGA>GAC |
| NC_014760 | 651219 | MBOVPG45_RS02785 | | | AA>GG |
| NC_014760 | 651245 | MBOVPG45_RS02785 | | | A>T |
| NC_014760 | 651310 | MBOVPG45_RS02785 | | | G>A |

| | | | | | |
|---|---|---|---|---|---|
| NC_014760 | 651341 | MBOVPG45_RS02785 | | | T>A |
| NC_014760 | 651369 | MBOVPG45_RS02785 | | | GT>AC |
| NC_014760 | 651372 | MBOVPG45_RS02785 | | | C>A |
| NC_014760 | 651376 | MBOVPG45_RS02785 | | | GG>AC |
| NC_014760 | 651392 | MBOVPG45_RS02785 | | | T>A |
| NC_014760 | 651430 | MBOVPG45_RS02790 | | | G>A |
| NC_014760 | 651447 | MBOVPG45_RS02790 | | | T>G |
| NC_014760 | 651475 | MBOVPG45_RS02790 | | | T>A |
| NC_014760 | 651498 | MBOVPG45_RS02790 | | | G>A |
| NC_014760 | 651511 | MBOVPG45_RS02790 | | | G>T |
| NC_014760 | 651633 | MBOVPG45_RS02790 | | | A > del 1 |
| NC_014760 | 651781 | MBOVPG45_RS02795 | | | T>C |
| NC_014760 | 651795 | MBOVPG45_RS02795 | | | C>A |
| NC_014760 | 651932 | MBOVPG45_RS02795 | | | G>A |
| NC_014760 | 651993 | MBOVPG45_RS02795 | | | C>A |
| NC_014760 | 652512 | MBOVPG45_RS02795 | | | G>A |
| NC_014760 | 653132 | MBOVPG45_RS02795 | | | A>G |
| NC_014760 | 653165 | MBOVPG45_RS02795 | | | GA>AC |
| NC_014760 | 653616 | MBOVPG45_RS02795 | | | A>G |
| NC_014760 | 653669 | MBOVPG45_RS02795 | | | A>T |
| NC_014760 | 654099 | MBOVPG45_RS02800 | | | G>C |
| NC_014760 | 654116 | MBOVPG45_RS02800 | | | - > ins CAAAC |
| NC_014760 | 654118 | MBOVPG45_RS02800 | | | - > ins A |
| NC_014760 | 655861 | MBOVPG45_RS02800, MBOVPG45_RS02805 | | | C>T |
| NC_014760 | 655873 | MBOVPG45_RS02805, MBOVPG45_RS02800 | | | GC>AA |
| NC_014760 | 656135 | MBOVPG45_RS02805 | | | G>A |
| NC_014760 | 656339 | MBOVPG45_RS02805 | | | G>A |
| NC_014760 | 656384 | MBOVPG45_RS02805 | | | A>G |
| NC_014760 | 656401 | MBOVPG45_RS02805 | | | A>G |
| NC_014760 | 656403 | MBOVPG45_RS02805 | | | G>T |
| NC_014760 | 656662 | MBOVPG45_RS02805 | | | G>A |
| NC_014760 | 656769 | MBOVPG45_RS02805 | | | C>G |
| NC_014760 | 656957 | MBOVPG45_RS02805 | | | A>G |
| NC_014760 | 657141 | MBOVPG45_RS02805 | | | AA>GG |
| NC_014760 | 657334 | MBOVPG45_RS02805 | | | A>G |
| NC_014760 | 657487 | MBOVPG45_RS02805 | | | C>G |
| NC_014760 | 657550 | MBOVPG45_RS02805 | | | A>G |
| NC_014760 | 657724 | MBOVPG45_RS02805 | | | G>A |
| NC_014760 | 659437 | rmuC | | | G>A |
| NC_014760 | 659637 | MBOVPG45_RS02815 | | | G>A |
| NC_014760 | 659676 | MBOVPG45_RS02815 | | | T>C |
| NC_014760 | 662114 | MBOVPG45_RS02820 | | G>A | |
| NC_014760 | 663026 | MBOVPG45_RS02825 | | | G>A |
| NC_014760 | 663029 | MBOVPG45_RS02825 | | | G>C |
| NC_014760 | 666001 | MBOVPG45_RS02835 | | | G>A |

| | | | | | |
|---|---|---|---|---|---|
| NC_014760 | 666071 | MBOVPG45_RS02835 | | C>A | |
| NC_014760 | 666194 | MBOVPG45_RS02835 | | | T>C |
| NC_014760 | 666558 | MBOVPG45_RS02835 | | | - > ins T |
| NC_014760 | 667694 | MBOVPG45_RS02845 | | T > del 1 | |
| NC_014760 | 669998 | MBOVPG45_RS02855 | | | A>G |
| NC_014760 | 670440 | MBOVPG45_RS02860 | | | T>C |
| NC_014760 | 670483 | MBOVPG45_RS02860 | | | T>C |
| NC_014760 | 670565 | MBOVPG45_RS02860 | | | A>T |
| NC_014760 | 671196 | MBOVPG45_RS02860 | | | A>T |
| NC_014760 | 671422 | MBOVPG45_RS02860 | | | C>T |
| NC_014760 | 674943 | dnaB | | | T>C |
| NC_014760 | 677174 | MBOVPG45_RS02885 | | | T>A |
| NC_014760 | 677211 | MBOVPG45_RS02885 | | | T>C |
| NC_014760 | 677282 | MBOVPG45_RS02885 | | | T>C |
| NC_014760 | 677565 | MBOVPG45_RS02885 | | | G>T |
| NC_014760 | 677990 | MBOVPG45_RS02890 | | | T>A |
| NC_014760 | 678204 | MBOVPG45_RS02890 | | | T>A |
| NC_014760 | 678237 | MBOVPG45_RS02890 | | | G>A |
| NC_014760 | 678361 | MBOVPG45_RS02890 | | | T>C |
| NC_014760 | 678373 | MBOVPG45_RS02890 | | | T>C |
| NC_014760 | 678403 | MBOVPG45_RS02890 | | | G>A |
| NC_014760 | 678405 | MBOVPG45_RS02890 | | | TT>AC |
| NC_014760 | 678475 | MBOVPG45_RS02890 | | | T>C |
| NC_014760 | 678491 | MBOVPG45_RS02890 | | | T>G |
| NC_014760 | 678702 | MBOVPG45_RS02890 | | | G>A |
| NC_014760 | 678763 | MBOVPG45_RS02890 | | | A>G |
| NC_014760 | 678813 | MBOVPG45_RS02890 | | | A>G |
| NC_014760 | 678846 | MBOVPG45_RS02890 | | | A>G |
| NC_014760 | 678882 | MBOVPG45_RS02890 | | | G>A |
| NC_014760 | 679145 | MBOVPG45_RS02895 | | | T>C |
| NC_014760 | 679603 | MBOVPG45_RS02895 | | | C>T |
| NC_014760 | 680552 | MBOVPG45_RS02895 | | | A>G |
| NC_014760 | 681045 | MBOVPG45_RS02900 | | | T>G |
| NC_014760 | 681061 | MBOVPG45_RS02900 | | | T>G |
| NC_014760 | 681068 | MBOVPG45_RS02900 | | | GTT>ACC |
| NC_014760 | 681212 | MBOVPG45_RS02900 | | | A>T |
| NC_014760 | 681239 | MBOVPG45_RS02900 | | | T>A |
| NC_014760 | 681260 | MBOVPG45_RS02900 | | | A>T |
| NC_014760 | 681271 | MBOVPG45_RS02900 | | | T>C |
| NC_014760 | 681396 | MBOVPG45_RS02900 | | | A>G |
| NC_014760 | 681477 | MBOVPG45_RS02900 | | | C>T |
| NC_014760 | 681563 | MBOVPG45_RS02900 | | | T>A |
| NC_014760 | 681565 | MBOVPG45_RS02900 | | | G>A |
| NC_014760 | 681567 | MBOVPG45_RS02900 | | | C>T |
| NC_014760 | 681724 | MBOVPG45_RS02900 | | | G>A |
| NC_014760 | 681754 | MBOVPG45_RS02900 | | | T>C |
| NC_014760 | 681765 | MBOVPG45_RS02900 | | | A>G |

| | | | | | |
|---|---|---|---|---|---|
| NC_014760 | 681816 | MBOVPG45_RS02900 | | | G>T |
| NC_014760 | 681947 | MBOVPG45_RS02900 | | | A>C |
| NC_014760 | 681988 | MBOVPG45_RS02900 | | | G>A |
| NC_014760 | 682057 | MBOVPG45_RS02900 | | | C>T |
| NC_014760 | 682061 | MBOVPG45_RS02900 | | | AG>GA |
| NC_014760 | 682066 | MBOVPG45_RS02900 | | | T>C |
| NC_014760 | 682073 | MBOVPG45_RS02900 | | | A>T |
| NC_014760 | 682131 | MBOVPG45_RS02900 | | | T>C |
| NC_014760 | 682153 | MBOVPG45_RS02900 | | | T>G |
| NC_014760 | 682161 | MBOVPG45_RS02900 | | | TC>GT |
| NC_014760 | 682166 | MBOVPG45_RS02900 | | | A>C |
| NC_014760 | 682178 | MBOVPG45_RS02900 | | | C>A |
| NC_014760 | 682191 | MBOVPG45_RS02900 | | | GC>TT |
| NC_014760 | 682360 | MBOVPG45_RS02900 | | | G>A |
| NC_014760 | 682496 | MBOVPG45_RS02900 | | | A>T |
| NC_014760 | 682552 | MBOVPG45_RS02900 | | | C>G |
| NC_014760 | 682614 | MBOVPG45_RS02900 | | | T>G |
| NC_014760 | 682619 | MBOVPG45_RS02900 | | | ATCT>TGAC |
| NC_014760 | 682689 | MBOVPG45_RS02900 | | | G>A |
| NC_014760 | 682961 | MBOVPG45_RS02905 | | C>T | |
| NC_014760 | 683395 | MBOVPG45_RS02905 | | | T>C |
| NC_014760 | 690881 | MBOVPG45_RS02945 | | A>C | |
| NC_014760 | 693602 | ruvA | | | A>G |
| NC_014760 | 693690 | ruvA | | | T>C |
| NC_014760 | 697337 | MBOVPG45_RS02990 | | | T>C |
| NC_014760 | 697407 | MBOVPG45_RS02990 | | | T>C |
| NC_014760 | 697545 | MBOVPG45_RS02990 | | | G>A |
| NC_014760 | 697668 | MBOVPG45_RS02990 | | | T>C |
| NC_014760 | 697741 | ileS | | | T > del 1 |
| NC_014760 | 697989 | ileS | | A>G | |
| NC_014760 | 698895 | ileS | | | A>G |
| NC_014760 | 699583 | ileS | | | T>C |
| NC_014760 | 699621 | ileS | | | TG>CA |
| NC_014760 | 700006 | ileS | | | C>T |
| NC_014760 | 700827 | MBOVPG45_RS03000 | | | G>C |
| NC_014760 | 701457 | MBOVPG45_RS03000 | | | A>G |
| NC_014760 | 704332 | MBOVPG45_RS03050 | | | G>A |
| NC_014760 | 704335 | MBOVPG45_RS03050 | | | G>C |
| NC_014760 | 713717 | MBOVPG45_RS03080 | | | C>T |
| NC_014760 | 713934 | MBOVPG45_RS03080 | | | T>A |
| NC_014760 | 714191 | MBOVPG45_RS03080 | | | T>G |
| NC_014760 | 714378 | MBOVPG45_RS03080 | | | AC>TT |
| NC_014760 | 714566 | MBOVPG45_RS03080 | | | T>C |
| NC_014760 | 714581 | MBOVPG45_RS03080 | | | T>C |
| NC_014760 | 715138 | MBOVPG45_RS03080 | | | T>A |
| NC_014760 | 715144 | MBOVPG45_RS03080 | | | T>A |
| NC_014760 | 715412 | MBOVPG45_RS03080 | | | T>C |

| | | | | | |
|---|---|---|---|---|---|
| NC_014760 | 715692 | MBOVPG45_RS03080 | | | A>T |
| NC_014760 | 716234 | MBOVPG45_RS03085 | | | T>C |
| NC_014760 | 717489 | MBOVPG45_RS03090 | | | G>A |
| NC_014760 | 719249 | MBOVPG45_RS03095 | | | A>C |
| NC_014760 | 719322 | MBOVPG45_RS03095 | | | GG>CA |
| NC_014760 | 719326 | MBOVPG45_RS03095 | | | GT>AC |
| NC_014760 | 719339 | MBOVPG45_RS03095 | | | C>T |
| NC_014760 | 719348 | MBOVPG45_RS03095 | | | T>A |
| NC_014760 | 719402 | MBOVPG45_RS03095 | | | A>G |
| NC_014760 | 719728 | MBOVPG45_RS03095 | | | G>A |
| NC_014760 | 730969 | MBOVPG45_RS03145 | | | A>G |
| NC_014760 | 731350 | MBOVPG45_RS03150 | | | G>A |
| NC_014760 | 731400 | MBOVPG45_RS03150 | | | A>T |
| NC_014760 | 731735 | MBOVPG45_RS03150 | | | GT>AC |
| NC_014760 | 731749 | MBOVPG45_RS03150 | | | A>G |
| NC_014760 | 732255 | MBOVPG45_RS03150 | | | G>A |
| NC_014760 | 732335 | MBOVPG45_RS03150 | | | AA>GC |
| NC_014760 | 732418 | MBOVPG45_RS03150 | | | G>A |
| NC_014760 | 732431 | MBOVPG45_RS03150 | | | G>C |
| NC_014760 | 732530 | MBOVPG45_RS03150 | | | A>G |
| NC_014760 | 732536 | MBOVPG45_RS03150 | | | G>A |
| NC_014760 | 732728 | MBOVPG45_RS03155 | | | C>T |
| NC_014760 | 733123 | ftsY | | - > ins AAGATG | - > ins AAGATG |
| NC_014760 | 733682 | ftsY | | | A>T |
| NC_014760 | 735556 | MBOVPG45_RS03180 | | | T>C |
| NC_014760 | 735776 | MBOVPG45_RS03180 | | | C>T |
| NC_014760 | 735791 | MBOVPG45_RS03180 | | | C>T |
| NC_014760 | 735844 | MBOVPG45_RS03180 | | | G>A |
| NC_014760 | 736022 | MBOVPG45_RS03180 | | | T>G |
| NC_014760 | 736034 | MBOVPG45_RS03180 | | | CG>TA |
| NC_014760 | 736037 | MBOVPG45_RS03180 | | | A>C |
| NC_014760 | 736051 | MBOVPG45_RS03180 | | | T>C |
| NC_014760 | 736179 | MBOVPG45_RS03180 | | | T>C |
| NC_014760 | 736184 | MBOVPG45_RS03180 | | | T>A |
| NC_014760 | 736186 | MBOVPG45_RS03180 | | | C>A |
| NC_014760 | 736512 | MBOVPG45_RS03185 | | | T>C |
| NC_014760 | 736530 | MBOVPG45_RS03185 | | | C>T |
| NC_014760 | 736781 | MBOVPG45_RS03190 | | | C>A |
| NC_014760 | 736814 | MBOVPG45_RS03190 | | | C>T |
| NC_014760 | 736820 | MBOVPG45_RS03190 | | | TG>AT |
| NC_014760 | 736867 | MBOVPG45_RS03190 | | | T>A |
| NC_014760 | 736878 | MBOVPG45_RS03190 | | | A>C |
| NC_014760 | 737001 | MBOVPG45_RS03190 | | | GCC > del 3 |
| NC_014760 | 737009 | MBOVPG45_RS03190 | | | G>A |
| NC_014760 | 737011 | MBOVPG45_RS03190 | | | TT>GC |
| NC_014760 | 737122 | MBOVPG45_RS03190 | | | C>G |
| NC_014760 | 737173 | MBOVPG45_RS03190 | | | G>T |

| | | | | | |
|---|---|---|---|---|---|
| NC_014760 | 737250 | MBOVPG45_RS03190 | | | C>G |
| NC_014760 | 737282 | MBOVPG45_RS03190 | | | C>T |
| NC_014760 | 737284 | MBOVPG45_RS03190 | | | T>C |
| NC_014760 | 737294 | MBOVPG45_RS03190 | | | G>C |
| NC_014760 | 737384 | MBOVPG45_RS03190 | | | T>C |
| NC_014760 | 737514 | MBOVPG45_RS03190 | | | C>A |
| NC_014760 | 737603 | MBOVPG45_RS03190 | | | T>G |
| NC_014760 | 737642 | MBOVPG45_RS03190 | | | C>T |
| NC_014760 | 737818 | MBOVPG45_RS03190 | | | A>T |
| NC_014760 | 737821 | MBOVPG45_RS03190 | | | A>G |
| NC_014760 | 737874 | MBOVPG45_RS03190 | | | G>T |
| NC_014760 | 737897 | MBOVPG45_RS03190 | | | C>T |
| NC_014760 | 738079 | MBOVPG45_RS03195 | | | C>G |
| NC_014760 | 738200 | MBOVPG45_RS03195 | | | C>T |
| NC_014760 | 738205 | MBOVPG45_RS03195 | | | C>T |
| NC_014760 | 738274 | MBOVPG45_RS03195 | | | C>T |
| NC_014760 | 738391 | MBOVPG45_RS03195 | | | C>G |
| NC_014760 | 738521 | MBOVPG45_RS03195 | | | G>T |
| NC_014760 | 738541 | MBOVPG45_RS03195 | | | T>C |
| NC_014760 | 738548 | MBOVPG45_RS03195 | | | C>T |
| NC_014760 | 738584 | MBOVPG45_RS03195 | | | CC>TG |
| NC_014760 | 738601 | MBOVPG45_RS03195 | | | - > ins AAT |
| NC_014760 | 738605 | MBOVPG45_RS03195 | | | GC>AT |
| NC_014760 | 738614 | MBOVPG45_RS03195 | | | GTTATC > del 6 |
| NC_014760 | 738692 | MBOVPG45_RS03195 | | | G>T |
| NC_014760 | 738965 | MBOVPG45_RS03195 | | | A>T |
| NC_014760 | 738990 | MBOVPG45_RS03195 | | | C>T |
| NC_014760 | 739076 | MBOVPG45_RS03195 | | | A>T |
| NC_014760 | 739142 | MBOVPG45_RS03195 | | | A>T |
| NC_014760 | 739153 | MBOVPG45_RS03195 | | | GA>CC |
| NC_014760 | 739158 | MBOVPG45_RS03195 | | | C>T |
| NC_014760 | 739252 | MBOVPG45_RS03195 | | | T>C |
| NC_014760 | 739479 | MBOVPG45_RS03195 | | | C>T |
| NC_014760 | 739484 | MBOVPG45_RS03195 | | | T>A |
| NC_014760 | 739487 | MBOVPG45_RS03195 | | | T>G |
| NC_014760 | 739506 | MBOVPG45_RS03195 | | | T>C |
| NC_014760 | 739569 | MBOVPG45_RS03195 | | | G>A |
| NC_014760 | 739606 | MBOVPG45_RS03195 | | | G>A |
| NC_014760 | 740336 | MBOVPG45_RS03200 | | | G>A |
| NC_014760 | 740345 | MBOVPG45_RS03200 | | | G>A |
| NC_014760 | 740525 | MBOVPG45_RS03200 | | | A>C |
| NC_014760 | 740780 | MBOVPG45_RS03200 | | | G>A |
| NC_014760 | 741395 | MBOVPG45_RS03210 | | | A>G |
| NC_014760 | 743052 | MBOVPG45_RS03215 | | | A>C |
| NC_014760 | 743059 | MBOVPG45_RS03215 | | | A>G |
| NC_014760 | 743453 | MBOVPG45_RS03215 | | | CA>TG |
| NC_014760 | 743837 | MBOVPG45_RS03215 | | | A>T |

| | | | | | |
|---|---|---|---|---|---|
| NC_014760 | 744092 | MBOVPG45_RS03220 | | | A>G |
| NC_014760 | 744495 | MBOVPG45_RS03220 | | | A>G |
| NC_014760 | 744536 | MBOVPG45_RS03220 | | | A>G |
| NC_014760 | 744545 | MBOVPG45_RS03220 | | | C>T |
| NC_014760 | 744726 | MBOVPG45_RS03220 | | | G>A |
| NC_014760 | 744739 | MBOVPG45_RS03220 | | | T>A |
| NC_014760 | 744754 | MBOVPG45_RS03220 | | | G>T |
| NC_014760 | 745757 | MBOVPG45_RS03225 | | | AC>GT |
| NC_014760 | 745764 | MBOVPG45_RS03225 | | | G>T |
| NC_014760 | 745780 | MBOVPG45_RS03225 | | | AT>GC |
| NC_014760 | 745790 | MBOVPG45_RS03225 | | | A>G |
| NC_014760 | 745794 | MBOVPG45_RS03225 | | | TA>AT |
| NC_014760 | 745826 | MBOVPG45_RS03225 | | | G>A |
| NC_014760 | 745868 | MBOVPG45_RS03225 | | | C>A |
| NC_014760 | 745877 | MBOVPG45_RS03225 | | | A>G |
| NC_014760 | 745902 | MBOVPG45_RS03225 | | | A>T |
| NC_014760 | 745930 | MBOVPG45_RS03225 | | | T>C |
| NC_014760 | 745997 | MBOVPG45_RS03225 | | | A>G |
| NC_014760 | 746021 | MBOVPG45_RS03225 | | | C>G |
| NC_014760 | 746047 | MBOVPG45_RS03225 | | | A>G |
| NC_014760 | 746049 | MBOVPG45_RS03225 | | | A>T |
| NC_014760 | 746080 | MBOVPG45_RS03225 | | | G>A |
| NC_014760 | 746086 | MBOVPG45_RS03225 | | | A>G |
| NC_014760 | 746150 | MBOVPG45_RS03225 | | | T>A |
| NC_014760 | 746206 | MBOVPG45_RS03225 | | | A>C |
| NC_014760 | 746213 | MBOVPG45_RS03225 | | | G>A |
| NC_014760 | 746224 | MBOVPG45_RS03225 | | | - > ins AATCGT |
| NC_014760 | 746227 | MBOVPG45_RS03225 | | | T>C |
| NC_014760 | 746243 | MBOVPG45_RS03225 | | | AG>GA |
| NC_014760 | 746246 | MBOVPG45_RS03225 | | | GA>AC |
| NC_014760 | 746251 | MBOVPG45_RS03225 | | | C>A |
| NC_014760 | 746253 | MBOVPG45_RS03225 | | | A>T |
| NC_014760 | 746327 | MBOVPG45_RS03225 | | | A>G |
| NC_014760 | 746331 | MBOVPG45_RS03225 | | | A>T |
| NC_014760 | 747240 | MBOVPG45_RS03230 | | | AC>GT |
| NC_014760 | 747247 | MBOVPG45_RS03230 | | | G>T |
| NC_014760 | 747252 | MBOVPG45_RS03230 | | | C>T |
| NC_014760 | 747263 | MBOVPG45_RS03230 | | | A>G |
| NC_014760 | 747273 | MBOVPG45_RS03230 | | | A>G |
| NC_014760 | 747325 | MBOVPG45_RS03230 | | | A>T |
| NC_014760 | 747385 | MBOVPG45_RS03230 | | | A>T |
| NC_014760 | 747420 | MBOVPG45_RS03230 | | | G>A |
| NC_014760 | 747431 | MBOVPG45_RS03230 | | | C>G |
| NC_014760 | 747444 | MBOVPG45_RS03230 | | | C>T |
| NC_014760 | 747456 | MBOVPG45_RS03230 | | | A>G |
| NC_014760 | 747471 | MBOVPG45_RS03230 | | | C>A |
| NC_014760 | 747480 | MBOVPG45_RS03230 | | | A>G |

| | | | | | |
|---|---|---|---|---|---|
| NC_014760 | 747504 | MBOVPG45_RS03230 | | | C>G |
| NC_014760 | 747633 | MBOVPG45_RS03230 | | | T>A |
| NC_014760 | 747689 | MBOVPG45_RS03230 | | | A>C |
| NC_014760 | 747696 | MBOVPG45_RS03230 | | | G>A |
| NC_014760 | 747707 | MBOVPG45_RS03230 | | | - > ins AATCGT |
| NC_014760 | 747710 | MBOVPG45_RS03230 | | | T>C |
| NC_014760 | 747730 | MBOVPG45_RS03230 | | | A>C |
| NC_014760 | 747741 | MBOVPG45_RS03230 | | | A>T |
| NC_014760 | 747810 | MBOVPG45_RS03230 | | | A>G |
| NC_014760 | 747814 | MBOVPG45_RS03230 | | | A>T |
| NC_014760 | 749051 | prs | | | T>G |
| NC_014760 | 749061 | rsmG (pseudo), prs | | | G>C |
| NC_014760 | 751100 | MBOVPG45_RS03260 | | | TG>CA |
| NC_014760 | 751138 | MBOVPG45_RS03260 | | | T>A |
| NC_014760 | 751487 | MBOVPG45_RS03260 | | | C>T |
| NC_014760 | 751600 | MBOVPG45_RS03260 | | | TT>GC |
| NC_014760 | 752783 | dcm | | | C>T |
| NC_014760 | 752808 | dcm | | | T>A |
| NC_014760 | 752824 | dcm | | | G>A |
| NC_014760 | 753684 | MBOVPG45_RS03275 | | | C>T |
| NC_014760 | 753796 | MBOVPG45_RS03275 | | | C>T |
| NC_014760 | 753856 | MBOVPG45_RS03275 | | C>T | |
| NC_014760 | 754072 | MBOVPG45_RS03275 | | | A>G |
| NC_014760 | 754116 | MBOVPG45_RS03275 | | | T>C |
| NC_014760 | 754547 | MBOVPG45_RS03280 | | | C>T |
| NC_014760 | 754796 | MBOVPG45_RS03280 | | | A>T |
| NC_014760 | 756247 | MBOVPG45_RS03285 | | | T>C |
| NC_014760 | 756346 | MBOVPG45_RS03285 | | | C>T |
| NC_014760 | 756353 | MBOVPG45_RS03285 | | | C>T |
| NC_014760 | 756812 | MBOVPG45_RS03285 | | | A>G |
| NC_014760 | 757435 | parC | MBOVPG45_RS0 | | G>A |
| NC_014760 | 757465 | parC | MBOVPG45_RS0 | | T>G |
| NC_014760 | 757589 | parC | MBOVPG45_RS0 | | A>C |
| NC_014760 | 757655 | parC | MBOVPG45_RS0 | | T>C |
| NC_014760 | 757712 | parC | MBOVPG45_RS0 | | G>A |
| NC_014760 | 757738 | parC | MBOVPG45_RS0 | | G>C |
| NC_014760 | 757769 | parC | MBOVPG45_RS0 | | T>C |
| NC_014760 | 757843 | parC | MBOVPG45_RS0 | | C>T |
| NC_014760 | 758012 | parC | MBOVPG45_RS0 | T>C | T>C |
| NC_014760 | 758282 | parC | MBOVPG45_RS0 | | A>T |
| NC_014760 | 758406 | parC | MBOVPG45_RS0 | | A>T |
| NC_014760 | 758594 | parC | MBOVPG45_RS0 | | GT>AC |
| NC_014760 | 758717 | parC | MBOVPG45_RS0 | | T>C |
| NC_014760 | 758798 | parC | MBOVPG45_RS0 | | T>C |
| NC_014760 | 758804 | parC | MBOVPG45_RS0 | | A>G |
| NC_014760 | 759116 | parC | MBOVPG45_RS0 | | C>T |
| NC_014760 | 759227 | parC | MBOVPG45_RS0 | | G>A |

| | | | | | |
|---|---|---|---|---|---|
| NC_014760 | 759293 | parC | MBOVPG45_RS0 | | T>G |
| NC_014760 | 759886 | parC | MBOVPG45_RS0 | | T>C |
| NC_014760 | 760101 | parE | | | TA>CG |
| NC_014760 | 760244 | parE | | | G>C |
| NC_014760 | 761359 | parE | | | T>C |
| NC_014760 | 761454 | parE | | | T>C |
| NC_014760 | 762108 | MBOVPG45_RS03300 | | | C>T |
| NC_014760 | 763025 | MBOVPG45_RS03300 | | | C>T |
| NC_014760 | 763284 | MBOVPG45_RS03300 | | | A>C |
| NC_014760 | 764252 | rsmD | | | G>A |
| NC_014760 | 764720 | MBOVPG45_RS03315 | | | T>C |
| NC_014760 | 764742 | MBOVPG45_RS03315 | | | T>G |
| NC_014760 | 764751 | MBOVPG45_RS03315 | | | A>G |
| NC_014760 | 764821 | MBOVPG45_RS03315 | | | C>T |
| NC_014760 | 764833 | MBOVPG45_RS03315 | | | T>C |
| NC_014760 | 764842 | MBOVPG45_RS03315 | | | C>T |
| NC_014760 | 764863 | MBOVPG45_RS03315 | | | T>C |
| NC_014760 | 765370 | MBOVPG45_RS03315 | | | T>C |
| NC_014760 | 765374 | MBOVPG45_RS03315 | | | C>T |
| NC_014760 | 765438 | MBOVPG45_RS03315 | | | A>G |
| NC_014760 | 765831 | MBOVPG45_RS03320 | | | T>A |
| NC_014760 | 766476 | MBOVPG45_RS03320 | | C>T | |
| NC_014760 | 766908 | MBOVPG45_RS03320 | | | A>G |
| NC_014760 | 766974 | MBOVPG45_RS03320 | | | A>G |
| NC_014760 | 767058 | MBOVPG45_RS03320 | | C>T | C>T |
| NC_014760 | 767941 | MBOVPG45_RS03325 | | | G>A |
| NC_014760 | 768041 | mnmG | | | A>C |
| NC_014760 | 768246 | mnmG | | | A>G |
| NC_014760 | 768374 | mnmG | | | A>G |
| NC_014760 | 768840 | mnmG | | | C>T |
| NC_014760 | 769402 | mnmG | | | A>C |
| NC_014760 | 769410 | mnmG | | | TG>CA |
| NC_014760 | 772736 | MBOVPG45_RS03345 | | | T>A |
| NC_014760 | 772745 | MBOVPG45_RS03345 | | | A>C |
| NC_014760 | 772837 | MBOVPG45_RS03345 | | | G>C |
| NC_014760 | 772864 | MBOVPG45_RS03345 | | | G>A |
| NC_014760 | 772902 | MBOVPG45_RS03345 | | | G>A |
| NC_014760 | 772980 | MBOVPG45_RS03345 | | | T>A |
| NC_014760 | 773025 | MBOVPG45_RS03345 | | | A>G |
| NC_014760 | 773157 | MBOVPG45_RS03345 | | | G>A |
| NC_014760 | 773193 | MBOVPG45_RS03345 | | | A>G |
| NC_014760 | 773244 | MBOVPG45_RS03345 | | | T>G |
| NC_014760 | 773281 | MBOVPG45_RS03345 | | | A>G |
| NC_014760 | 773386 | MBOVPG45_RS03345 | | - > ins A | |
| NC_014760 | 773818 | MBOVPG45_RS03345 | | | C>T |
| NC_014760 | 773820 | MBOVPG45_RS03345 | | | T>G |
| NC_014760 | 773826 | MBOVPG45_RS03345 | | | C>T |

| | | | | | |
|---|---|---|---|---|---|
| NC_014760 | 773847 | MBOVPG45_RS03345 | | | C>G |
| NC_014760 | 773853 | MBOVPG45_RS03345 | | | A>T |
| NC_014760 | 773986 | MBOVPG45_RS03345 | | | A>C |
| NC_014760 | 773989 | MBOVPG45_RS03345 | | | A>G |
| NC_014760 | 774226 | MBOVPG45_RS03345 | | | A>G |
| NC_014760 | 774230 | MBOVPG45_RS03345 | | | A>G |
| NC_014760 | 774291 | MBOVPG45_RS03345 | | | G>A |
| NC_014760 | 774357 | MBOVPG45_RS03345 | | | AT>GC |
| NC_014760 | 774539 | MBOVPG45_RS03345 | | | C>G |
| NC_014760 | 774540 | MBOVPG45_RS03345 | | | G>C |
| NC_014760 | 774671 | MBOVPG45_RS03350 | | | T>C |
| NC_014760 | 774743 | MBOVPG45_RS03350 | | | A>G |
| NC_014760 | 774977 | MBOVPG45_RS03350 | | | G>A |
| NC_014760 | 775004 | MBOVPG45_RS03350 | | | G>A |
| NC_014760 | 775089 | MBOVPG45_RS03350 | | | T>C |
| NC_014760 | 775129 | MBOVPG45_RS03350 | | | AC>TT |
| NC_014760 | 775262 | MBOVPG45_RS03350 | | | T>G |
| NC_014760 | 775399 | MBOVPG45_RS03350 | | | G>T |
| NC_014760 | 775641 | MBOVPG45_RS03350 | | | T>A |
| NC_014760 | 775897 | MBOVPG45_RS03350 | | | A>T |
| NC_014760 | 775902 | MBOVPG45_RS03350 | | | G>A |
| NC_014760 | 776724 | MBOVPG45_RS03360 | | | C>G |
| NC_014760 | 776846 | MBOVPG45_RS03360 | | | G>A |
| NC_014760 | 777040 | MBOVPG45_RS03360 | | | GC>TG |
| NC_014760 | 777239 | MBOVPG45_RS03360 | | | G>A |
| NC_014760 | 777261 | MBOVPG45_RS03360 | | | T>G |
| NC_014760 | 777314 | MBOVPG45_RS03360 | | | T>A |
| NC_014760 | 777455 | MBOVPG45_RS03360 | | | G>A |
| NC_014760 | 777584 | MBOVPG45_RS03360 | | | G>A |
| NC_014760 | 777660 | MBOVPG45_RS03360 | | | G>A |
| NC_014760 | 777706 | MBOVPG45_RS03360 | | | TG>CA |
| NC_014760 | 777821 | MBOVPG45_RS03360 | | | AA>GG |
| NC_014760 | 777842 | MBOVPG45_RS03360 | | | A>G |
| NC_014760 | 777882 | MBOVPG45_RS03360 | | | A>T |
| NC_014760 | 778219 | MBOVPG45_RS03365 | | | C>T |
| NC_014760 | 778351 | MBOVPG45_RS03365 | | | G>A |
| NC_014760 | 778461 | MBOVPG45_RS03365 | | | A>G |
| NC_014760 | 778638 | MBOVPG45_RS03365 | | | G>T |
| NC_014760 | 778656 | MBOVPG45_RS03365 | | | G>A |
| NC_014760 | 778669 | MBOVPG45_RS03365 | | | C>T |
| NC_014760 | 778912 | MBOVPG45_RS03365 | | | C>T |
| NC_014760 | 778917 | MBOVPG45_RS03365 | | | - > ins TTT |
| NC_014760 | 778944 | MBOVPG45_RS03365 | | | GT>AA |
| NC_014760 | 779074 | MBOVPG45_RS03365 | | | T>C |
| NC_014760 | 779101 | MBOVPG45_RS03365 | | | C>T |
| NC_014760 | 779213 | MBOVPG45_RS03365 | | | T>C |
| NC_014760 | 779393 | MBOVPG45_RS03365 | | | A>C |

| | | | | | |
|---|---|---|---|---|---|
| NC_014760 | 779403 | MBOVPG45_RS03365 | | | C>T |
| NC_014760 | 779442 | MBOVPG45_RS03365 | | | G>T |
| NC_014760 | 779448 | MBOVPG45_RS03365 | | | A>G |
| NC_014760 | 779454 | MBOVPG45_RS03365 | | | T>G |
| NC_014760 | 779470 | MBOVPG45_RS03365 | | | C>G |
| NC_014760 | 779477 | MBOVPG45_RS03365 | | | TA>TT |
| NC_014760 | 779542 | MBOVPG45_RS03365 | | | C>T |
| NC_014760 | 779620 | MBOVPG45_RS03365 | | | A>T |
| NC_014760 | 779784 | MBOVPG45_RS03365 | | | T>A |
| NC_014760 | 779806 | MBOVPG45_RS03365 | | | CA>TG |
| NC_014760 | 779868 | MBOVPG45_RS03365 | | | G>T |
| NC_014760 | 779880 | MBOVPG45_RS03365 | | | T>G |
| NC_014760 | 780828 | MBOVPG45_RS03375 | | | C>T |
| NC_014760 | 780846 | MBOVPG45_RS03375 | | | G>T |
| NC_014760 | 780957 | MBOVPG45_RS03375 | | | T > del 1 |
| NC_014760 | 780973 | MBOVPG45_RS03375 | | | T>A |
| NC_014760 | 781029 | MBOVPG45_RS03375 | | | G>A |
| NC_014760 | 781040 | MBOVPG45_RS03375 | | | AG>CT |
| NC_014760 | 781089 | MBOVPG45_RS03375 | | | A>C |
| NC_014760 | 783032 | MBOVPG45_RS03375 | | | CT>AC |
| NC_014760 | 783101 | MBOVPG45_RS03380,<br>MBOVPG45_RS03375 | | | A>G |
| NC_014760 | 783151 | MBOVPG45_RS03380 | | | G>A |
| NC_014760 | 783503 | MBOVPG45_RS03380 | | | A>G |
| NC_014760 | 783605 | MBOVPG45_RS03380 | | | C>A |
| NC_014760 | 783629 | MBOVPG45_RS03380 | | | A>G |
| NC_014760 | 783639 | MBOVPG45_RS03380 | | | G>A |
| NC_014760 | 783827 | MBOVPG45_RS03380 | | | G>A |
| NC_014760 | 783885 | MBOVPG45_RS03380 | | | C>G |
| NC_014760 | 784226 | MBOVPG45_RS03380 | | | G>A |
| NC_014760 | 784317 | MBOVPG45_RS03380 | | | G>A |
| NC_014760 | 784428 | MBOVPG45_RS03380 | | | A>C |
| NC_014760 | 784442 | MBOVPG45_RS03380 | | | C>T |
| NC_014760 | 784460 | MBOVPG45_RS03380 | | | C>A |
| NC_014760 | 785098 | dnaE | | | G>A |
| NC_014760 | 785455 | dnaE | | | A>G |
| NC_014760 | 785620 | dnaE | | | G>A |
| NC_014760 | 785746 | dnaE | | | A>C |
| NC_014760 | 785806 | dnaE | | | G>A |
| NC_014760 | 786067 | dnaE | | | A>G |
| NC_014760 | 786405 | dnaE | | G>T | |
| NC_014760 | 786551 | dnaE | | | A>G |
| NC_014760 | 786573 | dnaE | | | G>C |
| NC_014760 | 786720 | dnaE | | | A>T |
| NC_014760 | 786749 | dnaE | | | G>A |
| NC_014760 | 787120 | dnaE | | | T>G |
| NC_014760 | 787335 | dnaE | | | CG>TC |

| | | | | | |
|---|---|---|---|---|---|
| NC_014760 | 787452 | dnaE | | | AT>GG |
| NC_014760 | 787694 | dnaE | | | A>G |
| NC_014760 | 787750 | dnaE | | | G>T |
| NC_014760 | 788183 | MBOVPG45_RS03390 | | | A>G |
| NC_014760 | 788324 | MBOVPG45_RS03390 | | | A>G |
| NC_014760 | 788729 | MBOVPG45_RS03390 | | | G>A |
| NC_014760 | 788790 | MBOVPG45_RS03390 | | | AT>GC |
| NC_014760 | 789498 | MBOVPG45_RS03395 | | | T>C |
| NC_014760 | 790668 | MBOVPG45_RS03395 | | | G>A |
| NC_014760 | 791109 | MBOVPG45_RS03400 | | | C>A |
| NC_014760 | 791132 | MBOVPG45_RS03400 | | | C>G |
| NC_014760 | 791246 | MBOVPG45_RS03400 | | | C>T |
| NC_014760 | 791339 | MBOVPG45_RS03400 | | | C>T |
| NC_014760 | 791365 | MBOVPG45_RS03400 | | | C>T |
| NC_014760 | 791627 | MBOVPG45_RS03400 | | | G>T |
| NC_014760 | 791726 | MBOVPG45_RS03400 | | | A>G |
| NC_014760 | 791734 | MBOVPG45_RS03400 | | | T>C |
| NC_014760 | 792210 | MBOVPG45_RS03400 | | | T>A |
| NC_014760 | 792669 | MBOVPG45_RS03400 | | | A>C |
| NC_014760 | 792775 | MBOVPG45_RS03400 | | | A>T |
| NC_014760 | 796785 | MBOVPG45_RS03420 | | | A>G |
| NC_014760 | 796863 | MBOVPG45_RS03420 | | | G>A |
| NC_014760 | 796909 | MBOVPG45_RS03420 | | | G>A |
| NC_014760 | 796951 | MBOVPG45_RS03420 | | | G>T |
| NC_014760 | 797016 | MBOVPG45_RS03420 | | | T>C |
| NC_014760 | 797293 | MBOVPG45_RS03420 | | | C>A |
| NC_014760 | 798146 | MBOVPG45_RS03425 | | | A>G |
| NC_014760 | 802899 | MBOVPG45_RS03450 | | | A>G |
| NC_014760 | 803344 | MBOVPG45_RS03450 | | | A>G |
| NC_014760 | 803677 | MBOVPG45_RS03450 | | | G>A |
| NC_014760 | 805063 | MBOVPG45_RS03465 | | | C>T |
| NC_014760 | 805174 | MBOVPG45_RS03465 | | | C>T |
| NC_014760 | 805478 | MBOVPG45_RS03465 | | | T>A |
| NC_014760 | 805626 | MBOVPG45_RS03465 | | | A>G |
| NC_014760 | 806434 | MBOVPG45_RS03470 | | | T>C |
| NC_014760 | 806664 | MBOVPG45_RS03470 | | | A>T |
| NC_014760 | 806817 | MBOVPG45_RS03470 | | | G>T |
| NC_014760 | 806944 | MBOVPG45_RS03470 | | | G>A |
| NC_014760 | 807584 | MBOVPG45_RS03470 | | | C>T |
| NC_014760 | 807895 | MBOVPG45_RS03470 | | | AT>TG |
| NC_014760 | 808377 | MBOVPG45_RS03475 | | | AT>GC |
| NC_014760 | 808476 | MBOVPG45_RS03475 | | | C>A |
| NC_014760 | 808485 | MBOVPG45_RS03475 | | | T>C |
| NC_014760 | 808514 | MBOVPG45_RS03475 | | | T>A |
| NC_014760 | 808531 | MBOVPG45_RS03475 | | | G>A |
| NC_014760 | 808570 | MBOVPG45_RS03475 | | | TC>CT |
| NC_014760 | 808609 | MBOVPG45_RS03475 | | | T>C |

| | | | | | |
|---|---|---|---|---|---|
| NC_014760 | 808620 | MBOVPG45_RS03475 | | | C>T |
| NC_014760 | 808654 | MBOVPG45_RS03475 | | | C>T |
| NC_014760 | 808831 | MBOVPG45_RS03475 | | | T>C |
| NC_014760 | 808885 | MBOVPG45_RS03475 | | | T>A |
| NC_014760 | 808936 | MBOVPG45_RS03475 | | | C>T |
| NC_014760 | 809043 | MBOVPG45_RS03475 | | | T>C |
| NC_014760 | 809089 | MBOVPG45_RS03475 | | | C>T |
| NC_014760 | 809134 | MBOVPG45_RS03475 | | | T>C |
| NC_014760 | 809137 | MBOVPG45_RS03475 | | | A>C |
| NC_014760 | 809191 | MBOVPG45_RS03475 | | | T>C |
| NC_014760 | 809214 | MBOVPG45_RS03475 | | | C>T |
| NC_014760 | 809230 | MBOVPG45_RS03475 | | | T>C |
| NC_014760 | 809255 | MBOVPG45_RS03475 | | | A>T |
| NC_014760 | 809302 | MBOVPG45_RS03475 | | | A>T |
| NC_014760 | 809311 | MBOVPG45_RS03475 | | | T>C |
| NC_014760 | 809329 | MBOVPG45_RS03475 | | | A>C |
| NC_014760 | 809380 | MBOVPG45_RS03475 | | | A>G |
| NC_014760 | 809395 | MBOVPG45_RS03475 | | | T>C |
| NC_014760 | 809611 | MBOVPG45_RS03475 | | | T>C |
| NC_014760 | 810822 | MBOVPG45_RS03485 | | | G>A |
| NC_014760 | 810994 | MBOVPG45_RS03485 | | | A>G |
| NC_014760 | 811003 | MBOVPG45_RS03485 | | | TG>CA |
| NC_014760 | 811175 | MBOVPG45_RS03485 | | | TC>CT |
| NC_014760 | 811225 | MBOVPG45_RS03485 | | | C>T |
| NC_014760 | 811273 | MBOVPG45_RS03485 | | | T>C |
| NC_014760 | 811289 | MBOVPG45_RS03485 | | | T>C |
| NC_014760 | 811333 | MBOVPG45_RS03485 | | | G>A |
| NC_014760 | 811458 | MBOVPG45_RS03485 | | | C>T |
| NC_014760 | 811527 | MBOVPG45_RS03485 | | | A>G |
| NC_014760 | 812080 | MBOVPG45_RS03490 | | | G>A |
| NC_014760 | 814940 | MBOVPG45_RS03500 | | | T>G |
| NC_014760 | 815009 | MBOVPG45_RS03500 | | | CA>TC |
| NC_014760 | 817549 | MBOVPG45_RS03500 | | A>G | |
| NC_014760 | 817734 | MBOVPG45_RS03500 | | | C>A |
| NC_014760 | 817739 | MBOVPG45_RS03500 | | | T>C |
| NC_014760 | 817788 | MBOVPG45_RS03500 | | | T>A |
| NC_014760 | 817923 | MBOVPG45_RS03500 | | | A>T |
| NC_014760 | 818027 | MBOVPG45_RS03500 | | | C>T |
| NC_014760 | 818176 | MBOVPG45_RS03500 | | | C>T |
| NC_014760 | 818180 | MBOVPG45_RS03500 | | | T>C |
| NC_014760 | 818216 | MBOVPG45_RS03500 | | | T>C |
| NC_014760 | 818351 | MBOVPG45_RS03500 | | | C>T |
| NC_014760 | 818359 | MBOVPG45_RS03500 | | | T>C |
| NC_014760 | 818636 | MBOVPG45_RS03500 | | | T>C |
| NC_014760 | 818699 | MBOVPG45_RS03500 | | | T>C |
| NC_014760 | 818876 | MBOVPG45_RS03500 | | | TAT>CTC |
| NC_014760 | 818957 | MBOVPG45_RS03500 | | | A>C |

| | | | | | |
|---|---|---|---|---|---|
| NC_014760 | 819110 | MBOVPG45_RS03500 | | | T>C |
| NC_014760 | 819124 | MBOVPG45_RS03500 | | | G>A |
| NC_014760 | 819165 | MBOVPG45_RS03500 | | | T>A |
| NC_014760 | 819167 | MBOVPG45_RS03500 | | | G>A |
| NC_014760 | 819212 | MBOVPG45_RS03500 | | | T>C |
| NC_014760 | 819279 | MBOVPG45_RS03500 | | | G>T |
| NC_014760 | 819281 | MBOVPG45_RS03500 | | | T>C |
| NC_014760 | 819286 | MBOVPG45_RS03500 | | | T>C |
| NC_014760 | 819518 | MBOVPG45_RS03500 | | | T>C |
| NC_014760 | 819521 | MBOVPG45_RS03500 | | | A>G |
| NC_014760 | 819551 | MBOVPG45_RS03500 | | | T>G |
| NC_014760 | 819554 | MBOVPG45_RS03500 | | | C>T |
| NC_014760 | 819837 | MBOVPG45_RS03500 | | | A>T |
| NC_014760 | 819980 | MBOVPG45_RS03500 | | | TA>AG |
| NC_014760 | 820025 | MBOVPG45_RS03500 | | | T>G |
| NC_014760 | 820094 | MBOVPG45_RS03500 | | | A>T |
| NC_014760 | 820124 | MBOVPG45_RS03500 | | | C>T |
| NC_014760 | 820232 | MBOVPG45_RS03500 | | | C>T |
| NC_014760 | 820561 | MBOVPG45_RS03500 | | | A>G |
| NC_014760 | 820686 | MBOVPG45_RS03500 | | | C>A |
| NC_014760 | 820754 | MBOVPG45_RS03500 | | | T>C |
| NC_014760 | 820760 | MBOVPG45_RS03500 | | | T>G |
| NC_014760 | 820792 | MBOVPG45_RS03500 | | | T>A |
| NC_014760 | 820929 | MBOVPG45_RS03500 | | | A>C |
| NC_014760 | 820976 | MBOVPG45_RS03500 | | | T>C |
| NC_014760 | 821037 | MBOVPG45_RS03500 | | | T>G |
| NC_014760 | 821090 | MBOVPG45_RS03500 | | | C>T |
| NC_014760 | 821131 | MBOVPG45_RS03500 | | | A>G |
| NC_014760 | 821333 | MBOVPG45_RS03500 | | | C>T |
| NC_014760 | 821339 | MBOVPG45_RS03500 | | | T>G |
| NC_014760 | 821444 | MBOVPG45_RS03500 | | | G>A |
| NC_014760 | 821599 | MBOVPG45_RS03500 | | | G>A |
| NC_014760 | 821630 | MBOVPG45_RS03500 | | | CA>AT |
| NC_014760 | 821980 | MBOVPG45_RS03500 | | | C>T |
| NC_014760 | 822023 | MBOVPG45_RS03500 | | | A>C |
| NC_014760 | 822044 | MBOVPG45_RS03500 | | | T>C |
| NC_014760 | 822051 | MBOVPG45_RS03500 | | | A>T |
| NC_014760 | 822065 | MBOVPG45_RS03500 | | | G>C |
| NC_014760 | 822078 | MBOVPG45_RS03500 | | | A>T |
| NC_014760 | 822160 | MBOVPG45_RS03500 | | | T>C |
| NC_014760 | 822377 | MBOVPG45_RS03500 | | | T>A |
| NC_014760 | 826479 | rpoC | | | C>T |
| NC_014760 | 830647 | MBOVPG45_RS03510 | | | T>C |
| NC_014760 | 831335 | MBOVPG45_RS03515 | | | AA>GG |
| NC_014760 | 831344 | MBOVPG45_RS03515 | | | G>T |
| NC_014760 | 831422 | MBOVPG45_RS03515 | | | G>A |
| NC_014760 | 832675 | MBOVPG45_RS03525 | | | A>G |

| | | | | | |
|---|---|---|---|---|---|
| NC_014760 | 832862 | MBOVPG45_RS03525 | | | A>C |
| NC_014760 | 833816 | MBOVPG45_RS03530 | | | A>G |
| NC_014760 | 833936 | MBOVPG45_RS03530 | | | G>T |
| NC_014760 | 833992 | MBOVPG45_RS03530 | | | A>G |
| NC_014760 | 834059 | MBOVPG45_RS03530 | | | A>G |
| NC_014760 | 834110 | MBOVPG45_RS03530 | | | A>G |
| NC_014760 | 834722 | lysS | | | A>G |
| NC_014760 | 835100 | lysS | | | T>C |
| NC_014760 | 835161 | lysS | | | G>A |
| NC_014760 | 835897 | MBOVPG45_RS03540 | | | C>A |
| NC_014760 | 836023 | MBOVPG45_RS03540 | | | A>G |
| NC_014760 | 836284 | MBOVPG45_RS03540 | | | GCC>AGT |
| NC_014760 | 836508 | MBOVPG45_RS03540 | | | CG>TA |
| NC_014760 | 836599 | MBOVPG45_RS03540 | | | A>T |
| NC_014760 | 836611 | MBOVPG45_RS03540 | | | G>C |
| NC_014760 | 836887 | MBOVPG45_RS03545 | | | T>C |
| NC_014760 | 836970 | MBOVPG45_RS03545 | | | G>A |
| NC_014760 | 837576 | MBOVPG45_RS03545 | | | G>T |
| NC_014760 | 837813 | MBOVPG45_RS03545 | | | T>C |
| NC_014760 | 837877 | MBOVPG45_RS03545 | | | C>T |
| NC_014760 | 837916 | MBOVPG45_RS03545 | | | T>C |
| NC_014760 | 838455 | MBOVPG45_RS03545 | | | G>A |
| NC_014760 | 838869 | MBOVPG45_RS03545 | | | T>C |
| NC_014760 | 839138 | MBOVPG45_RS03550 | | | G>A |
| NC_014760 | 840182 | MBOVPG45_RS03550 | | | A>G |
| NC_014760 | 840434 | MBOVPG45_RS03550 | | | T>C |
| NC_014760 | 840918 | MBOVPG45_RS03550 | | | T>A |
| NC_014760 | 840993 | MBOVPG45_RS03550 | | | CA>AC |
| NC_014760 | 843830 | MBOVPG45_RS03560 | | G>A | |
| NC_014760 | 846198 | rpsP | | | AT>GC |
| NC_014760 | 846201 | rpsP | | | G>A |
| NC_014760 | 847407 | fmt | | | TTAGTATTC > del 9 |
| NC_014760 | 847477 | fmt | | | TG>AA |
| NC_014760 | 847760 | fmt | | | A>G |
| NC_014760 | 847842 | fmt | | | T>C |
| NC_014760 | 848393 | MBOVPG45_RS03590 | | | C>A |
| NC_014760 | 848426 | MBOVPG45_RS03590 | | | A>G |
| NC_014760 | 848551 | MBOVPG45_RS03590 | | | T>G |
| NC_014760 | 848639 | MBOVPG45_RS03590 | | | A>G |
| NC_014760 | 848668 | MBOVPG45_RS03590 | | | G>T |
| NC_014760 | 848747 | MBOVPG45_RS03590 | | | A>T |
| NC_014760 | 848835 | MBOVPG45_RS03590 | | | G>A |
| NC_014760 | 849064 | MBOVPG45_RS03590 | | | G>A |
| NC_014760 | 849300 | MBOVPG45_RS03590 | | | AG>GA |
| NC_014760 | 849304 | MBOVPG45_RS03590 | | | TTG>ACC |
| NC_014760 | 849308 | MBOVPG45_RS03590 | | | C>T |

| | | | | | |
|---|---|---|---|---|---|
| NC_014760 | 849495 | MBOVPG45_RS03590 | | | T>A |
| NC_014760 | 849569 | MBOVPG45_RS03590 | | | A>G |
| NC_014760 | 849618 | MBOVPG45_RS03590 | | | T>G |
| NC_014760 | 849691 | MBOVPG45_RS03590 | | | G>T |
| NC_014760 | 849858 | MBOVPG45_RS03590 | | | CA>TG |
| NC_014760 | 850069 | MBOVPG45_RS03590 | | | G>A |
| NC_014760 | 850162 | MBOVPG45_RS03590 | | | G>T |
| NC_014760 | 850180 | MBOVPG45_RS03590 | | | A>G |
| NC_014760 | 850273 | MBOVPG45_RS03590 | | | A>G |
| NC_014760 | 850282 | MBOVPG45_RS03590 | | | A>C |
| NC_014760 | 850284 | MBOVPG45_RS03590 | | | C>A |
| NC_014760 | 850840 | MBOVPG45_RS03590 | | | AA>GG |
| NC_014760 | 851190 | MBOVPG45_RS03590 | | | G>C |
| NC_014760 | 851192 | MBOVPG45_RS03590 | | | A>T |
| NC_014760 | 851233 | MBOVPG45_RS03590 | | | T>A |
| NC_014760 | 851236 | MBOVPG45_RS03590 | | | A>G |
| NC_014760 | 851272 | MBOVPG45_RS03590 | | | A>G |
| NC_014760 | 851558 | ptsP | | | T>C |
| NC_014760 | 851830 | ptsP | | | T>C |
| NC_014760 | 852125 | ptsP | | | G>C |
| NC_014760 | 852256 | ptsP | | | A>G |
| NC_014760 | 852347 | ptsP | | | T>C |
| NC_014760 | 852655 | ptsP | | | C>T |
| NC_014760 | 852983 | ptsP | | | A>G |
| NC_014760 | 853121 | MBOVPG45_RS03600 | | A > del 1 | |
| NC_014760 | 853237 | MBOVPG45_RS03600 | | | AG>GA |
| NC_014760 | 853967 | MBOVPG45_RS03600 | | | G>A |
| NC_014760 | 854021 | MBOVPG45_RS03605 | | | CA>TG |
| NC_014760 | 854164 | MBOVPG45_RS03605 | | | T>C |
| NC_014760 | 854276 | MBOVPG45_RS03605 | | | T>C |
| NC_014760 | 854467 | MBOVPG45_RS03605 | | | C>T |
| NC_014760 | 854531 | araD | | | A>T |
| NC_014760 | 855274 | MBOVPG45_RS03615 | | | A>C |
| NC_014760 | 855802 | MBOVPG45_RS03615 | | | A>C |
| NC_014760 | 855890 | MBOVPG45_RS03615 | | | C>T |
| NC_014760 | 855920 | MBOVPG45_RS03615 | | | T>C |
| NC_014760 | 856260 | MBOVPG45_RS03620 | | | T>C |
| NC_014760 | 856366 | MBOVPG45_RS03620 | | | TG>CA |
| NC_014760 | 856575 | MBOVPG45_RS03620 | | | G>T |
| NC_014760 | 856957 | MBOVPG45_RS03625 | | | G>A |
| NC_014760 | 858957 | MBOVPG45_RS03635 | | | A>C |
| NC_014760 | 860078 | MBOVPG45_RS03640 | | | C>A |
| NC_014760 | 860963 | MBOVPG45_RS03645 | | | A>G |
| NC_014760 | 861180 | MBOVPG45_RS03645 | | | A>G |
| NC_014760 | 861225 | MBOVPG45_RS03645 | | | A>G |
| NC_014760 | 861316 | MBOVPG45_RS03645 | | | A>G |
| NC_014760 | 861423 | MBOVPG45_RS03645 | | | C>G |

| | | | | | |
|---|---|---|---|---|---|
| NC_014760 | 861444 | MBOVPG45_RS03645 | | | T>A |
| NC_014760 | 861982 | MBOVPG45_RS03650 | | | T > del 1 |
| NC_014760 | 862036 | MBOVPG45_RS03650 | | | A>G |
| NC_014760 | 862054 | MBOVPG45_RS03650 | | | T>C |
| NC_014760 | 862065 | MBOVPG45_RS03650 | | | A>T |
| NC_014760 | 862076 | MBOVPG45_RS03650 | | | C>T |
| NC_014760 | 862079 | MBOVPG45_RS03650 | | | C>A |
| NC_014760 | 862082 | MBOVPG45_RS03650 | | | T>A |
| NC_014760 | 862088 | MBOVPG45_RS03650 | | | G>A |
| NC_014760 | 862107 | MBOVPG45_RS03650 | | | T>G |
| NC_014760 | 862525 | MBOVPG45_RS03650 | | | G>T |
| NC_014760 | 862609 | MBOVPG45_RS03650 | | | A>T |
| NC_014760 | 862657 | MBOVPG45_RS03650 | | | C>G |
| NC_014760 | 862854 | MBOVPG45_RS04565 | | | AG > del 2 |
| NC_014760 | 862890 | MBOVPG45_RS04565 | | | C>T |
| NC_014760 | 862893 | MBOVPG45_RS04565 | | | A>C |
| NC_014760 | 862923 | MBOVPG45_RS04565 | | | T>C |
| NC_014760 | 862928 | MBOVPG45_RS04565 | | | C>T |
| NC_014760 | 863158 | MBOVPG45_RS03655 | | | T>C |
| NC_014760 | 863249 | MBOVPG45_RS03655 | | | A>C |
| NC_014760 | 863267 | MBOVPG45_RS03655 | | | TT>TA |
| NC_014760 | 863293 | MBOVPG45_RS03655 | | | T>G |
| NC_014760 | 863311 | MBOVPG45_RS03655 | | | T>A |
| NC_014760 | 863341 | MBOVPG45_RS03655 | | | T>C |
| NC_014760 | 863345 | MBOVPG45_RS03655 | | | T>A |
| NC_014760 | 863386 | MBOVPG45_RS03655 | | | A>T |
| NC_014760 | 863406 | MBOVPG45_RS03655 | | | G>T |
| NC_014760 | 863408 | MBOVPG45_RS03655 | | | C>G |
| NC_014760 | 863416 | MBOVPG45_RS03655 | | | T>G |
| NC_014760 | 863421 | MBOVPG45_RS03655 | | | A>T |
| NC_014760 | 863437 | MBOVPG45_RS03655 | | | A>G |
| NC_014760 | 863462 | MBOVPG45_RS03655 | | | T>C |
| NC_014760 | 863487 | MBOVPG45_RS03655 | | | T>A |
| NC_014760 | 863535 | MBOVPG45_RS03655 | | | AT>TG |
| NC_014760 | 863546 | MBOVPG45_RS03655 | | | T>C |
| NC_014760 | 863607 | MBOVPG45_RS03655 | | | T>A |
| NC_014760 | 863612 | MBOVPG45_RS03655 | | | T>A |
| NC_014760 | 863620 | MBOVPG45_RS03655 | | | C>T |
| NC_014760 | 863684 | MBOVPG45_RS03655 | | | C>T |
| NC_014760 | 863689 | MBOVPG45_RS03655 | | | TC>GT |
| NC_014760 | 863741 | MBOVPG45_RS03655 | | | G>C |
| NC_014760 | 863764 | MBOVPG45_RS03655 | | | A>C |
| NC_014760 | 863774 | MBOVPG45_RS03655 | | | C>T |
| NC_014760 | 863839 | MBOVPG45_RS03655 | | | A>G |
| NC_014760 | 863869 | MBOVPG45_RS03655 | | | A>T |
| NC_014760 | 863949 | MBOVPG45_RS03655 | | | G>T |
| NC_014760 | 863960 | MBOVPG45_RS03655 | | | T>C |

| | | | | | |
|---|---|---|---|---|---|
| NC_014760 | 864006 | MBOVPG45_RS03655 | | | C>A |
| NC_014760 | 864018 | MBOVPG45_RS03655 | | | A>T |
| NC_014760 | 864022 | MBOVPG45_RS03655 | | | G>T |
| NC_014760 | 864032 | MBOVPG45_RS03655 | | | A>C |
| NC_014760 | 864038 | MBOVPG45_RS03655 | | | C>T |
| NC_014760 | 864116 | MBOVPG45_RS03655 | | | TA>CC |
| NC_014760 | 864315 | MBOVPG45_RS03660 | | - > ins G | |
| NC_014760 | 864316 | MBOVPG45_RS03660 | | | G > del 1 |
| NC_014760 | 864457 | MBOVPG45_RS03660 | | | A>G |
| NC_014760 | 864948 | MBOVPG45_RS03665 | | | A>G |
| NC_014760 | 864991 | MBOVPG45_RS03665 | | | T>A |
| NC_014760 | 865375 | MBOVPG45_RS03665 | | | GG>AA |
| NC_014760 | 865436 | MBOVPG45_RS03665 | | | A>G |
| NC_014760 | 865652 | MBOVPG45_RS03665 | | | A>G |
| NC_014760 | 865771 | MBOVPG45_RS03665 | | | AGG>GAA |
| NC_014760 | 865893 | MBOVPG45_RS04450 | | | C>T |
| NC_014760 | 866203 | MBOVPG45_RS04450 | | - > ins G | |
| NC_014760 | 866204 | MBOVPG45_RS04450 | | | GG > del 2 |
| NC_014760 | 866317 | MBOVPG45_RS04570 | | | T>A |
| NC_014760 | 867221 | MBOVPG45_RS04575 | | | C>T |
| NC_014760 | 867320 | MBOVPG45_RS04575 | | | A>G |
| NC_014760 | 867375 | MBOVPG45_RS04575 | | | T>C |
| NC_014760 | 867385 | MBOVPG45_RS04575 | | | G>A |
| NC_014760 | 867510 | MBOVPG45_RS04575 | | | GG>AA |
| NC_014760 | 870763 | MBOVPG45_RS03695 | | | C>T |
| NC_014760 | 871183 | MBOVPG45_RS03700 | | | T>C |
| NC_014760 | 871186 | MBOVPG45_RS03700 | | | C>T |
| NC_014760 | 871250 | MBOVPG45_RS03700 | | | T>A |
| NC_014760 | 871408 | MBOVPG45_RS03700 | | | C>T |
| NC_014760 | 871452 | MBOVPG45_RS03700 | | | T>A |
| NC_014760 | 871456 | MBOVPG45_RS03700 | | | T>G |
| NC_014760 | 871561 | MBOVPG45_RS03700 | | | CT>TG |
| NC_014760 | 871591 | MBOVPG45_RS03700 | | | T>C |
| NC_014760 | 871667 | MBOVPG45_RS03700 | | | T>C |
| NC_014760 | 872162 | MBOVPG45_RS03700 | | | G>T |
| NC_014760 | 872171 | MBOVPG45_RS03700 | | | T>C |
| NC_014760 | 872180 | MBOVPG45_RS03700 | | | GT>CA |
| NC_014760 | 872231 | MBOVPG45_RS03700 | | | T>G |
| NC_014760 | 872236 | MBOVPG45_RS03700 | | | T>C |
| NC_014760 | 872259 | MBOVPG45_RS03700 | | | T>G |
| NC_014760 | 872263 | MBOVPG45_RS03700 | | | TT>CC |
| NC_014760 | 872266 | MBOVPG45_RS03700 | | | G>T |
| NC_014760 | 872276 | MBOVPG45_RS03700 | | | C>T |
| NC_014760 | 872352 | MBOVPG45_RS03700 | | | T>G |
| NC_014760 | 872358 | MBOVPG45_RS03700 | | | A>C |
| NC_014760 | 872363 | MBOVPG45_RS03700 | | | C>T |
| NC_014760 | 872390 | MBOVPG45_RS03700 | | | T>C |

| | | | | | |
|---|---|---|---|---|---|
| NC_014760 | 872485 | MBOVPG45_RS03700 | | | T>C |
| NC_014760 | 872540 | MBOVPG45_RS03700 | | | TA>CT |
| NC_014760 | 872587 | MBOVPG45_RS03700 | | | G>T |
| NC_014760 | 872633 | MBOVPG45_RS03700 | | | C>T |
| NC_014760 | 872819 | MBOVPG45_RS03700 | | | C>T |
| NC_014760 | 872826 | MBOVPG45_RS03700 | | | T>A |
| NC_014760 | 872930 | MBOVPG45_RS03700 | | | T>C |
| NC_014760 | 872966 | MBOVPG45_RS03700 | | | A>T |
| NC_014760 | 873133 | MBOVPG45_RS03705 | | | A>T |
| NC_014760 | 873543 | MBOVPG45_RS03705 | | | T>C |
| NC_014760 | 873556 | MBOVPG45_RS03705 | | | TC>GT |
| NC_014760 | 873623 | MBOVPG45_RS03705 | | | C>T |
| NC_014760 | 873663 | MBOVPG45_RS03705 | | | G>A |
| NC_014760 | 873827 | MBOVPG45_RS03705 | | | T>A |
| NC_014760 | 873886 | MBOVPG45_RS03710 | | | C>T |
| NC_014760 | 876432 | MBOVPG45_RS03720 | | | C>T |
| NC_014760 | 876612 | MBOVPG45_RS03720 | | | T>C |
| NC_014760 | 876666 | MBOVPG45_RS03720 | | | T>G |
| NC_014760 | 876790 | MBOVPG45_RS03720 | | | G>T |
| NC_014760 | 876811 | MBOVPG45_RS03720 | | | T>G |
| NC_014760 | 877216 | MBOVPG45_RS03720 | | | C>T |
| NC_014760 | 877222 | MBOVPG45_RS03720 | | | T>A |
| NC_014760 | 877226 | MBOVPG45_RS03720 | | | A>T |
| NC_014760 | 877515 | MBOVPG45_RS03720 | | | G>A |
| NC_014760 | 877522 | MBOVPG45_RS03720 | | | A>T |
| NC_014760 | 877567 | MBOVPG45_RS03720 | | | C>T |
| NC_014760 | 877843 | MBOVPG45_RS03720 | | | TC>CT |
| NC_014760 | 877851 | MBOVPG45_RS03720 | | | TT>CC |
| NC_014760 | 877855 | MBOVPG45_RS03720 | | | T>C |
| NC_014760 | 878034 | MBOVPG45_RS03720 | | | T>C |
| NC_014760 | 878121 | MBOVPG45_RS03720 | | | T>C |
| NC_014760 | 878193 | MBOVPG45_RS03720 | | | C>T |
| NC_014760 | 878208 | MBOVPG45_RS03720 | | | TT>CG |
| NC_014760 | 878383 | MBOVPG45_RS03720 | | | A>C |
| NC_014760 | 878624 | MBOVPG45_RS03720 | | | AC>TT |
| NC_014760 | 878796 | MBOVPG45_RS03720 | | | A>T |
| NC_014760 | 878853 | MBOVPG45_RS03720 | | | C>T |
| NC_014760 | 878886 | MBOVPG45_RS03720 | | | T>C |
| NC_014760 | 878897 | MBOVPG45_RS03720 | | | T>A |
| NC_014760 | 878937 | MBOVPG45_RS03720 | | | AT>GC |
| NC_014760 | 879655 | MBOVPG45_RS03725 | | | A>T |
| NC_014760 | 879864 | MBOVPG45_RS03725 | | | T>C |
| NC_014760 | 880184 | MBOVPG45_RS03730 | | | C>T |
| NC_014760 | 880421 | MBOVPG45_RS03730 | | | T>C |
| NC_014760 | 880921 | nadE | | | C>T |
| NC_014760 | 880963 | nadE | | | C>T |
| NC_014760 | 881185 | nadE | | | A>G |

| | | | | | |
|---|---|---|---|---|---|
| NC_014760 | 881806 | MBOVPG45_RS03740 | | | C>T |
| NC_014760 | 882225 | MBOVPG45_RS03740 | | | A>T |
| NC_014760 | 882289 | MBOVPG45_RS03740 | | | T>C |
| NC_014760 | 882310 | MBOVPG45_RS03740 | | | AA>GG |
| NC_014760 | 882360 | MBOVPG45_RS03740 | | | C>T |
| NC_014760 | 882769 | MBOVPG45_RS03745 | | | T>C |
| NC_014760 | 882791 | MBOVPG45_RS03745 | | | T>C |
| NC_014760 | 883497 | MBOVPG45_RS03745 | | | C>T |
| NC_014760 | 883564 | MBOVPG45_RS03745 | | | T>G |
| NC_014760 | 883839 | rlmD | | | CT>TC |
| NC_014760 | 884159 | rlmD | | | T>A |
| NC_014760 | 884190 | rlmD | | | T>C |
| NC_014760 | 884219 | rlmD | | | C>G |
| NC_014760 | 884323 | rlmD | | | A>T |
| NC_014760 | 884353 | rlmD | | | A>T |
| NC_014760 | 884410 | rlmD | | | T>C |
| NC_014760 | 884449 | rlmD | | | C>T |
| NC_014760 | 884494 | rlmD | | | C>T |
| NC_014760 | 884623 | rlmD | | | C>T |
| NC_014760 | 884821 | rlmD | | | A>C |
| NC_014760 | 885156 | MBOVPG45_RS03755 | | G>A | |
| NC_014760 | 886043 | MBOVPG45_RS03760 | | | T>C |
| NC_014760 | 886105 | MBOVPG45_RS03760 | | | C>T |
| NC_014760 | 886205 | MBOVPG45_RS03760 | | | C>T |
| NC_014760 | 886419 | MBOVPG45_RS03760 | | | A>T |
| NC_014760 | 887092 | MBOVPG45_RS03765 | | | C>T |
| NC_014760 | 887209 | MBOVPG45_RS03765 | | T>C | |
| NC_014760 | 887211 | MBOVPG45_RS03765 | | | A>G |
| NC_014760 | 887501 | MBOVPG45_RS03765 | | | A>T |
| NC_014760 | 887599 | MBOVPG45_RS03770 | | | G>A |
| NC_014760 | 887784 | MBOVPG45_RS03770 | | | GC>AT |
| NC_014760 | 887878 | MBOVPG45_RS03770 | | | TT>AG |
| NC_014760 | 887881 | MBOVPG45_RS03770 | | | G>A |
| NC_014760 | 887932 | MBOVPG45_RS03770 | | | T>C |
| NC_014760 | 890617 | MBOVPG45_RS03795 | | | C>G |
| NC_014760 | 890631 | MBOVPG45_RS03795 | | | G>A |
| NC_014760 | 890787 | MBOVPG45_RS03795 | | | A>C |
| NC_014760 | 891013 | MBOVPG45_RS03795 | | | A>G |
| NC_014760 | 891116 | MBOVPG45_RS03795 | | | G>A |
| NC_014760 | 891165 | MBOVPG45_RS03795 | | | A>G |
| NC_014760 | 891348 | MBOVPG45_RS03795 | | | A>G |
| NC_014760 | 891370 | MBOVPG45_RS03795 | | | A>T |
| NC_014760 | 891446 | MBOVPG45_RS03795 | | | AA>TG |
| NC_014760 | 891450 | MBOVPG45_RS03795 | | | A>G |
| NC_014760 | 891550 | MBOVPG45_RS03795 | | | C>A |
| NC_014760 | 891887 | MBOVPG45_RS03800 | | | C>T |
| NC_014760 | 891919 | MBOVPG45_RS03800 | | | G>A |

| | | | | | |
|---|---|---|---|---|---|
| NC_014760 | 891926 | MBOVPG45_RS03800 | | | C>T |
| NC_014760 | 892442 | MBOVPG45_RS03800 | | | C>T |
| NC_014760 | 892502 | MBOVPG45_RS03800 | | | A>G |
| NC_014760 | 892553 | MBOVPG45_RS03800 | | | C>T |
| NC_014760 | 892589 | MBOVPG45_RS03800 | | | A>T |
| NC_014760 | 892961 | MBOVPG45_RS03800 | | | T>A |
| NC_014760 | 896766 | MBOVPG45_RS03820 | | | G>T |
| NC_014760 | 898026 | MBOVPG45_RS03825 | | T>C | T>C |
| NC_014760 | 899288 | MBOVPG45_RS03835 | | | G>T |
| NC_014760 | 900146 | trmB | | | G>A |
| NC_014760 | 900187 | trmB | | | C>T |
| NC_014760 | 900244 | trmB | | | C>A |
| NC_014760 | 900450 | trmB | | | T>C |
| NC_014760 | 901081 | MBOVPG45_RS03850 | | | T>A |
| NC_014760 | 901211 | MBOVPG45_RS03850 | | | G>A |
| NC_014760 | 904514 | rsmI | | | T>A |
| NC_014760 | 904552 | rsmI | | | G>A |
| NC_014760 | 904581 | rsmI | | | A>C |
| NC_014760 | 904584 | rsmI | | | A>T |
| NC_014760 | 904657 | rsmI | | | T>C |
| NC_014760 | 905196 | MBOVPG45_RS03870 | | | T>C |
| NC_014760 | 905402 | MBOVPG45_RS03870 | | | G>A |
| NC_014760 | 905514 | MBOVPG45_RS03870 | | | A>G |
| NC_014760 | 905540 | MBOVPG45_RS03870 | | | A>G |
| NC_014760 | 905559 | MBOVPG45_RS03870 | | | A>G |
| NC_014760 | 905604 | MBOVPG45_RS03870 | | | A>T |
| NC_014760 | 906098 | MBOVPG45_RS03875 | | | C>T |
| NC_014760 | 906191 | MBOVPG45_RS03875 | | G>T | |
| NC_014760 | 906637 | MBOVPG45_RS03875 | | | T>C |
| NC_014760 | 906851 | recR | | | T>A |
| NC_014760 | 907206 | recR | | | C>T |
| NC_014760 | 907225 | recR | | | A>G |
| NC_014760 | 907714 | dnaX | | | A>G |
| NC_014760 | 907986 | dnaX | | | C>T |
| NC_014760 | 908027 | dnaX | | | T>G |
| NC_014760 | 908113 | dnaX | | | C>T |
| NC_014760 | 908157 | dnaX | | | G>T |
| NC_014760 | 908775 | dnaX | | | G>A |
| NC_014760 | 911465 | MBOVPG45_RS03900 | | | A>G |
| NC_014760 | 911480 | MBOVPG45_RS03900 | | | C>T |
| NC_014760 | 911611 | MBOVPG45_RS03900 | | | A>G |
| NC_014760 | 911794 | MBOVPG45_RS03900 | | | C>A |
| NC_014760 | 912118 | MBOVPG45_RS03905 | | | A>G |
| NC_014760 | 912479 | MBOVPG45_RS03905 | | | G>A |
| NC_014760 | 912489 | MBOVPG45_RS03905 | | | A>T |
| NC_014760 | 912665 | MBOVPG45_RS03905 | | A>T | |
| NC_014760 | 912735 | MBOVPG45_RS03905 | | | A>T |

| | | | | | |
|---|---|---|---|---|---|
| NC_014760 | 912940 | MBOVPG45_RS03905 | | | A>C |
| NC_014760 | 913220 | MBOVPG45_RS03905 | | | T>C |
| NC_014760 | 913396 | MBOVPG45_RS03905 | | | G>A |
| NC_014760 | 913466 | MBOVPG45_RS03905 | | | G>A |
| NC_014760 | 913537 | MBOVPG45_RS03905 | | | G>A |
| NC_014760 | 913679 | MBOVPG45_RS03910 | | | A>C |
| NC_014760 | 913685 | MBOVPG45_RS03910 | | | G>A |
| NC_014760 | 913713 | MBOVPG45_RS03910 | | | C>T |
| NC_014760 | 913788 | MBOVPG45_RS03910 | | | A>C |
| NC_014760 | 914071 | MBOVPG45_RS03910 | | | A>G |
| NC_014760 | 914079 | MBOVPG45_RS03910 | | | A>G |
| NC_014760 | 914085 | MBOVPG45_RS03910 | | | T>G |
| NC_014760 | 914097 | MBOVPG45_RS03910 | | | A>G |
| NC_014760 | 914178 | MBOVPG45_RS03910 | | | G>A |
| NC_014760 | 914296 | MBOVPG45_RS03910 | | | A>G |
| NC_014760 | 914452 | MBOVPG45_RS03910 | | | A>C |
| NC_014760 | 914458 | MBOVPG45_RS03910 | | | T>A |
| NC_014760 | 914797 | MBOVPG45_RS03915 | | | C>T |
| NC_014760 | 914800 | MBOVPG45_RS03915 | | | T>C |
| NC_014760 | 914808 | MBOVPG45_RS03915 | | | C>G |
| NC_014760 | 914848 | MBOVPG45_RS03915 | | | A>G |
| NC_014760 | 915127 | MBOVPG45_RS03915 | | | G>A |
| NC_014760 | 915185 | MBOVPG45_RS03915 | | | GG>AA |
| NC_014760 | 915209 | MBOVPG45_RS03915 | | | A>C |
| NC_014760 | 915224 | MBOVPG45_RS03915 | | | A>T |
| NC_014760 | 915257 | MBOVPG45_RS03915 | | | T>A |
| NC_014760 | 915336 | MBOVPG45_RS03915 | | | G>A |
| NC_014760 | 915811 | MBOVPG45_RS03925 | | | GG>AA |
| NC_014760 | 915952 | MBOVPG45_RS03925 | | | C>A |
| NC_014760 | 916008 | MBOVPG45_RS03925 | | | T>A |
| NC_014760 | 916526 | pgsA | | | G>A |
| NC_014760 | 916566 | pgsA | | | A>G |
| NC_014760 | 917577 | MBOVPG45_RS03935 | | | T>C |
| NC_014760 | 917606 | MBOVPG45_RS03935 | | | C>T |
| NC_014760 | 918202 | MBOVPG45_RS03940 | | | A>G |
| NC_014760 | 918256 | MBOVPG45_RS03940 | | | A>G |
| NC_014760 | 918372 | MBOVPG45_RS03940 | | - > ins TTGT | - > ins TTGT |
| NC_014760 | 918571 | MBOVPG45_RS03945 | | | C>A |
| NC_014760 | 920868 | MBOVPG45_RS03955, MBOVPG45_RS03950 | | | C>T |
| NC_014760 | 920884 | MBOVPG45_RS03955 | | | C>T |
| NC_014760 | 921055 | MBOVPG45_RS03955 | | | C>T |
| NC_014760 | 921262 | nusA | | | A>G |
| NC_014760 | 921317 | nusA | | | C>T |
| NC_014760 | 921717 | nusA | | | CG>TA |
| NC_014760 | 921776 | nusA | | | G>C |
| NC_014760 | 921827 | nusA | | | T>C |

| | | | | | |
|---|---|---|---|---|---|
| NC_014760 | 922250 | nusA | | | T>C |
| NC_014760 | 928069 | MBOVPG45_RS03990 | | | C>T |
| NC_014760 | 929129 | MBOVPG45_RS03995 | | | A>T |
| NC_014760 | 929308 | MBOVPG45_RS03995 | | | T>C |
| NC_014760 | 930394 | MBOVPG45_RS04005 | | AG>GC | |
| NC_014760 | 930654 | MBOVPG45_RS04005 | | | T>C |
| NC_014760 | 931489 | MBOVPG45_RS04010 | | | G>C |
| NC_014760 | 931495 | MBOVPG45_RS04010 | | | C>T |
| NC_014760 | 931499 | MBOVPG45_RS04010 | | | A>T |
| NC_014760 | 931501 | MBOVPG45_RS04010 | | | C>T |
| NC_014760 | 931505 | MBOVPG45_RS04010 | | | GTA>TCT |
| NC_014760 | 931509 | MBOVPG45_RS04010 | | | T>A |
| NC_014760 | 931513 | MBOVPG45_RS04010 | | | CTA > del 3 |
| NC_014760 | 931517 | MBOVPG45_RS04010 | | | T>A |
| NC_014760 | 931519 | MBOVPG45_RS04010 | | | TT>CA |
| NC_014760 | 931673 | MBOVPG45_RS04010 | | A>T | |
| NC_014760 | 931677 | MBOVPG45_RS04010 | | TTTTTGCT > del 8 | |
| NC_014760 | 931693 | MBOVPG45_RS04010 | | TG>CT | |
| NC_014760 | 931700 | MBOVPG45_RS04010 | | TC>AT | |
| NC_014760 | 931952 | MBOVPG45_RS04010 | | | G>T |
| NC_014760 | 931995 | MBOVPG45_RS04010 | | | T>C |
| NC_014760 | 932093 | MBOVPG45_RS04010 | | | C>A |
| NC_014760 | 932142 | MBOVPG45_RS04010 | | | A>G |
| NC_014760 | 934781 | MBOVPG45_RS04465 | | A>G | |
| NC_014760 | 934783 | MBOVPG45_RS04465 | | A>C | |
| NC_014760 | 934794 | MBOVPG45_RS04465 | | | G > del 1 |
| NC_014760 | 934803 | MBOVPG45_RS04465 | | C>T | |
| NC_014760 | 934847 | MBOVPG45_RS04465 | | | A>G |
| NC_014760 | 934849 | MBOVPG45_RS04465 | | | A>C |
| NC_014760 | 934869 | MBOVPG45_RS04465 | | | C>T |
| NC_014760 | 934959 | MBOVPG45_RS04465 | | | G > del 1 |
| NC_014760 | 937718 | MBOVPG45_RS04470 | | | TG>CC |
| NC_014760 | 937722 | MBOVPG45_RS04470 | | | AG>GT |
| NC_014760 | 937794 | MBOVPG45_RS04470 | | | A>T |
| NC_014760 | 938130 | MBOVPG45_RS04470 | | | A>T |
| NC_014760 | 938269 | MBOVPG45_RS04470 | | AT>TC | |
| NC_014760 | 938304 | MBOVPG45_RS04470 | | C>T | |
| NC_014760 | 938341 | MBOVPG45_RS04470 | | | AT>TC |
| NC_014760 | 938447 | MBOVPG45_RS04470 | | TTTTTG > del 6 | |
| NC_014760 | 938456 | MBOVPG45_RS04470 | | T>A | |
| NC_014760 | 938459 | MBOVPG45_RS04470 | | TC>AT | T>C |
| NC_014760 | 938683 | MBOVPG45_RS04470 | | A>C | |
| NC_014760 | 938737 | MBOVPG45_RS04470 | | T>A | |
| NC_014760 | 938740 | MBOVPG45_RS04470 | | A>T | |
| NC_014760 | 938809 | MBOVPG45_RS04470 | | T > del 1 | |
| NC_014760 | 939874 | MBOVPG45_RS04050 | | | G>A |
| NC_014760 | 940058 | MBOVPG45_RS04050 | | | GC>AT |

| | | | | | |
|---|---|---|---|---|---|
| NC_014760 | 940364 | MBOVPG45_RS04055 | | | C>T |
| NC_014760 | 940612 | MBOVPG45_RS04055 | | | A>T |
| NC_014760 | 941162 | MBOVPG45_RS04055 | | | T>C |
| NC_014760 | 941166 | MBOVPG45_RS04055 | | | T>G |
| NC_014760 | 942397 | MBOVPG45_RS04065 | | | G>A |
| NC_014760 | 943606 | MBOVPG45_RS04070 | | | T>A |
| NC_014760 | 943620 | MBOVPG45_RS04070 | | | C>T |
| NC_014760 | 943637 | MBOVPG45_RS04070 | | | A>T |
| NC_014760 | 943665 | MBOVPG45_RS04070 | | | - > ins A |
| NC_014760 | 943668 | MBOVPG45_RS04070 | | | C>A |
| NC_014760 | 945959 | MBOVPG45_RS04085 | | | G>T |
| NC_014760 | 945994 | MBOVPG45_RS04085 | | | A>T |
| NC_014760 | 946001 | MBOVPG45_RS04085 | | | C>T |
| NC_014760 | 946084 | MBOVPG45_RS04085 | | - > ins T | |
| NC_014760 | 946967 | MBOVPG45_RS04090 | | | T>C |
| NC_014760 | 947105 | MBOVPG45_RS04090 | | | GA>AT |
| NC_014760 | 947119 | MBOVPG45_RS04090 | | G>A | |
| NC_014760 | 947129 | MBOVPG45_RS04090 | | | AT>GA |
| NC_014760 | 947191 | MBOVPG45_RS04090 | | | A>G |
| NC_014760 | 947249 | MBOVPG45_RS04090 | | | AT>GA |
| NC_014760 | 947369 | MBOVPG45_RS04090 | | | GA>AT |
| NC_014760 | 949632 | MBOVPG45_RS04110 | | | A>G |
| NC_014760 | 954349 | MBOVPG45_RS04120 | | | G>A |
| NC_014760 | 954358 | MBOVPG45_RS04120 | | | C>A |
| NC_014760 | 954382 | MBOVPG45_RS04120 | | | A>G |
| NC_014760 | 954426 | MBOVPG45_RS04120 | | | G>A |
| NC_014760 | 954741 | MBOVPG45_RS04120 | | | A>G |
| NC_014760 | 954762 | MBOVPG45_RS04120 | | | G>A |
| NC_014760 | 954862 | MBOVPG45_RS04120 | | | C>T |
| NC_014760 | 955262 | MBOVPG45_RS04120 | | | T>A |
| NC_014760 | 955291 | MBOVPG45_RS04120 | | | T>C |
| NC_014760 | 955295 | MBOVPG45_RS04120 | | | T>A |
| NC_014760 | 955374 | MBOVPG45_RS04125 | | | A>G |
| NC_014760 | 955383 | MBOVPG45_RS04125 | | | C>T |
| NC_014760 | 955396 | MBOVPG45_RS04125 | | | G>C |
| NC_014760 | 955399 | MBOVPG45_RS04125 | | | C>T |
| NC_014760 | 955410 | MBOVPG45_RS04125 | | | G>A |
| NC_014760 | 955420 | MBOVPG45_RS04125 | | | T>C |
| NC_014760 | 955497 | MBOVPG45_RS04125 | | | T>C |
| NC_014760 | 955603 | MBOVPG45_RS04125 | | | T>C |
| NC_014760 | 956001 | gatB | | | T>A |
| NC_014760 | 956070 | gatB | | | T>C |
| NC_014760 | 956076 | gatB | | | C>T |
| NC_014760 | 956269 | gatB | | | T>G |
| NC_014760 | 956278 | gatB | | | T>C |
| NC_014760 | 956651 | gatB | | | C>A |
| NC_014760 | 957002 | gatB | | | A>T |

| | | | | | |
|---|---|---|---|---|---|
| NC_014760 | 957030 | gatB | | | T>C |
| NC_014760 | 957458 | gatA | | | G>T |
| NC_014760 | 957656 | gatA | | | T>C |
| NC_014760 | 957764 | gatA | | | CA>TG |
| NC_014760 | 957899 | gatA | | | T>A |
| NC_014760 | 957907 | gatA | | | T>A |
| NC_014760 | 957939 | gatA | | | T>A |
| NC_014760 | 958067 | gatA | | | T>C |
| NC_014760 | 958307 | gatA | | | A>T |
| NC_014760 | 958689 | MBOVPG45_RS04140 | | | A>T |
| NC_014760 | 958715 | MBOVPG45_RS04140 | | | C>A |
| NC_014760 | 958889 | MBOVPG45_RS04140 | | | GA>AT |
| NC_014760 | 958948 | MBOVPG45_RS04140 | | | C>T |
| NC_014760 | 959225 | MBOVPG45_RS04145 | | | G>A |
| NC_014760 | 959334 | MBOVPG45_RS04145 | | | C>T |
| NC_014760 | 959484 | MBOVPG45_RS04145 | | | T>C |
| NC_014760 | 959498 | MBOVPG45_RS04145 | | | A>G |
| NC_014760 | 959511 | MBOVPG45_RS04145 | | | C>T |
| NC_014760 | 959538 | MBOVPG45_RS04145 | | | T>A |
| NC_014760 | 959658 | MBOVPG45_RS04145 | | | T>C |
| NC_014760 | 959730 | MBOVPG45_RS04145 | | | TC>CT |
| NC_014760 | 965875 | MBOVPG45_RS04185 | | | G>A |
| NC_014760 | 965899 | MBOVPG45_RS04185 | | | G>A |
| NC_014760 | 965909 | MBOVPG45_RS04185 | | | C>T |
| NC_014760 | 966273 | MBOVPG45_RS04185 | | | T>A |
| NC_014760 | 966639 | rbfA | | | A>G |
| NC_014760 | 967387 | MBOVPG45_RS04200 | | | A>C |
| NC_014760 | 967763 | MBOVPG45_RS04200 | | | G>A |
| NC_014760 | 967770 | MBOVPG45_RS04200 | | | G>A |
| NC_014760 | 968069 | MBOVPG45_RS04200 | | A > del 1 | |
| NC_014760 | 968085 | MBOVPG45_RS04200 | | | A>G |
| NC_014760 | 968097 | MBOVPG45_RS04200 | | | G>C |
| NC_014760 | 969173 | MBOVPG45_RS04210 | | | C>A |
| NC_014760 | 969241 | MBOVPG45_RS04210 | | | A>G |
| NC_014760 | 969325 | MBOVPG45_RS04210 | | | A>G |
| NC_014760 | 969553 | MBOVPG45_RS04210 | | | T>C |
| NC_014760 | 969720 | MBOVPG45_RS04210 | | | C>T |
| NC_014760 | 969977 | MBOVPG45_RS04210 | | | C>T |
| NC_014760 | 969985 | MBOVPG45_RS04210 | | | C>T |
| NC_014760 | 970093 | MBOVPG45_RS04210 | | | C>T |
| NC_014760 | 970262 | MBOVPG45_RS04210 | | | T>G |
| NC_014760 | 970301 | MBOVPG45_RS04215 | | | C>T |
| NC_014760 | 970346 | MBOVPG45_RS04215 | | | G>A |
| NC_014760 | 972044 | MBOVPG45_RS04230 | | | A>G |
| NC_014760 | 972501 | prmC | | | T>C |
| NC_014760 | 972552 | prmC | | | T>C |
| NC_014760 | 972789 | prmC | | | C>T |

| | | | | | |
|---|---|---|---|---|---|
| NC_014760 | 974517 | MBOVPG45_RS04245 | | | T>C |
| NC_014760 | 974574 | MBOVPG45_RS04245 | | | T>A |
| NC_014760 | 974700 | MBOVPG45_RS04245 | | | T>C |
| NC_014760 | 974738 | MBOVPG45_RS04245 | | | C>T |
| NC_014760 | 974741 | MBOVPG45_RS04245 | | | T>C |
| NC_014760 | 974766 | MBOVPG45_RS04245 | | | A>C |
| NC_014760 | 974775 | MBOVPG45_RS04245 | | | G>A |
| NC_014760 | 976131 | MBOVPG45_RS04250 | | | T>C |
| NC_014760 | 976184 | MBOVPG45_RS04250 | | | C>T |
| NC_014760 | 976191 | MBOVPG45_RS04250 | | | T>A |
| NC_014760 | 976211 | MBOVPG45_RS04250 | | | T>C |
| NC_014760 | 976226 | MBOVPG45_RS04250 | | | C>T |
| NC_014760 | 976501 | MBOVPG45_RS04255 | DNA topoisomerase subunit B ) | | A>T |
| NC_014760 | 976511 | MBOVPG45_RS04255 | DNA topoisomerase subunit B ) | | G>A |
| NC_014760 | 976954 | MBOVPG45_RS04255 | DNA topoisomerase subunit B ) | | G>A |
| NC_014760 | 976988 | MBOVPG45_RS04255 | DNA topoisomerase subunit B ) | | C>T |
| NC_014760 | 977608 | MBOVPG45_RS04255 | DNA topoisomerase subunit B ) | | G>A |
| NC_014760 | 978153 | MBOVPG45_RS04255 | DNA topoisomerase subunit B ) | | TG>CA |
| NC_014760 | 978606 | MBOVPG45_RS04260 | | | A>G |
| NC_014760 | 978825 | MBOVPG45_RS04260 | | | C>T |
| NC_014760 | 978986 | smpB | | | T>C |
| NC_014760 | 979282 | smpB | | | A>G |
| NC_014760 | 980663 | MBOVPG45_RS04275 | | | T>G |
| NC_014760 | 980670 | MBOVPG45_RS04275 | | | - > ins TC |
| NC_014760 | 980671 | MBOVPG45_RS04275 | | | G>T |
| NC_014760 | 980673 | MBOVPG45_RS04275 | | | CT>TC |
| NC_014760 | 980695 | MBOVPG45_RS04275 | | | T>C |
| NC_014760 | 980715 | MBOVPG45_RS04275 | | | C>T |
| NC_014760 | 980744 | MBOVPG45_RS04275 | | | A>T |
| NC_014760 | 980788 | MBOVPG45_RS04275 | | | T>A |
| NC_014760 | 980815 | MBOVPG45_RS04275 | | | G>A |
| NC_014760 | 980817 | MBOVPG45_RS04275 | | | TT>AA |
| NC_014760 | 980824 | MBOVPG45_RS04275 | | | A>G |
| NC_014760 | 980863 | MBOVPG45_RS04275 | | | T>A |
| NC_014760 | 980865 | MBOVPG45_RS04275 | | | TC>CT |

| | | | | | |
|---|---|---|---|---|---|
| NC_014760 | 980906 | MBOVPG45_RS04275 | | | G>T |
| NC_014760 | 980917 | MBOVPG45_RS04275 | | | C>T |
| NC_014760 | 980931 | MBOVPG45_RS04275 | | | C>A |
| NC_014760 | 980971 | MBOVPG45_RS04275 | | | T>C |
| NC_014760 | 980990 | MBOVPG45_RS04275 | | | T>A |
| NC_014760 | 981005 | MBOVPG45_RS04275 | | | T>A |
| NC_014760 | 981011 | MBOVPG45_RS04275 | | | A>T |
| NC_014760 | 981058 | MBOVPG45_RS04275 | | | T>C |
| NC_014760 | 981060 | MBOVPG45_RS04275 | | | GT>TC |
| NC_014760 | 981077 | MBOVPG45_RS04275 | | | T>A |
| NC_014760 | 981079 | MBOVPG45_RS04275 | | | C>T |
| NC_014760 | 981082 | MBOVPG45_RS04275 | | | T>A |
| NC_014760 | 981087 | MBOVPG45_RS04275 | | | A>G |
| NC_014760 | 981110 | MBOVPG45_RS04275 | | T>A | |
| NC_014760 | 981112 | MBOVPG45_RS04275 | | T>C | |
| NC_014760 | 981127 | MBOVPG45_RS04275 | | C>T | C>T |
| NC_014760 | 981134 | MBOVPG45_RS04275 | | | A>T |
| NC_014760 | 981135 | MBOVPG45_RS04275 | | T>G | |
| NC_014760 | 981141 | MBOVPG45_RS04275 | | TTATA>CCTCT | |
| NC_014760 | 981152 | MBOVPG45_RS04275 | | AGT>GCC | |
| NC_014760 | 981156 | MBOVPG45_RS04275 | | C>A | |
| NC_014760 | 981163 | MBOVPG45_RS04275 | | TAGA>ACTT | |
| NC_014760 | 981243 | MBOVPG45_RS04275 | | | G>T |
| NC_014760 | 981285 | MBOVPG45_RS04275 | | | T>G |
| NC_014760 | 981288 | MBOVPG45_RS04275 | | | T>C |
| NC_014760 | 981298 | MBOVPG45_RS04275 | | | C>T |
| NC_014760 | 981315 | MBOVPG45_RS04275 | | | T>G |
| NC_014760 | 981344 | MBOVPG45_RS04275 | | | C>A |
| NC_014760 | 981358 | MBOVPG45_RS04275 | | | T>C |
| NC_014760 | 981394 | MBOVPG45_RS04275 | | | T>C |
| NC_014760 | 981400 | MBOVPG45_RS04275 | | | CG>TC |
| NC_014760 | 981403 | MBOVPG45_RS04275 | | | C>G |
| NC_014760 | 981424 | MBOVPG45_RS04275 | | | C>T |
| NC_014760 | 981508 | MBOVPG45_RS04275 | | | A>T |
| NC_014760 | 981553 | MBOVPG45_RS04275 | | | T>C |
| NC_014760 | 981821 | MBOVPG45_RS04280 | | | G > del 1 |
| NC_014760 | 982458 | MBOVPG45_RS04285 | | | C>T |
| NC_014760 | 984371 | MBOVPG45_RS04300 | | G>A | |
| NC_014760 | 984390 | MBOVPG45_RS04300 | | | C>T |
| NC_014760 | 985381 | mgtA | | | T>G |
| NC_014760 | 985425 | mgtA | | | C>T |
| NC_014760 | 985446 | mgtA | | | T>C |
| NC_014760 | 985534 | mgtA | | | T>A |
| NC_014760 | 986482 | mgtA | | | C>G |
| NC_014760 | 986524 | mgtA | | | T>C |
| NC_014760 | 987109 | mgtA | | A>G | |
| NC_014760 | 987655 | mgtA | | | T>A |

| | | | | | |
|---|---|---|---|---|---|
| NC_014760 | 987798 | mgtA | | | T>C |
| NC_014760 | 988794 | MBOVPG45_RS04310 | | | T>C |
| NC_014760 | 988880 | MBOVPG45_RS04310 | | | A>T |
| NC_014760 | 989006 | MBOVPG45_RS04310 | | | C>T |
| NC_014760 | 989072 | MBOVPG45_RS04310 | | | A>G |
| NC_014760 | 989075 | MBOVPG45_RS04310 | | | G>A |
| NC_014760 | 989187 | MBOVPG45_RS04315 | | | C>T |
| NC_014760 | 989763 | MBOVPG45_RS04315 | | | T>G |
| NC_014760 | 990104 | MBOVPG45_RS04315 | | | T>G |
| NC_014760 | 990153 | MBOVPG45_RS04315 | | | CG>AA |
| NC_014760 | 990156 | MBOVPG45_RS04315 | | | T>C |
| NC_014760 | 990516 | MBOVPG45_RS04315 | | | G>T |
| NC_014760 | 990619 | MBOVPG45_RS04315 | | | T>A |
| NC_014760 | 990759 | MBOVPG45_RS04315 | | | A>T |
| NC_014760 | 991185 | MBOVPG45_RS04315 | | | AC>GT |
| NC_014760 | 991308 | MBOVPG45_RS04315 | | | T>C |
| NC_014760 | 991667 | MBOVPG45_RS04315 | | | G>A |
| NC_014760 | 991704 | MBOVPG45_RS04315 | | | A>G |
| NC_014760 | 991872 | MBOVPG45_RS04315 | | | A>T |
| NC_014760 | 992403 | MBOVPG45_RS04315 | | | T>C |
| NC_014760 | 992676 | MBOVPG45_RS04315 | | | G>C |
| NC_014760 | 992741 | MBOVPG45_RS04315 | | | C>T |
| NC_014760 | 992874 | MBOVPG45_RS04315 | | | T>A |
| NC_014760 | 992900 | MBOVPG45_RS04315 | | | A>G |
| NC_014760 | 992907 | MBOVPG45_RS04315 | | | C>T |
| NC_014760 | 992916 | MBOVPG45_RS04315 | | | G>A |
| NC_014760 | 993032 | MBOVPG45_RS04315 | | | T>G |
| NC_014760 | 993047 | MBOVPG45_RS04315 | | | C>T |
| NC_014760 | 993162 | MBOVPG45_RS04315 | | | GC>CT |
| NC_014760 | 993177 | MBOVPG45_RS04315 | | | A>G |
| NC_014760 | 993349 | MBOVPG45_RS04315 | | | AA>TC |
| NC_014760 | 993356 | MBOVPG45_RS04315 | | | T>C |
| NC_014760 | 993573 | MBOVPG45_RS04315 | | | C>T |
| NC_014760 | 993618 | MBOVPG45_RS04315 | | | T>C |
| NC_014760 | 993696 | MBOVPG45_RS04315 | | | A>C |
| NC_014760 | 993709 | MBOVPG45_RS04315 | | | AG>GA |
| NC_014760 | 993763 | MBOVPG45_RS04315 | | | AA>GG |
| NC_014760 | 993770 | MBOVPG45_RS04315 | | | G>A |
| NC_014760 | 993793 | MBOVPG45_RS04315 | | | T>A |
| NC_014760 | 993803 | MBOVPG45_RS04315 | | | G>T |
| NC_014760 | 994017 | MBOVPG45_RS04315 | | | T>C |
| NC_014760 | 994027 | MBOVPG45_RS04315 | | | G>T |
| NC_014760 | 994029 | MBOVPG45_RS04315 | | | C>T |
| NC_014760 | 994063 | MBOVPG45_RS04315 | | | GC>TT |
| NC_014760 | 994076 | MBOVPG45_RS04315 | | | GT>TC |
| NC_014760 | 994199 | MBOVPG45_RS04315 | | | C>T |
| NC_014760 | 994227 | MBOVPG45_RS04315 | | | C>T |

| | | | | | | |
|---|---|---|---|---|---|---|
| NC_014760 | 994235 | MBOVPG45_RS04315 | | | | T>A |
| NC_014760 | 994427 | MBOVPG45_RS04315 | | | G>A | |
| NC_014760 | 994440 | MBOVPG45_RS04315 | | | | C>T |
| NC_014760 | 994447 | MBOVPG45_RS04315 | | | | A>T |
| NC_014760 | 994452 | MBOVPG45_RS04315 | | | | C>T |
| NC_014760 | 994575 | MBOVPG45_RS04315 | | | | T>A |
| NC_014760 | 994602 | MBOVPG45_RS04315 | | | | T>C |
| NC_014760 | 994773 | MBOVPG45_RS04315 | | | | T>C |
| NC_014760 | 994849 | MBOVPG45_RS04315 | | | | T>A |
| NC_014760 | 994851 | MBOVPG45_RS04315 | | | | C>T |
| NC_014760 | 994950 | MBOVPG45_RS04315 | | | | T>A |
| NC_014760 | 995010 | MBOVPG45_RS04315 | | | | T>C |
| NC_014760 | 995076 | MBOVPG45_RS04315 | | | | C>T |
| NC_014760 | 995112 | MBOVPG45_RS04315 | | | | C>T |
| NC_014760 | 995114 | MBOVPG45_RS04315 | | | | G>A |
| NC_014760 | 995182 | MBOVPG45_RS04315 | | | | C>A |
| NC_014760 | 995429 | MBOVPG45_RS04315 | | | | C>T |
| NC_014760 | 995756 | MBOVPG45_RS04315 | | | | T>C |
| NC_014760 | 995786 | MBOVPG45_RS04315 | | | | G>T |
| NC_014760 | 995817 | MBOVPG45_RS04315 | | | | T>C |
| NC_014760 | 995894 | MBOVPG45_RS04315 | | | | G>A |
| NC_014760 | 995923 | MBOVPG45_RS04315 | | | | A>T |
| NC_014760 | 995944 | MBOVPG45_RS04315 | | | | A>T |
| NC_014760 | 995954 | MBOVPG45_RS04315 | | | | C>T |
| NC_014760 | 995963 | MBOVPG45_RS04315 | | | | A>T |
| NC_014760 | 996032 | MBOVPG45_RS04315 | | | | G>T |
| NC_014760 | 996161 | MBOVPG45_RS04315 | | | | T>G |
| NC_014760 | 996174 | MBOVPG45_RS04315 | | | | G>T |
| NC_014760 | 996198 | MBOVPG45_RS04315 | | | | T>C |
| NC_014760 | 996235 | MBOVPG45_RS04315 | | | | TG>AA |
| NC_014760 | 996238 | MBOVPG45_RS04315 | | | | G>T |
| NC_014760 | 996254 | MBOVPG45_RS04315 | | | | T>C |
| NC_014760 | 996282 | MBOVPG45_RS04315 | | | | C>T |
| NC_014760 | 996310 | MBOVPG45_RS04315 | | | | T>A |
| NC_014760 | 996401 | MBOVPG45_RS04315 | | | | A>T |
| NC_014760 | 996427 | MBOVPG45_RS04315 | | | | T>C |
| NC_014760 | 996457 | MBOVPG45_RS04315 | | | | A>T |
| NC_014760 | 996562 | MBOVPG45_RS04315 | | | | C>A |
| NC_014760 | 996684 | MBOVPG45_RS04315 | | | | T>C |
| NC_014760 | 996686 | MBOVPG45_RS04315 | | | | C>T |
| NC_014760 | 996720 | MBOVPG45_RS04315 | | | | T>C |
| NC_014760 | 996742 | MBOVPG45_RS04315 | | | | A>T |
| NC_014760 | 996762 | MBOVPG45_RS04315 | | | | TG>CA |
| NC_014760 | 996846 | MBOVPG45_RS04315 | | | | G>T |
| NC_014760 | 996865 | MBOVPG45_RS04315 | | | | TTC > del 3 |
| NC_014760 | 996875 | MBOVPG45_RS04315 | | | | T>C |
| NC_014760 | 996894 | MBOVPG45_RS04315 | | | | C>T |

| | | | | | |
|---|---|---|---|---|---|
| NC_014760 | 996965 | MBOVPG45_RS04315 | | | C>T |
| NC_014760 | 997012 | MBOVPG45_RS04315 | | | A>T |
| NC_014760 | 997722 | hflB | | C>T | |
| NC_014760 | 999101 | hflB | | | A>G |
| NC_014760 | 999168 | hflB | | | G>A |
| NC_014760 | 999725 | tilS | | | G>T |
| NC_014760 | 999771 | tilS | | | AT>TC |
| NC_014760 | 999855 | tilS | | | T>C |
| NC_014760 | 999866 | tilS | | | G>T |
| NC_014760 | 999880 | tilS | | | C>T |
| NC_014760 | 999959 | tilS | | | G>T |
| NC_014760 | 999983 | tilS | | | CT>AG |
| NC_014760 | 1000035 | tilS | | | C>T |
| NC_014760 | 1000569 | MBOVPG45_RS04330 | | | A>C |
| NC_014760 | 1001054 | MBOVPG45_RS04335 | | | C>T |
| NC_014760 | 1001152 | MBOVPG45_RS04335 | | | C>T |
| NC_014760 | 1001314 | MBOVPG45_RS04335 | | | C>G |
| NC_014760 | 1001438 | MBOVPG45_RS04335 | | | T>C |
| NC_014760 | 1002098 | MBOVPG45_RS04335 | | | T>C |
| NC_014760 | 1002380 | MBOVPG45_RS04335 | | | G>A |
| NC_014760 | 1002503 | MBOVPG45_RS04335 | | | GC>AT |
| NC_014760 | 1002623 | MBOVPG45_RS04335 | | | T>C |
| NC_014760 | 1002755 | MBOVPG45_RS04335 | | | G>T |
