## Supplemental Table 2 for "Identification of antimicrobial resistance-associated genes through whole genome sequencing of high and low MIC isolates of *Mycoplasma bovis*"

**Table S2 Summary of Nonsynonymous mutations of isolate 1982-M6152 (1982) and 19-043682 (2019)**

### S2-1 Potential AMR mutation sites:

| Strain | 1982 (Y/N) | 2019 (Y/N) | Ref Pos | Gene Name | PG45 Gene size (nt) | #Mutations | Mutation s/100 nt | Disrupted (Y/N) | Gene Description | Functional Role: | Notes: |
|---|---|---|---|---|---|---|---|---|---|---|---|
| 2019 | N | Y | 375 | dnaA | 1401 | 1 | 0.07 | N | chromosomal replication initiator protein DnaA | Replication | **Rifamycin inhibitor** |
| 2019 | N | Y | 1717 | MBOVPG45_RS00010 | 1110 | 1 | 0.09 | N | DNA polymerase III subunit beta | Replication | |
| 2019 | N | Y | 15908 | MBOVPG45_RS00090 | 1605 | 1 | 0.06 | N | ABC transporter ATP-binding protein | ABC transporter system | **ABC transporter system involved in efflux pump resistance** |
| Both | Y | Y | 22594 | MBOVPG45_RS00115 | 1872 | 2 | 0.11 | Both | peptidase S41 | Protein Synthesis | |
| 2019 | N | Y | 36577 | MBOVPG45_RS00180 | 1086 | 2 | 0.18 | N | ABC transporter permease | ABC transporter system | |
| 2019 | N | Y | 53148 | MBOVPG45_RS00205 | 1242 | 9 | 0.72 | N | class I tRNA ligase family protein | Protein Synthesis | |
| 2019 | N | Y | 54319 | rlmB | 702 | 5 | 0.71 | N | 23S rRNA (guanosine(2251)-2'-O)-methyltransferase RlmB | Protein Synthesis | **rlmA confers resistance to mycinamicin, tylosin and lincosamides** |
| 2019 | N | Y | 54964 | MBOVPG45_RS00215 | 585 | 5 | 0.85 | 2019 | sigma-70 family RNA polymerase sigma factor | Transcription | **Rifamycin inhibitor** |
| 2019 | N | Y | 55733 | secE | 237 | 1 | 0.42 | N | preprotein translocase subunit SecE | Protein transport | |
| 2019 | N | Y | 61321 | MBOVPG45_RS00245 | 552 | 3 | 0.54 | N | ribosome recycling factor (frr) | Protein Synthesis | |
| 2019 | N | Y | 67139 | rsmA | 789 | 6 | 0.76 | N | 16S rRNA (adenine(1518)-N(6)/adenine(1519)-N(6))-dimethyltransferase RsmA | Protein Synthesis | **16S methyltransferases mutations confer aminoglycoside resistance** |
| 2019 | N | Y | 68835 | mnmE | 1338 | 4 | 0.30 | N | tRNA uridine-5-carboxymethylaminomethyl(34) synthesis GTPase MnmE | Protein Synthesis | |
| 2019 | N | Y | 72682 | serS | 1269 | 1 | 0.08 | N | serine--tRNA ligase | Protein Synthesis | |
| Both | Y | Y | 75564 | yidC | 2166 | 10 | 0.46 | 1982 | membrane protein insertase YidC | Protein transport | |
| Both | Y | Y | 81331 | MBOVPG45_RS00350 | 4377 | 15 | 0.34 | N | PolC-type DNA polymerase III | Replication | |
| 2019 | N | Y | 86893 | MBOVPG45_RS00365 | 1224 | 3 | 0.25 | N | replication-associated recombination protein A | Replication, DNA recombination | |
| 2019 | N | Y | 87536 | pheS | 966 | 2 | 0.21 | N | phenylalanine--tRNA ligase subunit alpha | Protein Synthesis | |
| 2019 | N | Y | 89028 | MBOVPG45_RS00380 | 2187 | 18 | 0.82 | N | phenylalanine--tRNA ligase subunit beta | Protein Synthesis | |
| 2019 | N | Y | 93312 | MBOVPG45_RS00395 | 1161 | 10 | 0.86 | N | aminotransferase class V-fold PLP-dependent enzyme | Protein Synthesis | |
| 2019 | N | Y | 94484 | MBOVPG45_RS00400 | 429 | 7 | 1.63 | N | iron-sulfur cluster assembly scaffold protein | Protein assembly | |
| 2019 | N | Y | 94954 | MBOVPG45_RS00405 | 1248 | 6 | 0.48 | N | DNA polymerase IV | Replication | |
| Both | Y | Y | 97237 | MBOVPG45_RS00415 | 768 | 2 | 0.26 | N | rRNA pseudouridine synthase | Protein Synthesis | |
| 2019 | N | Y | 102115 | MBOVPG45_RS00445 | 696 | 1 | 0.14 | N | 50S ribosomal protein L1 | Protein Synthesis | |
| 2019 | N | Y | 105222 | MBOVPG45_RS00465 | 534 | 2 | 0.37 | N | tRNA (cytidine(34)-2'-O)-methyltransferase | Protein Synthesis | |
| 2019 | N | Y | 105633 | MBOVPG45_RS00470 | 732 | 7 | 0.96 | N | RNA methyltransferase | Protein Synthesis | |
| 2019 | N | Y | 106703 | ylqF | 861 | 3 | 0.35 | N | ribosome biogenesis GTPase YlqF | Protein Synthesis | |
| 2019 | N | Y | 122978 | MBOVPG45_RS00555 | 1122 | 2 | 0.18 | N | ABC transporter permease | ABC transporter system | |
| 2019 | N | Y | 126802 | MBOVPG45_RS00570 | 2403 | 4 | 0.17 | N | ATP-binding cassette domain-containing protein | ABC transporter system | |
| 2019 | N | Y | 133930 | MBOVPG45_RS00600 | 2058 | 1 | 0.05 | N | ATP-binding cassette domain-containing protein | ABC transporter system | |
| 2019 | N | Y | 155399 | thiI | 1137 | 4 | 0.35 | N | tRNA 4-thiouridine(8) synthase ThiI | Protein Synthesis | |
| 2019 | N | Y | 170114 | MBOVPG45_RS00740 | 2490 | 7 | 0.28 | 2019 | valine--tRNA ligase | Protein Synthesis | |
| 2019 | N | Y | 180543 | dnaK | 1779 | 2 | 0.11 | N | molecular chaperone DnaK | Protein Synthesis | |
| 2019 | N | Y | 181437 | MBOVPG45_RS00790 | 1284 | 7 | 0.55 | N | methylenetetrahydrofolate--tRNA-(uracil(54)-C(5))-methyltransferase (FADH(2)-oxidizing) TrmFO | DNA methylation | |
| 2019 | N | Y | 187768 | MBOVPG45_RS00815 | 2424 | 9 | 0.37 | N | DEAD/DEAH box helicase family protein | Transcription, Protein synthesis | |
| 2019 | N | Y | 195409 | tig | 1506 | 3 | 0.20 | N | trigger factor | Protein Synthesis | |
| Both | Y | Y | 248926 | MBOVPG45_RS01075 | 1266 | 33 | 2.61 | Both | endonuclease/exonuclease/phosphatase family protein | Cell signalling, DNA repair | |
| 2019 | N | Y | 264264 | MBOVPG45_RS01150 | 1392 | 6 | 0.43 | 2019 | glutamate--tRNA ligase | Protein Synthesis | |
| 2019 | N | Y | 265487 | MBOVPG45_RS01155 | 2040 | 29 | 1.42 | 2019 | peptidase S41 | Protein Synthesis | |
| 2019 | N | Y | 267529 | MBOVPG45_RS01160 | 1932 | 25 | 1.29 | N | peptidase S41 | Protein Synthesis | |
| 2019 | N | Y | 289078 | gyrA | 2760 | 8 | 0.29 | N | DNA gyrase subunit A | Topoisomerase | **fluoroquinolone resistance** |
| 2019 | N | Y | 293726 | rpsD | 597 | 2 | 0.34 | N | 30S ribosomal protein S4 | Protein Synthesis | |
| 2019 | N | Y | 294448 | rpmE | 213 | 1 | 0.47 | N | 50S ribosomal protein L31 | Protein Synthesis | |
| 2019 | N | Y | 294901 | hrcA | 1008 | 1 | 0.10 | N | heat-inducible transcriptional repressor HrcA | Transcription | |
| 2019 | N | Y | 295867 | grpE | 1026 | 4 | 0.39 | N | nucleotide exchange factor GrpE | Protein Synthesis | |
| 2019 | N | Y | 298696 | MBOVPG45_RS01270 | 711 | 2 | 0.28 | N | rRNA pseudouridine synthase | Protein Synthesis | |
| 1982 | Y | N | 304123 | rpsJ | 324 | 1 | 0.31 | N | 30S ribosomal protein S10 | Protein Synthesis | **rpsJ confers tetracycline resistance** |
| Both | Y | Y | 305116 | rplC | 831 | 1 | 0.12 | N | 50S ribosomal protein L3 | Protein Synthesis | |
| 2019 | N | Y | 305204 | rplD | 951 | 10 | 1.05 | N | 50S ribosomal protein L4 | Protein Synthesis | **rplD confers macrolide resistance** |
| 2019 | N | Y | 306754 | rplB | 846 | 1 | 0.12 | N | 50S ribosomal protein L2 | Protein Synthesis | |
| 2019 | N | Y | 307518 | rpsS | 279 | 1 | 0.36 | N | 30S ribosomal protein S19 | Protein Synthesis | |
| 2019 | N | Y | 308038 | rplV | 345 | 1 | 0.29 | N | 50S ribosomal protein L22 | Protein Synthesis | |
| 2019 | N | Y | 308343 | rpsC | 651 | 3 | 0.46 | N | 30S ribosomal protein S3 | Protein Synthesis | |
| 2019 | N | Y | 310225 | MBOVPG45_RS01360 | 327 | 1 | 0.31 | N | 50S ribosomal protein L24 | Protein Synthesis | |
| 2019 | N | Y | 311251 | rpsH | 396 | 1 | 0.25 | N | 30S ribosomal protein S8 | Protein Synthesis | |
| Both | Y | Y | 312554 | rpsE | 696 | 3 | 0.43 | N | 30S ribosomal protein S5 | Protein Synthesis | |
| 2019 | N | Y | 333960 | MBOVPG45_RS01485 | 906 | 2 | 0.22 | N | energy-coupling factor transporter transmembrane protein EcfT | ABC transporter system | |
| 2019 | N | Y | 341084 | obgE | 1269 | 2 | 0.16 | N | GTPase ObgE | Protein Synthesis | |
| 2019 | N | Y | 345146 | MBOVPG45_RS01540 | 1068 | 1 | 0.09 | N | sugar ABC transporter permease | ABC transporter system | |
| 2019 | N | Y | 345580 | MBOVPG45_RS01545 | 2103 | 4 | 0.19 | N | ATP-binding cassette domain-containing protein | ABC transporter system | |
| 2019 | N | Y | 371209 | MBOVPG45_RS01640 | 1620 | 5 | 0.31 | N | arginine--tRNA ligase | Protein Synthesis | |
| 2019 | N | Y | 380382 | lepA | 1794 | 4 | 0.22 | N | elongation factor 4 | Protein Synthesis | |
| 2019 | N | Y | 392689 | MBOVPG45_RS01720 | 1119 | 1 | 0.09 | N | ABC transporter permease subunit | ABC transporter system | |
| 2019 | N | Y | 401183 | MBOVPG45_RS01770 | 705 | 1 | 0.14 | N | ABC transporter ATP-binding protein | ABC transporter system | |
| Both | Y | Y | 401939 | MBOVPG45_RS01775 | 1863 | 8 | 0.43 | N | ABC transporter permease | ABC transporter system | |
| 2019 | N | Y | 411488 | MBOVPG45_RS01820 | 963 | 7 | 0.73 | N | DNA polymerase III subunit delta | Replication | |
| 2019 | N | Y | 416161 | whiA | 882 | 3 | 0.34 | N | DNA-binding protein WhiA | Replication | |
| 2019 | N | Y | 419734 | ligA | 1965 | 2 | 0.10 | N | DNA ligase (NAD(+)) LigA | Replication, DNA repair | |
| 2019 | N | Y | 454173 | asnS | 1359 | 1 | 0.07 | N | asparagine--tRNA ligase | Protein Synthesis | |
| 2019 | N | Y | 462062 | MBOVPG45_RS02005 | 1797 | 5 | 0.28 | N | ABC transporter ATP-binding protein | ABC transporter system | |

| | | | | | | | | | | | |
|---|---|---|---|---|---|---|---|---|---|---|---|
| 2019 | N | Y | 477575 | tsaE | 405 | 2 | 0.49 | N | tRNA (adenosine(37)-N6)-threonylcarbamoyltransferase complex ATPase subunit type 1 TsaE | Protein Synthesis | |
| 2019 | N | Y | 477995 | MBOVPG45_RS02070 | 558 | 4 | 0.72 | N | tRNA (adenosine(37)-N6)-threonylcarbamoyltransferase complex dimerization subunit type 1 TsaB | Protein Synthesis | |
| 2019 | N | Y | 478888 | tsaD | 930 | 3 | 0.32 | N | tRNA (adenosine(37)-N6)-threonylcarbamoyltransferase complex transferase subunit TsaD | Protein Synthesis | |
| 2019 | N | Y | 487031 | MBOVPG45_RS02105 | 1959 | 1 | 0.05 | N | peptidase S41 | Protein Synthesis | |
| 2019 | N | Y | 504209 | trpS | 996 | 1 | 0.10 | N | tryptophan--tRNA ligase | Protein Synthesis | |
| Both | Y | Y | 505264 | MBOVPG45_RS02170 | 1749 | 9 | 0.51 | 2019 | threonine--tRNA ligase | Protein Synthesis | |
| 2019 | N | Y | 523679 | MBOVPG45_RS02255 | 2340 | 16 | 0.68 | N | leucine--tRNA ligase | Protein Synthesis | |
| Both | Y | Y | 527473 | ssb | 561 | 5 | 0.89 | N | single-stranded DNA-binding protein | Replication, DNA recombination, DNA repair | |
| 2019 | N | Y | 529969 | MBOVPG45_RS02280 | 684 | 4 | 0.58 | N | 16S rRNA (uracil(1498)-N(3))-methyltransferase | Protein Synthesis | |
| 2019 | N | Y | 531421 | MBOVPG45_RS02290 | 1143 | 2 | 0.17 | N | cell division protein FtsZ | Replication | |
| 2019 | N | Y | 534120 | rsmH | 903 | 5 | 0.55 | N | 16S rRNA (cytosine(1402)-N(4))-methyltransferase RsmH | Protein Synthesis | |
| 2019 | N | Y | 538752 | dcm | 948 | 7 | 0.74 | N | DNA cytosine methyltransferase | DNA methylation | |
| 2019 | N | Y | 545064 | MBOVPG45_RS02345 | 687 | 5 | 0.73 | N | NERD domain-containing protein | Replication | |
| 2019 | N | Y | 566912 | MBOVPG45_RS02435 | 432 | 2 | 0.46 | N | single-stranded DNA-binding protein | Replication, DNA recombination, DNA repair | |
| Both | Y | Y | 582733 | alaS | 1136 | 9 | 0.79 | 2019 | alanine--tRNA ligase | Protein Synthesis | **novobiocin resistance** |
| 2019 | N | Y | 585222 | mnmA | 1125 | 5 | 0.44 | N | tRNA 2-thiouridine(34) synthase MnmA | Protein Synthesis | |
| 2019 | N | Y | 606351 | MBOVPG45_RS02615 | 1002 | 1 | 0.10 | N | bifunctional oligoribonuclease/PAP phosphatase NrnA | Protein Synthesis | |
| 2019 | N | Y | 613725 | MBOVPG45_RS02640 | 1317 | 1 | 0.08 | N | histidine--tRNA ligase | Protein Synthesis | |
| 2019 | N | Y | 624992 | **topA** | **1842** | **1** | **0.05** | **2019** | **type I DNA topoisomerase** | **Topoisomerase** | |
| 2019 | N | Y | 631133 | MBOVPG45_RS02710 | 1752 | 1 | 0.06 | N | ABC transporter permease subunit | ABC transporter system | |
| 2019 | N | Y | 631191 | MBOVPG45_RS02715 | 933 | 2 | 0.21 | N | ATP-binding cassette domain-containing protein | ABC transporter system | |
| 2019 | N | Y | 636688 | MBOVPG45_RS02730 | 1377 | 2 | 0.15 | N | glycine--tRNA ligase | Protein Synthesis | |
| 2019 | N | Y | 637373 | dnaG | 1941 | 4 | 0.21 | N | DNA primase | Replication | |
| 2019 | N | Y | 639516 | MBOVPG45_RS02740 | 1530 | 1 | 0.07 | N | RNA polymerase sigma factor | Transcription | **Rifamycin inhibitor** |
| 2019 | N | Y | 643759 | MBOVPG45_RS02760 | 1875 | 18 | 0.96 | 2019 | peptidase S41 | Protein Synthesis | |
| 2019 | N | Y | 645973 | MBOVPG45_RS02765 | 897 | 5 | 0.56 | N | elongation factor Ts | Protein Synthesis | |
| 2019 | N | Y | 646646 | rpsB | 954 | 3 | 0.31 | N | 30S ribosomal protein S2 | Protein Synthesis | |
| 2019 | N | Y | 656135 | MBOVPG45_RS02805 | 2088 | 14 | 0.67 | N | peptidase S41 | Protein Synthesis | |
| 2019 | N | Y | 674943 | dnaB | 1476 | 1 | 0.07 | N | replicative DNA helicase | Replication | |
| Both | Y | Y | 682961 | MBOVPG45_RS02905 | 825 | 2 | 0.24 | N | ABC transporter permease subunit | ABC transporter system | |
| Both | Y | Y | 697741 | ileS | 2679 | 6 | 0.22 | 2019 | isoleucine--tRNA ligase | Protein Synthesis | **pseudomonic acid target** |
| 2019 | N | Y | 700827 | MBOVPG45_RS03000 | 1620 | 2 | 0.12 | N | CTP synthase | Nucleotide synthesis | |
| 2019 | N | Y | 730969 | MBOVPG45_RS03145 | 807 | 1 | 0.12 | N | isoleucine--tRNA ligase | Protein Synthesis | |
| 2019 | N | Y | 731350 | MBOVPG45_RS03150 | 1551 | 10 | 0.64 | 2019 | methionine--tRNA ligase | Protein Synthesis | |
| Both | Y | Y | 757435 | parC | 2586 | 19 | 0.73 | N | DNA topoisomerase IV subunit A | Topoisomerase | **fluoroquinolone resistance** |
| 2019 | N | Y | 760101 | parE | 1917 | 4 | 0.21 | N | DNA topoisomerase IV subunit B | Topoisomerase | **fluoroquinolone resistance** |
| 2019 | N | Y | 764252 | rsmD | 549 | 1 | 0.18 | N | 16S rRNA (guanine(966)-N(2))-methyltransferase RsmD | Protein Synthesis | |
| 2019 | N | Y | 767941 | MBOVPG45_RS03325 | 453 | 1 | 0.22 | N | 23S rRNA (pseudouridine(1915)-N(3))-methyltransferase RlmH | Protein Synthesis | |
| 2019 | N | Y | 768041 | mnmG | 1842 | 6 | 0.33 | N | tRNA uridine-5-carboxymethylaminomethyl(34) synthesis enzyme MnmG | Protein Synthesis | |
| Both | Y | Y | 785098 | dnaE | 2934 | 16 | 0.55 | N | DNA polymerase III subunit alpha | Replication | |
| 2019 | N | Y | 798146 | MBOVPG45_RS03425 | 1614 | 1 | 0.06 | N | ATP-binding cassette domain-containing protein | ABC transporter system | **tylosin resistance (GenBank), macrolide resistance (CARD)** |
| 2019 | N | Y | 805063 | MBOVPG45_RS03465 | 1833 | 4 | 0.22 | N | ABC transporter ATP-binding protein | ABC transporter system | |
| 2019 | N | Y | 806434 | MBOVPG45_RS03470 | 1785 | 6 | 0.34 | N | ABC transporter ATP-binding protein | ABC transporter system | |
| 2019 | N | Y | 826479 | rpoC | 4437 | 1 | 0.02 | N | DNA-directed RNA polymerase subunit beta' | Transcription | **Rifamycin inhibitor** |
| 2019 | N | Y | 830647 | MBOVPG45_RS03510 | 3636 | 1 | 0.03 | N | DNA-directed RNA polymerase subunit beta | Transcription | |
| 2019 | N | Y | 832675 | MBOVPG45_RS03525 | 522 | 2 | 0.38 | N | 50S ribosomal protein L10 | Protein Synthesis | |
| 2019 | N | Y | 834722 | lysS | 1470 | 3 | 0.20 | N | lysine--tRNA ligase | Protein Synthesis | |
| 2019 | N | Y | 836887 | MBOVPG45_RS03545 | 2160 | 8 | 0.37 | 2019 | S1 RNA-binding domain-containing protein | Protein Synthesis | |
| 2019 | N | Y | 846198 | rpsP | 291 | 2 | 0.69 | N | 30S ribosomal protein S16 | Protein Synthesis | |
| 2019 | N | Y | 847407 | fmt | 840 | 4 | 0.48 | N | methionyl-tRNA formyltransferase | Protein Synthesis | |
| Both | Y | Y | 853121 | MBOVPG45_RS03600 | 867 | 3 | 0.35 | 1982 | MurR/RpiR family transcriptional regulator | Transcription | |
| 2019 | N | Y | 873133 | MBOVPG45_RS03705 | 813 | 6 | 0.74 | 2019 | carbohydrate ABC transporter permease | ABC transporter system | |
| 2019 | N | Y | 873886 | MBOVPG45_RS03710 | 996 | 1 | 0.10 | N | sugar ABC transporter permease | ABC transporter system | |
| 2019 | N | Y | 880184 | MBOVPG45_RS03730 | 738 | 2 | 0.27 | N | YebC/PmpR family DNA-binding transcriptional regulator | Transcription | |
| 2019 | N | Y | 883839 | rlmD | 1323 | 11 | 0.83 | N | 23S rRNA (uracil(1939)-C(5))-methyltransferase RlmD | Protein Synthesis | |
| Both | Y | Y | 898026 | MBOVPG45_RS03825 | 918 | 1 | 0.11 | N | RluA family pseudouridine synthase | Protein Synthesis | |
| 2019 | N | Y | 900146 | trmB | 618 | 4 | 0.65 | 2019 | tRNA (guanosine(46)-N7)-methyltransferase TrmB | Protein Synthesis | |
| 2019 | N | Y | 904514 | rsmI | 726 | 5 | 0.69 | N | 16S rRNA (cytidine(1402)-2'-O)-methyltransferase | Protein Synthesis | |
| 2019 | N | Y | 905196 | MBOVPG45_RS03870 | 903 | 6 | 0.66 | N | DNA polymerase III subunit delta | Replication | |
| Both | Y | Y | 906098 | MBOVPG45_RS03875 | 648 | 3 | 0.46 | N | dTMP kinase | Nucleotide synthesis | |
| 2019 | N | Y | 907714 | dnaX | 1851 | 6 | 0.32 | N | DNA polymerase III subunit gamma/tau | Replication | |
| 2019 | N | Y | 918571 | MBOVPG45_RS03945 | 723 | 1 | 0.14 | N | adenine phosphoribosyltransferase | Nucleotide synthesis | |
| 2019 | N | Y | 920868 | MBOVPG45_RS03950 | 1806 | 1 | 0.06 | N | translation initiation factor IF-2 | Protein Synthesis | |
| 2019 | N | Y | 921262 | nusA | 1635 | 6 | 0.37 | N | transcription termination/antitermination protein NusA | Transcription | |
| 2019 | N | Y | 956001 | gatB | 1422 | 8 | 0.56 | N | Asp-tRNA(Asn)/Glu-tRNA(Gln) amidotransferase subunit GatB | Protein Synthesis | |
| 2019 | N | Y | 957458 | gatA | 1314 | 8 | 0.61 | N | Asp-tRNA(Asn)/Glu-tRNA(Gln) amidotransferase subunit GatA | Protein Synthesis | |
| 2019 | N | Y | 958689 | MBOVPG45_RS04140 | 306 | 4 | 1.31 | N | glutamyl-tRNA(Gln) amidotransferase | Protein Synthesis | |
| 2019 | N | Y | 965875 | MBOVPG45_RS04185 | 1134 | 4 | 0.35 | N | DnaJ domain-containing protein | Protein Synthesis | |
| 2019 | N | Y | 966639 | rbfA | 342 | 1 | 0.29 | N | 30S ribosome-binding factor RbfA | Protein Synthesis | |
| 2019 | N | Y | 969173 | MBOVPG45_RS04210 | 1236 | 9 | 0.73 | 2019 | tyrosine--tRNA ligase | Protein Synthesis | |
| 2019 | N | Y | 970301 | MBOVPG45_RS04215 | 603 | 2 | 0.33 | N | tRNA-binding protein | Protein Synthesis | |

| | | | | | | | | | | | |
|---|---|---|---|---|---|---|---|---|---|---|---|
| 2019 | N | Y | 972501 | prmC | 723 | 3 | 0.41 | N | peptide chain release factor N(5)-glutamine methyltransferase | DNA methylation | |
| 2019 | N | Y | 974517 | MBOVPG45_RS04245 | 573 | 7 | 1.22 | N | cell division protein Fic | Replication | |
| 2019 | N | Y | 976501 | MBOVPG45_RS04255 | 1968 | 6 | 0.30 | N | type IIA DNA topoisomerase subunit B (gyrB) | Topoisomerase | **fluoroquinolone resistance** |
| 2019 | N | Y | 978986 | smpB | 447 | 2 | 0.45 | N | SsrA-binding protein | Protein Synthesis | |
| 2019 | N | Y | 982458 | MBOVPG45_RS04285 | 960 | 1 | 0.10 | N | LacI family transcriptional regulator | Transcription | |
| 2019 | N | Y | 988794 | MBOVPG45_RS04310 | 1089 | 5 | 0.46 | N | ABC transporter ATP-binding protein | ABC transporter system | |
| **Both** | Y | Y | 989187 | MBOVPG45_RS04315 | 8127 | 90 | 1.11 | N | ABC transporter permease | ABC transporter system | |
| 2019 | N | Y | 999725 | tilS | 861 | 8 | 0.93 | N | tRNA lysidine(34) synthetase TilS | Protein Synthesis | |
| 2019 | N | Y | 1000569 | MBOVPG45_RS04330 | 567 | 1 | 0.18 | N | aminoacyl-tRNA hydrolase | Protein Synthesis | |

### S2-2. Unlikely AMR Candidates:

| Strain | 1982 (Y/N) | 2019 (Y/N) | Ref Pos | Gene Name | PG45 Gene size (nt) | #Mutations | Mutation s/100 nt | Disrupted (Y/N) | Gene Description | |
|---|---|---|---|---|---|---|---|---|---|---|
| 1982 | Y | N | 3257 | MBOVPG45_RS00030 | 786 | 1 | 0.13 | N | alpha/beta hydrolase | Multi-role family |
| 2019 | N | Y | 4439 | MBOVPG45_RS00035 | 792 | 1 | 0.13 | N | alpha/beta hydrolase | Multi-role family |
| 2019 | N | Y | 5197 | MBOVPG45_RS00040 | 1182 | 3 | 0.25 | N | NADH-dependent flavin oxidoreductase | Metabolism |
| 2019 | N | Y | 6185 | MBOVPG45_RS00045 | 1047 | 3 | 0.29 | N | lipoate--protein ligase A | Metabolism |
| 2019 | N | Y | 7253 | MBOVPG45_RS00050 | 1038 | 1 | 0.10 | N | lipoate--protein ligase | Metabolism |
| 2019 | N | Y | 11281 | MBOVPG45_RS00070 | 1413 | 1 | 0.07 | N | IS1634-like element ISMbov2 family transposase | Transposable elements |
| 2019 | N | Y | 36735 | MBOVPG45_RS00185 | 2952 | 9 | 0.30 | N | variable surface lipoprotein | variable surface lipoprotein |
| 2019 | N | Y | 61188 | MBOVPG45_RS00240 | 723 | 1 | 0.14 | N | UMP kinase | Metabolism |
| 2019 | N | Y | 66075 | MBOVPG45_RS00275 | 999 | 7 | 0.70 | N | NAD(P)-binding domain-containing protein | Metabolism |
| 2019 | N | Y | 67846 | MBOVPG45_RS00285 | 762 | 5 | 0.66 | N | TatD family hydrolase | DNA repair |
| 2019 | N | Y | 73768 | MBOVPG45_RS00310 | 1032 | 3 | 0.29 | N | glycosyltransferase family 2 protein | Metabolism |
| 2019 | N | Y | 74822 | rnpA | 324 | 3 | 0.93 | N | ribonuclease P protein component | Rnase |
| 2019 | N | Y | 78149 | MBOVPG45_RS00330 | 981 | 1 | 0.10 | N | lipoate--protein ligase | Metabolism |
| **Both** | Y | Y | 78548 | MBOVPG45_RS00335 | 822 | 6 | 0.73 | N | alpha/beta hydrolase | Multi-role family |
| 2019 | N | Y | 80472 | MBOVPG45_RS00345 | 981 | 3 | 0.31 | N | aspartate--ammonia ligase | Metabolism |
| 2019 | N | Y | 88161 | MBOVPG45_RS00375 | 660 | 4 | 0.61 | N | uracil-DNA glycosylase | DNA repair |
| 2019 | N | Y | 91227 | MBOVPG45_RS00385 | 1266 | 5 | 0.39 | N | serine hydroxymethyltransferase | Methylation |
| 2019 | N | Y | 92255 | pip | 972 | 7 | 0.72 | 2019 | prolyl aminopeptidase | Metabolism |
| 2019 | N | Y | 96345 | MBOVPG45_RS00410 | 1095 | 2 | 0.18 | N | nicotinate-nucleotide adenylyltransferase | Metabolism |
| 2019 | N | Y | 99358 | fba | 876 | 3 | 0.34 | N | class II fructose-1,6-bisphosphate aldolase | Metabolism |
| 2019 | N | Y | 100528 | MBOVPG45_RS00435 | 573 | 3 | 0.52 | N | nuclease | DNA repair |
| 2019 | N | Y | 107976 | MBOVPG45_RS00490 | 963 | 4 | 0.42 | N | HPr kinase/phosphorylase | Metabolism |
| 2019 | N | Y | 108881 | MBOVPG45_RS00495 | 984 | 4 | 0.41 | N | prolipoprotein diacylglyceryl transferase | Posttranslational modification |
| 2019 | N | Y | 110067 | MBOVPG45_RS00500 | 924 | 1 | 0.11 | N | FAD-dependent oxidoreductase | Metabolism |
| **Both** | Y | Y | 115075 | lpdA | 1614 | 2 | 0.12 | N | dihydrolipoyl dehydrogenase | Metabolism |
| 2019 | N | Y | 129778 | MBOVPG45_RS00585 | 1218 | 1 | 0.08 | N | YitT family protein | Stress protein |
| 2019 | N | Y | 132891 | MBOVPG45_RS00595 | 447 | 1 | 0.22 | N | RpiB/LacA/LacB family sugar-phosphate isomerase | Metabolism |
| **Both** | Y | Y | 144605 | MBOVPG45_RS00650 | 2388 | 9 | 0.38 | N | phosphoketolase family protein | Metabolism |
| 2019 | N | Y | 146413 | pepF | 1842 | 4 | 0.22 | N | oligoendopeptidase F | Metabolism |
| 2019 | N | Y | 157477 | MBOVPG45_RS00705 | 1434 | 4 | 0.28 | N | transposase | Transposable elements |
| 2019 | N | Y | 170942 | MBOVPG45_RS00745 | 861 | 2 | 0.23 | N | bifunctional 5,10-methylenetetrahydrofolate dehydrogenase/5,10-methenyltetrahydrofolate cyclohydrolase | Metabolism |
| 2019 | N | Y | 172051 | MBOVPG45_RS00750 | 957 | 4 | 0.42 | N | phosphate acetyltransferase | Metabolism |
| 2019 | N | Y | 173363 | MBOVPG45_RS00755 | 1194 | 2 | 0.17 | 2019 | acetate/propionate family kinase | Metabolism |
| 2019 | N | Y | 174264 | MBOVPG45_RS00760 | 423 | 1 | 0.24 | N | pantetheine-phosphate adenylyltransferase | Metabolism |
| 2019 | N | Y | 174840 | MBOVPG45_RS00765 | 582 | 4 | 0.69 | N | YihA family ribosome biogenesis GTP-binding protein | Metabolism |
| 2019 | N | Y | 178094 | pyk | 1437 | 1 | 0.07 | N | pyruvate kinase | Metabolism |
| **Both** | Y | Y | 183309 | MBOVPG45_RS00800 | 990 | 7 | 0.71 | N | lactate dehydrogenase | Metabolism |
| 2019 | N | Y | 184839 | MBOVPG45_RS00805 | 801 | 6 | 0.75 | N | alpha/beta hydrolase | Multi-role family |
| 2019 | N | Y | 186667 | MBOVPG45_RS00810 | 1029 | 1 | 0.10 | N | IS30 family transposase | Transposable elements |
| 2019 | N | Y | 201195 | MBOVPG45_RS00870 | 2307 | 13 | 0.56 | 2019 | S8 family serine peptidase | Peptidase |
| 2019 | N | Y | 218020 | MBOVPG45_RS00935 | 453 | 3 | 0.66 | 2019 | SocA family protein | Transport protein |
| 2019 | N | Y | 222893 | MBOVPG45_RS00955 | 1029 | 1 | 0.10 | N | IS30-like element ISMbov1 family transposase | Transposable elements |
| 2019 | N | Y | 224139 | MBOVPG45_RS00960 | 1413 | 1 | 0.07 | N | IS1634-like element ISMbov2 family transposase | Transposable elements |
| 2019 | N | Y | 230209 | MBOVPG45_RS00975 | 1434 | 2 | 0.14 | N | transposase | Transposable elements |
| 2019 | N | Y | 252748 | MBOVPG45_RS01085 | 1017 | 1 | 0.10 | N | IS30 family transposase | Transposable elements |
| 2019 | N | Y | 256515 | pgk | 1185 | 2 | 0.17 | N | phosphoglycerate kinase | Metabolism |
| 2019 | N | Y | 257653 | MBOVPG45_RS01110 | 1002 | 2 | 0.20 | N | nicotinate phosphoribosyltransferase | Metabolism |
| 2019 | N | Y | 269648 | MBOVPG45_RS01165 | 555 | 4 | 0.72 | N | 5-formyltetrahydrofolate cyclo-ligase | Metabolism |
| 2019 | N | Y | 273077 | MBOVPG45_RS01185 | 2679 | 33 | 1.23 | 2019 | type I restriction-modification system subunit M | Restriction enzyme |
| **Both** | Y | Y | 275617 | MBOVPG45_RS04625 | 1158 | 6 | 0.52 | N | restriction endonuclease subunit S | Restriction enzyme |
| 2019 | N | Y | 276761 | MBOVPG45_RS01195 | 3198 | 12 | 0.38 | 2019 | type I restriction endonuclease subunit R | Restriction enzyme |
| 2019 | N | Y | 280080 | MBOVPG45_RS01200 | 921 | 1 | 0.11 | N | site-specific integrase | Recombination |
| 2019 | N | Y | 281463 | MBOVPG45_RS01205 | 1194 | 7 | 0.59 | N | restriction endonuclease subunit S | Restriction enzyme |
| 2019 | N | Y | 282460 | MBOVPG45_RS01210 | 1245 | 2 | 0.16 | N | IS3 family transposase | Transposable elements |
| 2019 | N | Y | 299555 | MBOVPG45_RS01275 | 1086 | 6 | 0.55 | N | M42 family metallopeptidase | Metabolism |
| 2019 | N | Y | 302153 | recA | 993 | 2 | 0.20 | N | recombinase RecA | Recombination |
| 2019 | N | Y | 303010 | recU | 492 | 1 | 0.20 | N | Holliday junction resolvase RecU | Recombination |
| 2019 | N | Y | 314969 | MBOVPG45_RS01405 | 993 | 7 | 0.70 | N | histidine acid phosphatase | Multi-role family |
| 2019 | N | Y | 335706 | tpiA | 783 | 2 | 0.26 | N | triose-phosphate isomerase | Metabolism |
| 2019 | N | Y | 336442 | deoC | 669 | 4 | 0.60 | N | deoxyribose-phosphate aldolase | Metabolism |
| 2019 | N | Y | 337141 | MBOVPG45_RS01505 | 1296 | 2 | 0.15 | N | pyrimidine-nucleoside phosphorylase | Metabolism |
| 2019 | N | Y | 338666 | deoD | 702 | 3 | 0.43 | N | purine-nucleoside phosphorylase | Metabolism |
| **Both** | Y | Y | 349035 | MBOVPG45_RS01555 | 2187 | 31 | 1.42 | N | P80 family lipoprotein | Metabolism |
| 2019 | N | Y | 351154 | MBOVPG45_RS01560 | 1434 | 5 | 0.35 | N | transposase | Transposable elements |
| 2019 | N | Y | 353654 | MBOVPG45_RS01570 | 1716 | 6 | 0.35 | N | GIY-YIG nuclease family protein | Multi-role family |
| 2019 | N | Y | 357311 | MBOVPG45_RS01585 | 837 | 2 | 0.24 | N | deoxyribonuclease IV | DNA repair |
| **Both** | Y | Y | 358227 | MBOVPG45_RS01590 | 837 | 5 | 0.60 | N | HAD hydrolase family protein | Metabolism |
| 2019 | N | Y | 359126 | MBOVPG45_RS01595 | 1467 | 1 | 0.07 | N | MFS transporter | Transport protein |
| 2019 | N | Y | 364058 | MBOVPG45_RS01610 | 1203 | 6 | 0.50 | N | acetate/propionate family kinase | Metabolism |
| 2019 | N | Y | 365312 | MBOVPG45_RS01615 | 975 | 8 | 0.82 | N | phosphate acetyltransferase | Metabolism |
| 2019 | N | Y | 369370 | MBOVPG45_RS01630 | 972 | 3 | 0.31 | N | lactate dehydrogenase | Metabolism |
| 2019 | N | Y | 370053 | MBOVPG45_RS01635 | 1245 | 4 | 0.32 | N | malate transporter | Metabolism |
| 2019 | N | Y | 373493 | MBOVPG45_RS01645 | 1572 | 7 | 0.45 | N | glycosyltransferase family 2 protein | Metabolism |

| | | | | | | | | | | | |
|---|---|---|---|---|---|---|---|---|---|---|---|
| 2019 | N | Y | 374344 | MBOVPG45_RS01650 | 1434 | 6 | 0.42 | N | transposase | Transposable elements | |
| 2019 | N | Y | 376115 | MBOVPG45_RS01655 | 795 | 5 | 0.63 | 2019 | alpha/beta hydrolase | Multi-role family | |
| 2019 | N | Y | 377305 | MBOVPG45_RS01660 | 732 | 1 | 0.14 | N | nitroreductase | Metabolism | |
| 2019 | N | Y | 377784 | MBOVPG45_RS01665 | 1452 | 18 | 1.24 | N | phosphotransferase | Metabolism | |
| 2019 | N | Y | 381462 | MBOVPG45_RS01675 | 759 | 2 | 0.26 | 2019 | phosphotransferase | Metabolism | |
| 2019 | N | Y | 397770 | MBOVPG45_RS01750 | 1572 | 6 | 0.38 | N | phospho-sugar mutase | Metabolism | |
| 2019 | N | Y | 404108 | MBOVPG45_RS01780 | 2733 | 15 | 0.55 | N | cation-transporting P-type ATPase | Transport protein | |
| 2019 | N | Y | 410113 | MBOVPG45_RS01815 | 1029 | 1 | 0.10 | N | IS30-like element ISMbov1 family transposase | Transposable elements | |
| 2019 | N | Y | 417290 | MBOVPG45_RS01845 | 1188 | 4 | 0.34 | N | phosphopentomutase | Metabolism | |
| Both | Y | Y | 426836 | MBOVPG45_RS01870 | 2232 | 7 | 0.31 | N | putative immunoglobulin-blocking virulence protein | Virulence | |
| 2019 | N | Y | 430161 | MBOVPG45_RS01875 | 2250 | 1 | 0.04 | N | putative immunoglobulin-blocking virulence protein | Virulence | |
| Both | Y | Y | 449716 | MBOVPG45_RS01940 | 2205 | 4 | 0.18 | 2019 | UvrD-helicase domain-containing protein | DNA repair | Mutation increases error rate, increases chance for AMR mutation |
| 2019 | N | Y | 466370 | msrB | 930 | 2 | 0.22 | N | peptide-methionine (R)-S-oxide reductase MsrB | Metabolism | |
| Both | Y | Y | 466982 | MBOVPG45_RS02020 | 276 | 4 | 1.45 | 1982 | Smr/MutS family protein | DNA repair, DNA recombination | Mutation increases error rate, increases chance for AMR mutation |
| 2019 | N | Y | 467471 | mutM | 840 | 4 | 0.48 | N | bifunctional DNA-formamidopyrimidine glycosylase/DNA-(apurinic or apyrimidinic site) lyase | DNA repair | Mutation increases error rate, increases chance for AMR mutation |
| 2019 | N | Y | 468171 | MBOVPG45_RS02030 | 1293 | 4 | 0.31 | N | glucose-6-phosphate isomerase | Metabolism | |
| 2019 | N | Y | 469380 | MBOVPG45_RS02035 | 1251 | 11 | 0.88 | N | glucose-6-phosphate isomerase | Metabolism | |
| 2019 | N | Y | 471148 | MBOVPG45_RS02045 | 1365 | 4 | 0.29 | N | phosphopyruvate hydratase (eno) | Metabolism | |
| 2019 | N | Y | 473268 | MBOVPG45_RS02050 | 1434 | 5 | 0.35 | N | transposase | Transposable elements | |
| 2019 | N | Y | 476136 | MBOVPG45_RS02060 | 1017 | 1 | 0.10 | N | IS30 family transposase | Transposable elements | |
| 2019 | N | Y | 488484 | MBOVPG45_RS02110 | 1302 | 1 | 0.08 | N | ATP-binding protein | Multi-role family | |
| 2019 | N | Y | 496543 | MBOVPG45_RS02130 | 945 | 8 | 0.85 | 2019 | variable surface lipoprotein | variable surface lipoprotein | |
| 2019 | N | Y | 497823 | MBOVPG45_RS02135 | 552 | 5 | 0.91 | N | variable surface lipoprotein | variable surface lipoprotein | |
| 2019 | N | Y | 498724 | MBOVPG45_RS02140 | 1017 | 1 | 0.10 | N | IS30 family transposase | Transposable elements | |
| 2019 | N | Y | 501954 | MBOVPG45_RS02155 | 1434 | 3 | 0.21 | N | transposase | Transposable elements | |
| 2019 | N | Y | 507923 | MBOVPG45_RS02180 | 702 | 14 | 1.99 | N | variable surface lipoprotein | variable surface lipoprotein | |
| 2019 | N | Y | 512844 | atpH | 561 | 4 | 0.71 | N | ATP synthase F1 subunit delta | ATP synthesis | |
| 2019 | N | Y | 513447 | MBOVPG45_RS02215 | 1587 | 1 | 0.06 | N | F0F1 ATP synthase subunit alpha | ATP synthesis | |
| 1982 | Y | N | 515730 | MBOVPG45_RS02220 | 864 | 1 | 0.12 | N | F0F1 ATP synthase subunit gamma | ATP synthesis | |
| 2019 | N | Y | 517631 | MBOVPG45_RS02230 | 414 | 1 | 0.24 | N | ATP synthase F1 subunit epsilon | ATP synthesis | |
| 2019 | N | Y | 519897 | MBOVPG45_RS02245 | 717 | 1 | 0.14 | N | glycerol-3-phosphate acyltransferase | Metabolism | |
| Both | Y | Y | 521292 | lon | 3132 | 5 | 0.16 | N | endopeptidase La | Metabolism | |
| 2019 | N | Y | 528299 | MBOVPG45_RS02275 | 1947 | 9 | 0.46 | N | M13 family metallopeptidase | Metabolism | |
| 2019 | N | Y | 530918 | MBOVPG45_RS02285 | 378 | 5 | 1.32 | N | glycine cleavage system protein H | Metabolism | |
| 2019 | N | Y | 535888 | MBOVPG45_RS02310 | 669 | 1 | 0.15 | N | TrkA family potassium uptake protein | Metabolism | |
| 2019 | N | Y | 536751 | MBOVPG45_RS02315 | 1560 | 2 | 0.13 | N | TrkH family potassium uptake protein | Metabolism | |
| 2019 | N | Y | 540693 | uvrB | 2010 | 9 | 0.45 | N | excinuclease ABC subunit UvrB | DNA repair | Mutation increases error rate, increases chance for AMR mutation |
| 2019 | N | Y | 542155 | uvrA | 2832 | 10 | 0.35 | N | excinuclease ABC subunit UvrA | DNA repair | Mutation increases error rate, increases chance for AMR mutation |
| Both | Y | Y | 557979 | MBOVPG45_RS02400 | 2826 | 3 | 0.11 | N | conjugal transfer protein TraE | Transformation | |
| 2019 | N | Y | 569339 | MBOVPG45_RS02455 | 1773 | 5 | 0.28 | N | type IV secretory system conjugative DNA transfer family protein | Transformation | |
| Both | Y | Y | 575129 | MBOVPG45_RS02480 | 1602 | 4 | 0.25 | 1982 | IS1634-like element ISMbov3 family transposase | Transposable elements | |
| 2019 | N | Y | 577264 | MBOVPG45_RS02490 | 453 | 4 | 0.88 | 2019 | SocA family protein | Transport protein | |
| 2019 | N | Y | 578011 | MBOVPG45_RS02495 | 723 | 1 | 0.14 | N | SDR family oxidoreductase | Metabolism | |
| 2019 | N | Y | 582294 | ruvX | 441 | 1 | 0.23 | N | Holliday junction resolvase RuvX | Recombination | |
| 2019 | N | Y | 586464 | MBOVPG45_RS02525 | 729 | 3 | 0.41 | N | purine-nucleoside phosphorylase | Metabolism | |
| 2019 | N | Y | 591273 | MBOVPG45_RS02550 | 1050 | 6 | 0.57 | N | zinc-dependent alcohol dehydrogenase | Metabolism | |
| 2019 | N | Y | 592432 | MBOVPG45_RS02555 | 585 | 10 | 1.71 | 2019 | variable surface lipoprotein | variable surface lipoprotein | |
| 2019 | N | Y | 593925 | MBOVPG45_RS02560 | 1041 | 1 | 0.10 | N | zinc-dependent alcohol dehydrogenase | Metabolism | |
| 2019 | N | Y | 594709 | MBOVPG45_RS02565 | 732 | 3 | 0.41 | N | DNA processing protein DprA | Transformation | |
| 2019 | N | Y | 595603 | MBOVPG45_RS02570 | 837 | 4 | 0.48 | N | HAD family phosphatase | Metabolism | |
| 2019 | N | Y | 596888 | MBOVPG45_RS02575 | 1353 | 3 | 0.22 | N | YitT family protein | Stress protein | |
| 2019 | N | Y | 597675 | MBOVPG45_RS02580 | 2979 | 5 | 0.17 | N | AAA family ATPase | Multi-role family | |
| 2019 | N | Y | 609179 | glpK | 1509 | 3 | 0.20 | N | glycerol kinase GlpK | Metabolism | |
| 2019 | N | Y | 615303 | MBOVPG45_RS02645 | 1671 | 2 | 0.12 | N | APC family permease | Transport protein | |
| 2019 | N | Y | 620753 | MBOVPG45_RS02665 | 1245 | 2 | 0.16 | N | IS3 family transposase | Transposable elements | |
| 2019 | N | Y | 633285 | MBOVPG45_RS02720 | 1356 | 1 | 0.07 | N | lipoprotein ABC transporter substrate-binding protein | Metabolism | |
| 2019 | N | Y | 641833 | MBOVPG45_RS02750 | 1365 | 3 | 0.22 | N | FAD-dependent oxidoreductase | Metabolism | |
| 2019 | N | Y | 647839 | MBOVPG45_RS02775 | 927 | 1 | 0.11 | N | type II restriction endonuclease | Restriction enzyme | |
| 2019 | N | Y | 648784 | MBOVPG45_RS02780 | 849 | 13 | 1.53 | N | DNA adenine methylase | DNA repair | |
| Both | Y | Y | 650147 | MBOVPG45_RS02785 | 1374 | 24 | 1.75 | N | variable surface lipoprotein | variable surface lipoprotein | |
| 2019 | N | Y | 659437 | rmuC | 1440 | 1 | 0.07 | N | DNA recombination protein RmuC | Recombination | |
| 2019 | N | Y | 659637 | MBOVPG45_RS02815 | 1611 | 2 | 0.12 | N | dicarboxylate/amino acid:cation symporter | Metabolism | |
| 1982 | Y | N | 662114 | MBOVPG45_RS02820 | 1350 | 1 | 0.07 | N | FAD-dependent oxidoreductase | Metabolism | |
| 2019 | N | Y | 663026 | MBOVPG45_RS02825 | 1245 | 2 | 0.16 | N | IS3 family transposase | Transposable elements | |
| Both | Y | Y | 666001 | MBOVPG45_RS02835 | 900 | 4 | 0.44 | Both | variable surface lipoprotein | variable surface lipoprotein | |
| 2019 | N | Y | 670440 | MBOVPG45_RS02860 | 1434 | 5 | 0.35 | N | transposase | Transposable elements | |
| 2019 | N | Y | 677174 | MBOVPG45_RS02885 | 2001 | 4 | 0.20 | N | DHH family phosphoesterase | DNA repair, DNA recombination | |
| 2019 | N | Y | 677990 | MBOVPG45_RS02890 | 1068 | 14 | 1.31 | N | variable surface lipoprotein | variable surface lipoprotein | |
| 2019 | N | Y | 679145 | MBOVPG45_RS02895 | 1602 | 3 | 0.19 | N | IS1634 family transposase | Transposable elements | |
| 2019 | N | Y | 693602 | ruvA | 597 | 2 | 0.34 | N | Holliday junction branch migration protein RuvA | Recombination | |
| 2019 | N | Y | 697337 | MBOVPG45_RS02990 | 711 | 4 | 0.56 | N | signal peptidase II | Cell signalling | |
| 2019 | N | Y | 704332 | MBOVPG45_RS03050 | 1245 | 2 | 0.16 | N | IS3 family transposase | Transposable elements | |
| 2019 | N | Y | 713717 | MBOVPG45_RS03080 | 2271 | 10 | 0.44 | N | S8 family peptidase | Peptidase | |
| 2019 | N | Y | 716234 | MBOVPG45_RS03085 | 1044 | 1 | 0.10 | N | AAA family ATPase | Multi-role family | |

| | | | | | | | | | | |
|---|---|---|---|---|---|---|---|---|---|---|
| 2019 | N | Y | 717489 | MBOVPG45_RS03090 | 1434 | 1 | 0.07 | N | transposase | Transposable elements |
| 2019 | N | Y | 719249 | MBOVPG45_RS03095 | 2226 | 7 | 0.31 | N | AAA family ATPase | Multi-role family |
| Both | Y | Y | 733123 | ftsY | 1059 | 2 | 0.19 | N | signal recognition particle-docking protein FtsY | Cell signalling |
| 2019 | N | Y | 736512 | MBOVPG45_RS03185 | 336 | 2 | 0.60 | N | HIT family protein | Multi-role family |
| 2019 | N | Y | 741395 | MBOVPG45_RS03210 | 555 | 1 | 0.18 | N | inorganic diphosphatase | Metabolism |
| 2019 | N | Y | 743052 | MBOVPG45_RS03215 | 1947 | 4 | 0.21 | N | transketolase | Metabolism |
| 2019 | N | Y | 749051 | prs | 996 | 1 | 0.10 | N | ribose-phosphate pyrophosphokinase | Metabolism |
| 2019 | N | Y | 749061 | rsmG (pseudo), prs | 695 | 1 | 0.14 | N | pseudogene | None |
| 2019 | N | Y | 751100 | MBOVPG45_RS03260 | 1029 | 4 | 0.39 | N | IS30-like element ISMbov1 family transposase | Transposable elements |
| Both | Y | Y | 753684 | MBOVPG45_RS03275 | 717 | 5 | 0.70 | N | ribonuclease HIII | Rnase |
| 2019 | N | Y | 756247 | MBOVPG45_RS03285 | 1602 | 4 | 0.25 | N | IS1634-like element ISMbov3 family transposase | Transposable elements |
| 2019 | N | Y | 762108 | MBOVPG45_RS03300 | 1398 | 3 | 0.21 | N | CvpA family protein | Toxin biosynthesis |
| Both | Y | Y | 772736 | MBOVPG45_RS03345 | 2316 | 25 | 1.08 | 1982 | S8 family serine peptidase | Peptidase |
| 2019 | N | Y | 778219 | MBOVPG45_RS03365 | 1812 | 25 | 1.38 | 2019 | LppA family lipoprotein | Metabolism |
| 2019 | N | Y | 780828 | MBOVPG45_RS03375 | 2310 | 9 | 0.39 | 2019 | S8 family serine peptidase | Peptidase |
| 2019 | N | Y | 788183 | MBOVPG45_RS03390 | 918 | 4 | 0.44 | N | 5'-3' exonuclease | DNA repair |
| 2019 | N | Y | 789498 | MBOVPG45_RS03395 | 1602 | 2 | 0.12 | N | IS1634 family transposase | Transposable elements |
| 2019 | N | Y | 791109 | MBOVPG45_RS03400 | 2043 | 11 | 0.54 | N | bifunctional metallophosphatase/5'-nucleotidase | DNA repair |
| 2019 | N | Y | 802899 | MBOVPG45_RS03450 | 1389 | 3 | 0.22 | N | transposase | Transposable elements |
| 2019 | N | Y | 810822 | MBOVPG45_RS03485 | 1089 | 10 | 0.92 | N | M42 family peptidase | Metabolism |
| 2019 | N | Y | 812080 | MBOVPG45_RS03490 | 1017 | 1 | 0.10 | N | IS30 family transposase | Transposable elements |
| 2019 | N | Y | 839138 | MBOVPG45_RS03550 | 2169 | 5 | 0.23 | N | AAA domain-containing protein | Multi-role family |
| 1982 | Y | N | 843830 | MBOVPG45_RS03560 | 1098 | 1 | 0.09 | N | DNA adenine methylase | DNA repair |
| 2019 | N | Y | 848393 | MBOVPG45_RS03590 | 2973 | 28 | 0.94 | N | phosphoglucomutase | Metabolism |
| 2019 | N | Y | 851558 | ptsP | 1713 | 7 | 0.41 | N | phosphoenolpyruvate--protein phosphotransferase | Metabolism |
| 2019 | N | Y | 854531 | araD | 732 | 1 | 0.14 | N | L-ribulose-5-phosphate 4-epimerase | Metabolism |
| 2019 | N | Y | 855274 | MBOVPG45_RS03615 | 885 | 4 | 0.45 | N | L-ribulose-5-phosphate 3-epimerase | Metabolism |
| 2019 | N | Y | 856260 | MBOVPG45_RS03620 | 654 | 3 | 0.46 | N | 3-keto-L-gulonate-6-phosphate decarboxylase UlaD | Metabolism |
| 2019 | N | Y | 856957 | MBOVPG45_RS03625 | 489 | 1 | 0.20 | N | PTS transporter subunit EIIA | Transport protein |
| 2019 | N | Y | 858957 | MBOVPG45_RS03635 | 1809 | 1 | 0.06 | N | PTS ascorbate transporter subunit IIC | Transport protein |
| 2019 | N | Y | 860078 | MBOVPG45_RS03640 | 1062 | 1 | 0.09 | N | phosphotriesterase | Metabolism |
| 2019 | N | Y | 862854 | MBOVPG45_RS04565 | 168 | 5 | 2.98 | 2019 | variable surface lipoprotein | variable surface lipoprotein |
| Both | Y | Y | 864315 | MBOVPG45_RS03660 | 207 | 3 | 1.45 | 1982 | variable surface lipoprotein | variable surface lipoprotein |
| 2019 | N | Y | 864948 | MBOVPG45_RS03665 | 900 | 6 | 0.67 | N | Fic family protein | Posttranslational modification |
| 2019 | N | Y | 866317 | MBOVPG45_RS04570 | 903 | 1 | 0.11 | N | site-specific DNA-methyltransferase | Methylation |
| 2019 | N | Y | 867221 | MBOVPG45_RS04575 | 522 | 5 | 0.96 | 2019 | site-specific DNA-methyltransferase | Methylation |
| 2019 | N | Y | 870763 | MBOVPG45_RS03695 | 1017 | 1 | 0.10 | N | IS30 family transposase | Transposable elements |
| 2019 | N | Y | 876432 | MBOVPG45_RS03720 | 2757 | 25 | 0.91 | N | C1 family peptidase | Metabolism |
| 2019 | N | Y | 879655 | MBOVPG45_RS03725 | 1008 | 2 | 0.20 | N | ZIP family metal transporter | Metabolism |
| 2019 | N | Y | 880921 | nadE | 813 | 3 | 0.37 | N | NAD(+) synthase | Metabolism |
| 2019 | N | Y | 882769 | MBOVPG45_RS03745 | 924 | 4 | 0.43 | N | alpha/beta hydrolase | Multi-role family |
| 1982 | Y | N | 885156 | MBOVPG45_RS03755 | 900 | 1 | 0.11 | N | ATP-binding protein | Multi-role family |
| 2019 | N | Y | 886043 | MBOVPG45_RS03760 | 1008 | 4 | 0.40 | N | DNA damage-inducible protein DnaD | DNA repair |
| Both | Y | Y | 887092 | MBOVPG45_RS03765 | 573 | 4 | 0.70 | N | dephospho-CoA kinase | Metabolism |
| 2019 | N | Y | 890617 | MBOVPG45_RS03795 | 1125 | 11 | 0.98 | 2019 | site-specific DNA-methyltransferase | Methylation |
| 2019 | N | Y | 891887 | MBOVPG45_RS03800 | 1284 | 8 | 0.62 | N | MFS transporter | Transport protein |
| 2019 | N | Y | 899288 | MBOVPG45_RS03835 | 654 | 1 | 0.15 | N | chromate transporter | Metabolism |
| 2019 | N | Y | 906851 | recR | 588 | 3 | 0.51 | N | recombination protein RecR | Recombination |
| Both | Y | Y | 912118 | MBOVPG45_RS03905 | 1497 | 10 | 0.67 | N | 2,3-bisphosphoglycerate-independent phosphoglycerate mutase | Metabolism |
| 2019 | N | Y | 916526 | pgsA | 609 | 2 | 0.33 | 2019 | CDP-diacylglycerol--glycerol-3-phosphate 3-phosphatidyltransferase | Metabolism |
| 2019 | N | Y | 917577 | MBOVPG45_RS03935 | 906 | 2 | 0.22 | N | phosphatidate cytidylyltransferase | Metabolism |
| 2019 | N | Y | 928069 | MBOVPG45_RS03990 | 1029 | 1 | 0.10 | N | IS30 family transposase | Transposable elements |
| Both | Y | Y | 930394 | MBOVPG45_RS04005 | 1092 | 2 | 0.18 | N | variable surface lipoprotein | variable surface lipoprotein |
| Both | Y | Y | 931489 | MBOVPG45_RS04010 | 765 | 17 | 2.22 | 1982 | variable surface lipoprotein | variable surface lipoprotein |
| 2019 | N | Y | 939874 | MBOVPG45_RS04050 | 615 | 2 | 0.33 | N | abortive infection protein | Immune response |
| 2019 | N | Y | 940364 | MBOVPG45_RS04055 | 861 | 4 | 0.46 | N | nucleotidyl transferase AbiEii/AbiGii toxin family protein | Toxin biosynthesis |
| 2019 | N | Y | 942397 | MBOVPG45_RS04065 | 1017 | 1 | 0.10 | N | IS30 family transposase | Transposable elements |
| Both | Y | Y | 945959 | MBOVPG45_RS04085 | 759 | 4 | 0.53 | 1982 | variable surface lipoprotein | variable surface lipoprotein |
| Both | Y | Y | 946967 | MBOVPG45_RS04090 | 609 | 7 | 1.15 | N | variable surface lipoprotein | variable surface lipoprotein |
| 2019 | N | Y | 949632 | MBOVPG45_RS04110 | 1845 | 1 | 0.05 | N | ribonuclease J | Rnase |
| 2019 | N | Y | 959225 | MBOVPG45_RS04145 | 855 | 8 | 0.94 | N | RluA family pseudouridine synthase | Posttranslational modification |
| Both | Y | Y | 967387 | MBOVPG45_RS04200 | 1602 | 6 | 0.37 | 1982 | IS1634-like element ISMbov3 family transposase | Transposable elements |
| 2019 | N | Y | 972044 | MBOVPG45_RS04230 | 234 | 1 | 0.43 | N | acyl carrier protein | Metabolism |
| 2019 | N | Y | 976131 | MBOVPG45_RS04250 | 1422 | 5 | 0.35 | N | YitT family protein | Stress protein |
| 2019 | N | Y | 978606 | MBOVPG45_RS04260 | 315 | 2 | 0.63 | N | thioredoxin family protein | Metabolism |
| 2019 | N | Y | 981821 | MBOVPG45_RS04280 | 207 | 1 | 0.48 | 2019 | variable surface lipoprotein | variable surface lipoprotein |
| Both | Y | Y | 984371 | MBOVPG45_RS04300 | 1245 | 2 | 0.16 | N | IS3 family transposase | Transposable elements |
| Both | Y | Y | 985381 | mgtA | 2709 | 9 | 0.33 | N | magnesium-translocating P-type ATPase | Metabolism |
| Both | Y | Y | 997722 | hflB | 2031 | 3 | 0.15 | N | ATP-dependent zinc metalloprotease FtsH | Membrane maintenance |
| 2019 | N | Y | 1001054 | MBOVPG45_RS04335 | 2271 | 9 | 0.40 | N | AAA family ATPase | Multi-role family |

### S2-3. Uncharacterized Proteins:

| Strain | 1982 (Y/N) | 2019 (Y/N) | Ref Pos | Gene Name | PG45 Gene size (nt) | #Mutations | Mutations/100 nt | Disrupted (Y/N) | Gene Description | |
|---|---|---|---|---|---|---|---|---|---|---|
| Both | Y | Y | 41706 | MBOVPG45_RS00190 | 9981 | 5 | 0.05 | 2019 | PDxFFG protein | Uncharacterized protein |
| 2019 | N | Y | 60006 | MBOVPG45_RS00235 | 1035 | 1 | 0.10 | N | membrane protein | Uncharacterized protein |
| 2019 | N | Y | 117143 | MBOVPG45_RS00540 | 546 | 3 | 0.55 | N | DJ-1/PfpI family protein | Uncharacterized protein |
| Both | Y | Y | 141369 | MBOVPG45_RS00640 | 1818 | 2 | 0.11 | N | membrane protein | Uncharacterized protein |
| 2019 | N | Y | 154022 | MBOVPG45_RS00690 | 1629 | 12 | 0.74 | 2019 | membrane protein | Uncharacterized protein |
| Both | Y | Y | 158505 | MBOVPG45_RS00710 | 741 | 24 | 3.24 | N | DUF285 domain-containing protein | Uncharacterized protein |

| | | | | | | | | | | |
|---|---|---|---|---|---|---|---|---|---|---|
| Both | Y | Y | 162561 | MBOVPG45_RS00725 | 1521 | 17 | 1.12 | Both | DUF285 domain-containing protein | Uncharacterized protein |
| Both | Y | Y | 164311 | MBOVPG45_RS00730 | 1521 | 16 | 1.05 | 2019 | DUF285 domain-containing protein | Uncharacterized protein |
| 2019 | N | Y | 182781 | MBOVPG45_RS00795 | 384 | 3 | 0.78 | N | membrane protein | Uncharacterized protein |
| 2019 | N | Y | 283862 | MBOVPG45_RS01215 | 4713 | 26 | 0.55 | N | DUF4011 domain-containing protein | Uncharacterized protein |
| 2019 | N | Y | 313704 | MBOVPG45_RS01400 | 1056 | 13 | 1.23 | 2019 | Cof-type HAD-IIB family hydrolase | Uncharacterized protein |
| 2019 | N | Y | 339998 | MBOVPG45_RS01515 | 825 | 2 | 0.24 | N | Cof-type HAD-IIB family hydrolase | Uncharacterized protein |
| 2019 | N | Y | 356763 | MBOVPG45_RS01580 | 1707 | 1 | 0.06 | N | membrane protein | Uncharacterized protein |
| 2019 | N | Y | 414758 | MBOVPG45_RS01835 | 1185 | 24 | 2.03 | N | DUF285 domain-containing protein | Uncharacterized protein |
| 2019 | N | Y | 455097 | MBOVPG45_RS01960 | 762 | 1 | 0.13 | N | membrane protein | Uncharacterized protein |
| Both | Y | Y | 493605 | MBOVPG45_RS02125 | 1989 | 11 | 0.55 | 1982 | BspA family leucine-rich repeat surface protein | Uncharacterized protein |
| 2019 | N | Y | 545712 | MBOVPG45_RS02350 | 810 | 5 | 0.62 | N | DUF45 domain-containing protein | Uncharacterized protein |
| 2019 | N | Y | 561474 | MBOVPG45_RS02405 | 2238 | 5 | 0.22 | N | membrane protein | Uncharacterized protein |
| 2019 | N | Y | 573609 | MBOVPG45_RS02470 | 1119 | 10 | 0.89 | N | BspA family leucine-rich repeat surface protein | Uncharacterized protein |
| 2019 | N | Y | 587200 | MBOVPG45_RS02530 | 1524 | 10 | 0.66 | N | DUF285 domain-containing protein | Uncharacterized protein |
| 2019 | N | Y | 641166 | MBOVPG45_RS02745 | 786 | 4 | 0.51 | N | Nif3-like dinuclear metal center hexameric protein | Uncharacterized protein |
| 2019 | N | Y | 651781 | MBOVPG45_RS02795 | 2310 | 9 | 0.39 | 2019 | leucine-rich repeat domain-containing protein | Uncharacterized protein |
| 2019 | N | Y | 654099 | MBOVPG45_RS02800 | 1839 | 4 | 0.22 | 2019 | leucine-rich repeat domain-containing protein | Uncharacterized protein |
| 1982 | Y | N | 667694 | MBOVPG45_RS02845 | 909 | 1 | 0.11 | 1982 | BspA family leucine-rich repeat surface protein | Uncharacterized protein |
| 1982 | Y | N | 690881 | MBOVPG45_RS02945 | 2367 | 1 | 0.04 | N | membrane protein | Uncharacterized protein |
| 2019 | N | Y | 754547 | MBOVPG45_RS03280 | 930 | 2 | 0.22 | N | Cof-type HAD-IIB family hydrolase | Uncharacterized protein |
| Both | Y | Y | 814940 | MBOVPG45_RS03500 | 8013 | 58 | 0.72 | 2019 | SGNH/GDSL hydrolase family protein | Uncharacterized protein |
| 2019 | N | Y | 833816 | MBOVPG45_RS03530 | 726 | 5 | 0.69 | N | leucine-rich repeat domain-containing protein | Uncharacterized protein |
| 2019 | N | Y | 860963 | MBOVPG45_RS03645 | 858 | 6 | 0.70 | N | Cof-type HAD-IIB family hydrolase | Uncharacterized protein |
| 2019 | N | Y | 863158 | MBOVPG45_RS03655 | 999 | 35 | 3.50 | N | DUF285 domain-containing protein | Uncharacterized protein |
| 2019 | N | Y | 887599 | MBOVPG45_RS03770 | 381 | 5 | 1.31 | N | YigZ family protein | Uncharacterized protein |
| 2019 | N | Y | 896766 | MBOVPG45_RS03820 | 1059 | 1 | 0.09 | N | membrane protein | Uncharacterized protein |
| 2019 | N | Y | 901081 | MBOVPG45_RS03850 | 612 | 2 | 0.33 | N | DUF402 domain-containing protein | Uncharacterized protein |
| 2019 | N | Y | 913679 | MBOVPG45_RS03910 | 900 | 12 | 1.33 | N | Cof-type HAD-IIB family hydrolase | Uncharacterized protein |
| 2019 | N | Y | 920884 | MBOVPG45_RS03955 | 303 | 3 | 0.99 | N | YlxR family protein | Uncharacterized protein |
| 2019 | N | Y | 929129 | MBOVPG45_RS03995 | 1008 | 2 | 0.20 | N | membrane protein | Uncharacterized protein |
| 2019 | N | Y | 954349 | MBOVPG45_RS04120 | 2418 | 10 | 0.41 | N | DUF2779 domain-containing protein | Uncharacterized protein |
| Both | Y | Y | 980663 | MBOVPG45_RS04275 | 999 | 48 | 4.80 | 2019 | DUF285 domain-containing protein | Uncharacterized protein |

### S2-4. Hypothetical Proteins

| Strain | 1982 (Y/N) | 2019 (Y/N) | Ref Pos | Gene Name | PG45 Gene size (nt) | #Mutations | Mutations/100 nt | Disrupted (Y/N) | Gene Description | |
|---|---|---|---|---|---|---|---|---|---|---|
| 2019 | N | Y | 14976 | MBOVPG45_RS00085 | 453 | 1 | 0.22 | N | hypothetical protein | hypothetical protein |
| 2019 | N | Y | 30658 | MBOVPG45_RS00155 | 1422 | 1 | 0.07 | N | hypothetical protein | hypothetical protein |
| 2019 | N | Y | 65553 | MBOVPG45_RS00270 | 513 | 2 | 0.39 | N | hypothetical protein | hypothetical protein |
| 2019 | N | Y | 79432 | MBOVPG45_RS00340 | 765 | 1 | 0.13 | N | hypothetical protein | hypothetical protein |
| 2019 | N | Y | 97890 | MBOVPG45_RS00420 | 1194 | 13 | 1.09 | N | hypothetical protein | hypothetical protein |
| 2019 | N | Y | 99221 | MBOVPG45_RS00425 | 261 | 2 | 0.77 | N | hypothetical protein | hypothetical protein |
| 2019 | N | Y | 102344 | MBOVPG45_RS00450 | 459 | 3 | 0.65 | N | hypothetical protein | hypothetical protein |
| 2019 | N | Y | 104072 | MBOVPG45_RS00460 | 762 | 16 | 2.10 | N | hypothetical protein | hypothetical protein |
| 2019 | N | Y | 106274 | MBOVPG45_RS00475 | 234 | 3 | 1.28 | N | hypothetical protein | hypothetical protein |
| Both | Y | Y | 107419 | MBOVPG45_RS00485 | 390 | 5 | 1.28 | N | hypothetical protein | hypothetical protein |
| 2019 | N | Y | 110917 | MBOVPG45_RS00505 | 690 | 10 | 1.45 | N | hypothetical protein | hypothetical protein |
| 2019 | N | Y | 118227 | MBOVPG45_RS00545 | 909 | 1 | 0.11 | N | hypothetical protein | hypothetical protein |
| Both | Y | Y | 119112 | MBOVPG45_RS00550 | 2886 | 48 | 1.66 | N | hypothetical protein | hypothetical protein |
| Both | Y | Y | 128082 | MBOVPG45_RS00575 | 984 | 2 | 0.20 | N | hypothetical protein | hypothetical protein |
| 2019 | N | Y | 132243 | MBOVPG45_RS00590 | 1941 | 2 | 0.10 | N | hypothetical protein | hypothetical protein |
| 1982 | Y | N | 156567 | MBOVPG45_RS00700 | 279 | 1 | 0.36 | N | hypothetical protein | hypothetical protein |
| 2019 | N | Y | 165866 | MBOVPG45_RS00735 | 2436 | 6 | 0.25 | N | hypothetical protein | hypothetical protein |
| 2019 | N | Y | 175241 | MBOVPG45_RS00770 | 1503 | 22 | 1.46 | N | hypothetical protein | hypothetical protein |
| 2019 | N | Y | 178282 | MBOVPG45_RS00780 | 981 | 19 | 1.94 | N | hypothetical protein | hypothetical protein |
| 2019 | N | Y | 185839 | MBOVPG45_RS04405 | 234 | 5 | 2.14 | N | hypothetical protein | hypothetical protein |
| 2019 | N | Y | 200561 | MBOVPG45_RS00865 | 1410 | 2 | 0.14 | N | hypothetical protein | hypothetical protein |
| 1982 | Y | N | 235507 | MBOVPG45_RS01005 | 990 | 2 | 0.20 | 1982 | hypothetical protein | hypothetical protein |
| 2019 | N | Y | 239758 | MBOVPG45_RS01020 | 306 | 1 | 0.33 | N | hypothetical protein | hypothetical protein |
| 2019 | N | Y | 241469 | MBOVPG45_RS01030 | 1521 | 2 | 0.13 | 2019 | hypothetical protein | hypothetical protein |
| 2019 | N | Y | 242137 | MBOVPG45_RS01035 | 660 | 9 | 1.36 | N | hypothetical protein | hypothetical protein |
| Both | Y | Y | 242831 | MBOVPG45_RS01040 | 636 | 7 | 1.10 | 1982 | hypothetical protein | hypothetical protein |
| 2019 | N | Y | 253208 | MBOVPG45_RS01090 | 897 | 1 | 0.11 | N | hypothetical protein | hypothetical protein |
| 2019 | N | Y | 271059 | MBOVPG45_RS01175 | 480 | 21 | 4.38 | 2019 | hypothetical protein | hypothetical protein |
| 2019 | N | Y | 291937 | MBOVPG45_RS01225 | 1503 | 7 | 0.47 | N | hypothetical protein | hypothetical protein |
| 2019 | N | Y | 334428 | MBOVPG45_RS01490 | 1053 | 18 | 1.71 | N | hypothetical protein | hypothetical protein |
| 2019 | N | Y | 341776 | MBOVPG45_RS01525 | 1539 | 9 | 0.58 | N | hypothetical protein | hypothetical protein |
| 2019 | N | Y | 347859 | MBOVPG45_RS01550 | 1170 | 6 | 0.51 | N | hypothetical protein | hypothetical protein |
| 2019 | N | Y | 352974 | MBOVPG45_RS01565 | 405 | 2 | 0.49 | N | hypothetical protein | hypothetical protein |
| 2019 | N | Y | 361083 | MBOVPG45_RS01600 | 2238 | 28 | 1.25 | N | hypothetical protein | hypothetical protein |
| 2019 | N | Y | 363231 | MBOVPG45_RS01605 | 906 | 5 | 0.55 | N | hypothetical protein | hypothetical protein |
| 2019 | N | Y | 398393 | MBOVPG45_RS01755 | 888 | 27 | 3.04 | N | hypothetical protein | hypothetical protein |
| 2019 | N | Y | 399190 | MBOVPG45_RS01760 | 891 | 22 | 2.47 | N | hypothetical protein | hypothetical protein |
| Both | Y | Y | 400121 | MBOVPG45_RS01765 | 1149 | 35 | 3.05 | 2019 | hypothetical protein | hypothetical protein |
| 2019 | N | Y | 418892 | MBOVPG45_RS01850 | 1248 | 3 | 0.24 | N | hypothetical protein | hypothetical protein |
| 2019 | N | Y | 424904 | MBOVPG45_RS01865 | 1171 | 34 | 2.90 | N | hypothetical protein | hypothetical protein |
| 2019 | N | Y | 433117 | MBOVPG45_RS01880 | 2571 | 1 | 0.04 | N | hypothetical protein | hypothetical protein |
| 2019 | N | Y | 434748 | MBOVPG45_RS01885 | 1512 | 1 | 0.07 | N | hypothetical protein | hypothetical protein |
| 2019 | N | Y | 437928 | MBOVPG45_RS01905 | 2244 | 3 | 0.13 | N | hypothetical protein | hypothetical protein |
| 2019 | N | Y | 443328 | MBOVPG45_RS01920 | 1071 | 2 | 0.19 | 2019 | hypothetical protein | hypothetical protein |
| 2019 | N | Y | 444935 | MBOVPG45_RS01925 | 2385 | 1 | 0.04 | N | hypothetical protein | hypothetical protein |
| 1982 | Y | N | 447801 | MBOVPG45_RS01935 | 1647 | 1 | 0.06 | N | hypothetical protein | hypothetical protein |
| 2019 | N | Y | 451398 | MBOVPG45_RS01945 | 669 | 2 | 0.30 | N | hypothetical protein | hypothetical protein |
| 2019 | N | Y | 456260 | MBOVPG45_RS01970 | 579 | 1 | 0.17 | N | hypothetical protein | hypothetical protein |
| Both | Y | Y | 463976 | MBOVPG45_RS02010 | 1944 | 38 | 1.95 | 2019 | hypothetical protein | hypothetical protein |
| 2019 | N | Y | 479290 | MBOVPG45_RS02080 | 1872 | 21 | 1.12 | N | hypothetical protein | hypothetical protein |
| 2019 | N | Y | 491269 | MBOVPG45_RS02120 | 1725 | 17 | 0.99 | N | hypothetical protein | hypothetical protein |
| Both | Y | Y | 500449 | MBOVPG45_RS02150 | 687 | 18 | 2.62 | N | hypothetical protein | hypothetical protein |
| 2019 | N | Y | 503162 | MBOVPG45_RS02160 | 765 | 2 | 0.26 | N | hypothetical protein | hypothetical protein |
| 2019 | N | Y | 506905 | MBOVPG45_RS04420 | 441 | 4 | 0.91 | N | hypothetical protein | hypothetical protein |
| 2019 | N | Y | 508825 | MBOVPG45_RS02185 | 1674 | 21 | 1.25 | N | hypothetical protein | hypothetical protein |
| 2019 | N | Y | 510784 | MBOVPG45_RS02190 | 471 | 4 | 0.85 | 2019 | hypothetical protein | hypothetical protein |
| 2019 | N | Y | 526035 | MBOVPG45_RS02260 | 729 | 9 | 1.23 | N | hypothetical protein | hypothetical protein |

| | | | | | | | | | | |
|---|---|---|---|---|---|---|---|---|---|---|
| **Both** | Y | Y | 532709 | MBOVPG45_RS02295 | 1218 | 5 | 0.41 | N | hypothetical protein | hypothetical protein |
| 2019 | N | Y | 538024 | MBOVPG45_RS02320 | 453 | 1 | 0.22 | N | hypothetical protein | hypothetical protein |
| 2019 | N | Y | 551171 | MBOVPG45_RS02380 | 1233 | 1 | 0.08 | N | hypothetical protein | hypothetical protein |
| 2019 | N | Y | 552200 | MBOVPG45_RS02385 | 645 | 1 | 0.16 | N | hypothetical protein | hypothetical protein |
| 2019 | N | Y | 553973 | MBOVPG45_RS02390 | 4548 | 27 | 0.59 | N | hypothetical protein | hypothetical protein |
| 2019 | N | Y | 557345 | MBOVPG45_RS02395 | 588 | 4 | 0.68 | N | hypothetical protein | hypothetical protein |
| 2019 | N | Y | 563125 | MBOVPG45_RS02410 | 306 | 1 | 0.33 | N | hypothetical protein | hypothetical protein |
| 2019 | N | Y | 564824 | MBOVPG45_RS02420 | 1509 | 2 | 0.13 | 2019 | hypothetical protein | hypothetical protein |
| 2019 | N | Y | 565492 | MBOVPG45_RS02425 | 660 | 8 | 1.21 | N | hypothetical protein | hypothetical protein |
| 2019 | N | Y | 566186 | MBOVPG45_RS02430 | 663 | 5 | 0.75 | N | hypothetical protein | hypothetical protein |
| 2019 | N | Y | 567250 | MBOVPG45_RS02440 | 267 | 6 | 2.25 | 2019 | hypothetical protein | hypothetical protein |
| 2019 | N | Y | 567514 | MBOVPG45_RS02445 | 852 | 3 | 0.35 | N | hypothetical protein | hypothetical protein |
| 2019 | N | Y | 570980 | MBOVPG45_RS02460 | 1362 | 10 | 0.73 | N | hypothetical protein | hypothetical protein |
| 2019 | N | Y | 572379 | MBOVPG45_RS04635 | 153 | 6 | 3.92 | 2019 | hypothetical protein | hypothetical protein |
| 2019 | N | Y | 576731 | MBOVPG45_RS02485 | 390 | 3 | 0.77 | 2019 | hypothetical protein | hypothetical protein |
| **Both** | Y | Y | 578792 | MBOVPG45_RS02500 | 2172 | 37 | 1.70 | N | hypothetical protein | hypothetical protein |
| 2019 | N | Y | 581268 | MBOVPG45_RS02505 | 729 | 7 | 0.96 | N | hypothetical protein | hypothetical protein |
| 2019 | N | Y | 634259 | MBOVPG45_RS02725 | 1791 | 19 | 1.06 | N | hypothetical protein | hypothetical protein |
| 2019 | N | Y | 643726 | MBOVPG45_RS02755 | 504 | 1 | 0.20 | N | hypothetical protein | hypothetical protein |
| 2019 | N | Y | 651430 | MBOVPG45_RS02790 | 253 | 6 | 2.37 | 2019 | hypothetical protein | hypothetical protein |
| 2019 | N | Y | 669998 | MBOVPG45_RS02855 | 990 | 1 | 0.10 | N | hypothetical protein | hypothetical protein |
| 2019 | N | Y | 681045 | MBOVPG45_RS02900 | 1170 | 34 | 2.91 | 2019 | hypothetical protein | hypothetical protein |
| 2019 | N | Y | 732728 | MBOVPG45_RS03155 | 285 | 1 | 0.35 | N | hypothetical protein | hypothetical protein |
| 2019 | N | Y | 735556 | MBOVPG45_RS03180 | 699 | 11 | 1.57 | N | hypothetical protein | hypothetical protein |
| 2019 | N | Y | 736781 | MBOVPG45_RS03190 | 1386 | 22 | 1.59 | N | hypothetical protein | hypothetical protein |
| 2019 | N | Y | 738079 | MBOVPG45_RS03195 | 2175 | 26 | 1.20 | 2019 | hypothetical protein | hypothetical protein |
| 2019 | N | Y | 740336 | MBOVPG45_RS03200 | 588 | 4 | 0.68 | N | hypothetical protein | hypothetical protein |
| 2019 | N | Y | 744092 | MBOVPG45_RS03220 | 945 | 7 | 0.74 | N | hypothetical protein | hypothetical protein |
| 2019 | N | Y | 745757 | MBOVPG45_RS03225 | 717 | 27 | 3.77 | N | hypothetical protein | hypothetical protein |
| 2019 | N | Y | 747240 | MBOVPG45_RS03230 | 717 | 23 | 3.21 | N | hypothetical protein | hypothetical protein |
| 2019 | N | Y | 764720 | MBOVPG45_RS03315 | 1020 | 10 | 0.98 | 2019 | hypothetical protein | hypothetical protein |
| **Both** | Y | Y | 765831 | MBOVPG45_RS03320 | 1716 | 5 | 0.29 | N | hypothetical protein | hypothetical protein |
| 2019 | N | Y | 774671 | MBOVPG45_RS03350 | 1332 | 11 | 0.83 | N | hypothetical protein | hypothetical protein |
| 2019 | N | Y | 776724 | MBOVPG45_RS03360 | 1434 | 13 | 0.91 | N | hypothetical protein | hypothetical protein |
| 2019 | N | Y | 783151 | MBOVPG45_RS03380 | 1425 | 13 | 0.91 | N | hypothetical protein | hypothetical protein |
| 2019 | N | Y | 796785 | MBOVPG45_RS03420 | 1218 | 6 | 0.49 | N | hypothetical protein | hypothetical protein |
| 2019 | N | Y | 808377 | MBOVPG45_RS03475 | 1470 | 26 | 1.77 | 2019 | hypothetical protein | hypothetical protein |
| 2019 | N | Y | 831335 | MBOVPG45_RS03515 | 645 | 3 | 0.47 | N | hypothetical protein | hypothetical protein |
| 2019 | N | Y | 835897 | MBOVPG45_RS03540 | 891 | 6 | 0.67 | 2019 | hypothetical protein | hypothetical protein |
| 2019 | N | Y | 854021 | MBOVPG45_RS03605 | 471 | 4 | 0.85 | N | hypothetical protein | hypothetical protein |
| 2019 | N | Y | 861982 | MBOVPG45_RS03650 | 696 | 12 | 1.72 | 2019 | hypothetical protein | hypothetical protein |
| **Both** | Y | Y | 865893 | MBOVPG45_RS04450 | 372 | 3 | 0.81 | Both | hypothetical protein | hypothetical protein |
| 2019 | N | Y | 871183 | MBOVPG45_RS03700 | 1854 | 30 | 1.62 | N | hypothetical protein | hypothetical protein |
| 2019 | N | Y | 881806 | MBOVPG45_RS03740 | 906 | 5 | 0.55 | N | hypothetical protein | hypothetical protein |
| 2019 | N | Y | 911465 | MBOVPG45_RS03900 | 666 | 4 | 0.60 | N | hypothetical protein | hypothetical protein |
| 2019 | N | Y | 914797 | MBOVPG45_RS03915 | 753 | 10 | 1.33 | N | hypothetical protein | hypothetical protein |
| 2019 | N | Y | 915811 | MBOVPG45_RS03925 | 534 | 3 | 0.56 | N | hypothetical protein | hypothetical protein |
| **Both** | Y | Y | 918202 | MBOVPG45_RS03940 | 657 | 3 | 0.46 | Both | hypothetical protein | hypothetical protein |
| **Both** | Y | Y | 934781 | MBOVPG45_RS04465 | 468 | 8 | 1.71 | 2019 | hypothetical protein | hypothetical protein |
| **Both** | Y | Y | 937718 | MBOVPG45_RS04470 | 1113 | 14 | 1.26 | 1982 | hypothetical protein | hypothetical protein |
| 2019 | N | Y | 943606 | MBOVPG45_RS04070 | 225 | 5 | 2.22 | 2019 | hypothetical protein | hypothetical protein |
| 2019 | N | Y | 955374 | MBOVPG45_RS04125 | 309 | 8 | 2.59 | N | hypothetical protein | hypothetical protein |
